# Exploiting Charge State Distribution to Probe Intramolecular Interactions in Gas-Phase Phosphopeptides and Enhance Proteomics Analyses

**DOI:** 10.1101/2023.09.21.558604

**Authors:** Vladimir Gorshkov, Frank Kjeldsen

**Affiliations:** Department of Biochemistry and Molecular Biology, University of Southern Denmark, DK-5230 Odense M, Denmark

## Abstract

Charging of analytes is a prerequisite for performing mass spectrometry analysis. In proteomics, electrospray ionization stands as the dominant technique for this process. Although the observation of differences in the peptide charge state distribution (CSD) is well-known among experimentalists, its analytical value remains underexplored. To investigate the utility of this dimension, we analyzed several public datasets, comprising over 250,000 peptide CSD profiles from the human proteome. We found that the dimension of the CSD demonstrates high reproducibility across multiple laboratories, mass analyzers, and extensive time intervals. The general observation was that the CSD enabled effective partitioning of the peptide property space, resulting in enhanced discrimination between sequence or constitutional peptide isomers. Next, by evaluating the CSD values of phosphorylated peptides, we were able to differentiate between phosphopeptides that indicate the formation of intramolecular structures in the gas phase and those that do not. The reproducibility of the CSD values (mean cosine similarity above 0.97 for most of the experiments), qualified CSD data suitable to train a deep learning model capable of accurately predicting CSD values (mean cosine similarity – 0.98). When we applied the CSD dimension to MS1- and MS2-based proteomics experiments, we consistently observed around a 5% increase in protein and peptide identification rate. Even though the CSD dimension is not as effective a discriminator as the widely-used retention time dimension, it still holds potential for application in direct infusion proteomics.

---

Electrospray ionization (ESI) is by far the most dominant ionization technique used for biomolecule ionization in mass spectrometry^1^, including the extensive field of proteomics. ESI is frequently coupled with liquid chromatography-tandem mass spectrometry (LC-MS/MS) and offers efficient ionization of a broad range of biomolecules.

Under ESI conditions, peptides can adopt several protonation states. Some of these states are more likely and thus more abundant, leading to the formation of a peptide charge state distribution (CSD). Currently, proteomics almost exclusively uses positive ion mode, i.e., ionization resulting in positively charged ions. Peptide cations are preferentially produced by protonation of the most basic sites in peptides. The gas-phase basicity (GB) of these sites is determined by various factors, including the intrinsic GB of amino acids, although it is not the sole decisive parameter. Other factors also play substantial roles. For instance, the presence of functional groups capable of sharing protons, known as charge solvation groups, can increase the energy required to remove a proton from a charged site, effectively enhancing the GB of that site. Additionally, the Coulomb repulsion between neighboring protons already present reduces the proton affinity of a potential protonation site. This reduction in GB can make a residue that is inherently less basic but more distant from existing protons more favorable for protonation. Combined, these factors influence the pattern of protonation which has been measured experimentally^2,3^ and has been a subject of investigation since the beginning of tandem mass spectrometry^4,5^. The peptide CSD is also influenced by the peptide conformational space, which encompasses the array of possible conformations arising from different combinations of bond rotations, dihedral angles, and side chain orientations. In that way, different peptide structures can affect the accessibility of specific amino acid residues for protonation or deprotonation, thereby impacting the observed charge states. Finally, some polypeptides can form salt bridges in the gas phase, which are electrostatic interactions between amino acid residues of opposing charge and have been widely studied due to their structural and functional importance in proteins and peptides^6–9^. In the context of CSD, the formation of a salt bridge can have the potential to affect this parameter since such interaction involves deprotonation of the phosphate motif. Earlier important work has noted measurable compaction of specific phosphopeptides in ion mobility mass spectrometry (IM-MS)^10–12^. In this relation, Clemmer and co-workers^13^ determined lower intrinsic size parameters for the phosphorylation motif compared to any other amino acid. The intrinsic size parameter is a measure of the average value of each amino acid residue or modification that contributes to a peptide’s cross-section. It was suggested that the presence of the phosphoryl group can play a role in intramolecular interactions that cause an overall compacting effect on the peptide structure compared to sequences without this modification.

Peptide CSDs and their relation to peptide sequence have been the focus of two recent studies by Guan et al.^14^ and Xu et al.^15^. The study of Guan et al. demonstrated the possibility of accurate prediction of the CSD using a deep learning long short-term memory (LSTM) model. The study of Xu et al. aims to uncover sequence features responsible for peptide charging by employing a more explainable logistic regression model based on peptide amino acid composition (i.e., not accounting for amino acid interactions). Both studies have indicated the potential utility of their models and findings to advance peptide identification, though definitive outcomes have yet to be presented. Additionally, neither study has considered phosphorylated peptides – a prevalent and extensively researched category of post-translationally modified peptides.

Building on these foundational studies, we pursue here to explore the broader analytical importance of CSD in proteomics and investigate in particular the CSD landscape of phosphorylated peptides. In doing so, we examined multiple public datasets, encompassing over 250,000 peptide/phosphopeptide CSD profiles from the human proteome. We emphasize CSD consistency, its likely capacity to distinguish phosphorylated peptides with gas-phase intermolecular structures, and its potential to enhance peptide and protein identification rates in both MS1 and MS2 proteomics experiments.

## Methods

Complete experimental details are provided in Supplementary Material.

### Software tools

Unless stated otherwise, data manipulation and plotting were performed in Python (3.10.2) with the following modules installed: ***numpy*** (1.23.5), ***scipy*** (1.10.0), ***matplotlib*** (3.5.1), ***pandas*** (1.4.0), ***scikit-learn*** (1.0.2), ***seaborn*** (0.11.2), ***pyteomics*** (4.6), ***keras*** (2.10.0), ***statsmodels*** (0.13.1), ***pygam*** (0.8.0). Relevant scripts are provided as supplementary material.

### Database search

The non-phosphorylated dataset was searched using MSGF+ (2022.01.07)^16^ against the UniProtKB human protein database supplemented with reversed decoy sequences. Search results were validated using Percolator (3.05)^17^ and PSMs were filtered to q-value < 0.01 for any further processing. Deposited MaxQuant results were used for phosphoproteomics datasets when available. Raw files belonging to the PXD000138 dataset (synthetic phosphopeptides) were reanalyzed using MaxQuant (2.0.3.0)^18^. Results were filtered to 0.01 FDR for proteins, peptides, and sites. PSMs with all phosphorylated sites having a localization score above 0.95 and Andromeda score above 50 were considered for further processing. For the synthetic phosphopeptide datasets, peptides matching the synthesis goal were considered valid and accepted for further processing.

### Charge-state distribution calculation

Raw files were converted to mzML format using ***ThermoRawFileParser*** (1.4.2)^19^ and further processed by ***biosaur2*** (0.2.11)^20^ to detect LC-MS features. Peptide-spectrum matches (PSMs) were mapped to detected LC-MS features and further grouped using retention time, charge, and mass. Mass, elution apex, start, and end retention times of feature groups were used to create extracted ion chromatograms for all charge states from 1 to 6 using ThermoRawFileParser. The area under the curve (AUC) was calculated for every XIC and assembled into a vector representing CSD. CSDs belonging to the same peptide sequence were grouped and the consensus CSD was calculated and stored. For the timsTOF data, CSDs were calculated as published earlier^15^.

### CSD prediction model

Peptide sequences were one-hot encoded, with special codes used for N-terminal acetylation, oxidated methionine, and phosphorylated serine, threonine, and tyrosine. Peptides longer than 40 elements after encoding were discarded. Deep learning model architecture was described earlier^14^. The model was compiled with an Adam optimizer and negative cosine similarity loss function.

### DirectMS1 analysis

Raw files were converted to mzML using ***ThermoRawFileParser*** (1.4.2) and, next, ***biosaur2*** (0.2.11) has been used for feature detection. The search was performed by ***ms1searchpy***^21^ against the SwissProt human protein database, concatenated with decoy sequences generated by pseudo-shuffling^22^. CSD-specific features were included in the feature table used in the machine-learning-based peptide-feature match (PFM) classification. For the reduced CSD experiment, the following features were included: the number of times a peptide sequence is observed in the PFM list; the number of unique charge states a peptide sequence is observed in; the number of unique ion mobility values a peptide sequence is observed in. For the complete CSD experiment, a CSD prediction step has been implemented in ***ms1searchpy*** (version 2.6.3). To this end m/z values corresponding to charge states from 1 to 4 were calculated for all detected PFMs; ThermoRawFileParser (1.4.2) was used to produce XICs; AUC has been calculated for each XIC and these values were assembled to experimental CSD vectors; the model trained earlier was employed to predict CSDs for every unique peptide sequence. The following properties were calculated for every PFM: spectral angle between the experimental and predicted CSD, the difference between the experiment and the prediction of each charge state, and the average observed charge all expressed as z-scores.

### MS/MS analysis with CSD

Raw files were searched using MSGF+ and Percolator and experimental CSDs were calculated as described earlier. The Percolator input file was modified to include CSD-related features for each entry in it. The developed deep learning model was employed to predict CSDs for each unique peptide sequence. Errors between the experimental and predicted values of each charge state relative abundance and observed average charge represented as z-score were used as features for the Percolator. For PSMs lacking experimental CSD, an average value of the corresponding features was used (for target and decoy populations separately). Finally, the simple matching score was calculated consisting of the absolute and relative (to peptide length) number of matched theoretical fragments. A separate input file was generated for every combination of employed features and submitted to Percolator. Another set of input files was created by shuffling randomly score-related features with and without charge features.

## Results and Discussion

### Charge-state distribution definition

Throughout this manuscript, under the term CSD, we assume the normalized (sum of all abundances equal to one) relative abundance of the peptide ions having charge states from 1 to 6 for the same peptide sequence. Thus, the CSD of a peptide is a vector with six dimensions. Furthermore, one can use this vector to calculate the average observed charge state as a scalar product of CSD and vector of charge states (*CSD · [1 2 3 4 5 6]*). Details of CSD calculation can be found in the Extended Methods section. Noteworthy, in this study, we deal solely with experimentally observed peptide abundance, that can be influenced by, for example, 1) difference in ion transmission with respect to charge and *m/z*; 2) ion image current (used for detection in Orbitrap mass analyzer) dependency on ion charge; 3) larger propensity of highly charged peptide precursors for in-source fragmentation. We do not account specifically for such effects and, thus, consider them internal for the charge-state distribution property.

### Charge-state distribution is a universal property of peptides

To assess the reproducibility of the peptide CSD feature, we have analyzed several published DDA experiments (see dataset description in Supporting Information) and extracted CSDs for all identified peptides. Focusing first on peptides without ubiquitous modifications allows us to test our analytical approach on a simpler (more standardized) case. The analyzed datasets contain HeLa digests analyzed at five different research laboratories over five years, employing several instrument models, resolutions, AGC targets, and sample load amounts. Supplementary Figure S1 represents the average cosine similarity between CSDs of the same peptides for all analyzed experiments and Supplementary Figure S2 shows concrete examples covering several similarity ranges. The majority of the LC-MS experiments display a striking level of similarity (average cosine similarity of 0.97 and higher) independently of the employed instrument, peptide sample load, gradient length, and other parameters. Especially, the independence from the gradient length and peptide sample load is quite fascinating. One could expect that short gradients will force more peptides to be eluted simultaneously and, thus, increase possible peptide ion suppression and distortion of the CSD during the ionization process. Larger peptide sample loads should have a similar effect. The data, however, clearly contradicts these expectations (compare CPR_QEX datasets for gradient length influence) and IMP_QEX (25 ng on column) with other LC-MS experiments (200 – 1000 ng on column). Some earlier reports^15^ demonstrated changes in the CSD of certain peptides resulting from different gradient lengths. However, our data clearly show that such an observation is rather uncommon. Despite these vast differences in experimental conditions the smallest cosine similarity is 0.92 and observed only for two combinations of experimental sets. It should be noted, however, that all presented LC-MS experiments have been acquired using nano-HPLC setups employing water and acetonitrile with formic acid as chromatographic buffers on Orbitrap mass analyzer, thus, a possibility exists that a different MS instrument (i.e. mass analyzer or ion interface) or a different LC conditions (buffer composition, flow rate) will influence the CSD. The potential impact of different flow rates and solvent composition was investigated by analyzing the public data from a microflow LC system with DMSO added in the buffers (Supplementary Figure S3) – the mean cosine similarity decreased to 0.98 – 0.86. To test the influence of the mass analyzer we have employed the CSD extraction routine published earlier^15^ using a dataset acquired on a timsTOFPro instrument coupled with a nano-HPLC system and compared it to our Orbitrap-based dataset. The mean cosine similarity (Supplementary Figure S4) was in the range of 0.99 – 0.88. Combined these observations suggest that the CSD (given comparable LC-MS conditions) is an intrinsic property of the peptide and, thus, could be potentially employed for peptide characterization.

### Exploring the charge-state distribution properties and discriminatory power

To further explore the properties of the CSD dimension we reanalyzed a portion of a recently published dataset (PXD024364)^23^ containing K562 cell lysates digested by trypsin and LysC and fragmented with higher-energy collisional dissociation (HCD). We extracted in total 538,124 CSDs from both trypsin and LysC datasets, these resulted in 234,175 unique peptide CSD features. Supplementary Figures S5 a,b show the distribution of the cosine similarity between the CSDs and the standard deviation of the average calculated charge for the same peptides (for peptides detected more than once). Both figures indicate high reproducibility of the detected CSDs, for example, over 88% of peptides have a deviation of the calculated average charge below 0.1. Supplementary Table 1 summarizes the number of different observed charge states (i.e., the number of non-zero elements in the CSD vector) for detected peptides. Most peptides are detected in two or three different charge states, this observation aligns well with the earlier knowledge. Noteworthy, for approximately 42% of peptides analyzed, a single charge state accounts for over 90% of the total ion abundance (Supplementary Figure S6). As expected, more abundant peptides tend to have a larger number of observed charge states (Supplementary Figure S5d), although, the trend is rather weak, thus, it is likely that the number of observed charge states is primarily influenced by the structure of the peptide itself. As seen in Supplementary Figure S5c the average calculated peptide charge of tryptic peptides spans a wide range from 1 to 5.5, while most of them are detected with an average charge from around 1.75 to 4.

Since we have established that the CSDs can be reliably determined and are largely independent of the LC-MS conditions, we decided to investigate if this property is capable of distinguishing peptides. Modern MS instruments allow extremely robust and precise measurement of peptide mass, however, many peptide sequences are sequence isomers (i.e., the same amino acids in a different order), or constitutional isomers (i.e., the same composition of chemical elements that can be organized differently in the sequence, such as, for example, Q and GA). Both sequence and constitutional isomers cannot be distinguished by mass alone, since they have identical masses, and, thus, present a challenge for MS-based proteomics if the fragmentation of a peptide is incomplete or not used at all (MS/MS-free approaches). Around 6% and 52% of peptides in the K562 dataset have at least one sequence or constitutional isomer, respectively. The histogram of the cosine similarity and the average charge difference for peptides in each class is presented in Figure 1. The presented data corresponds to the minimal pairwise cosine similarity and the maximal pairwise difference in the average charge of the peptides having the same amino acid or elemental composition, thus, indicating if it is possible to differentiate at least one pair of these peptides, using the corresponding metric. The shaded area indicates the region of cosine similarity below 0.95 and the average charge difference above 0.1. These cutoffs have been selected according to data presented in Supplementary Figure S1 (cosine similarity of replicated CSD measurements) and Supplementary Figure S5b (average charge state difference of the same peptide). The CSD is only slightly beneficial for the discrimination of sequence isomers (9%), while it allows substantial differentiation of constitutional isomers (53%). The average charge state difference (Figure 1) presents a stronger discriminator in general. For sequence and constitutional isomers, 26% and 69% of peptides, respectively, could be distinguished. The second mode (with apex around 0.7) in constitutional isomers charge difference (Figure 1b) is particularly interesting. It indicates that a particular subset of constitutional isomers possesses an elemental composition that drastically changes the average charge of a peptide. An example of such a substitution can be an interchange of a WN pair to a YH pair that introduces amino acids with notably different gas-phase basicity. Supplementary Figure S7 shows complete cosine similarity distributions for sequence isomers, constitutional isomers (including the sequence ones), and 25000 randomly selected peptide pairs.

**Figure 1.**
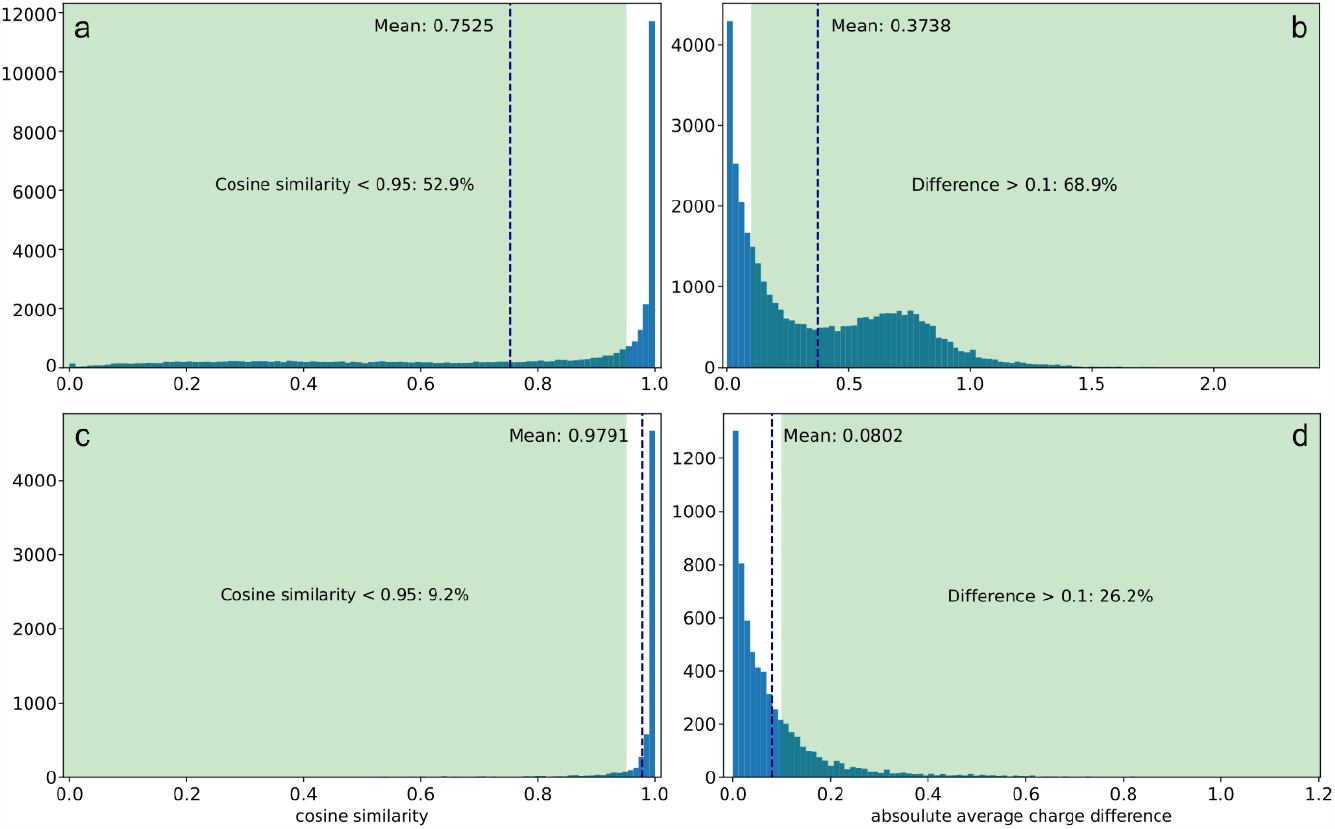
(a,b) cosine similarity and absolute difference in observed charge for strictly constitutional isomers (peptides with the same elemental composition, but different amino acid content); (c,d) the same for sequence isomers

### Charge-state distribution of phosphorylated peptides

Phosphorylation of hydroxyl-containing amino acids (serine, threonine, and tyrosine) is one of the most common and widely studied post-translational modifications (PTM) of proteins. Due to the labile nature of this PTM, precise localization of the modification is still a challenging task. Under ESI conditions, the phosphate motif can engage in a range of intramolecular interactions in the gas phase. Due to this feature, it is expected that phosphorylation (and its position) can substantially affect the CSD of a peptide. In this section, we aim to characterize this impact.

As the first step, we obtained several datasets of phosphorylated peptides and extracted reliably identified phosphopeptides having confidently localized phosphosites (please, refer to the Methods section for details). The analyzed dataset contains LC-MS/MS experiments from four different research facilities, obtained over four years (2016 – 2019) and analyzed using several Orbitrap-based instruments. Like the results obtained for non-phosphorylated peptides, we observe very high reproducibility of CSD of phosphopeptides (most cosine correlations are above 0.97, and the lowest one is around 0.935), independent of the employed instrument model, research facility, LC gradient, and other LC-MS parameters (Supplementary Figure S8).

Phosphopeptide identifications from all experiments were combined with two synthetic phosphopeptide datasets (see dataset description in Supporting Information) resulting in 531,267 CSD vectors that were merged into 42,531 unique (including phosphosite position) phosphopeptide CSDs. Supplementary Figure S9a shows the distribution of the standard deviation of the measured average charge of the same phosphopeptides over multiple measurements (only for phosphopeptides measured at least twice). The distribution is wider than for the non-phosphorylated peptides (compared to Supplementary Figure S5b) with 84% of the phosphopeptides showing a standard deviation below 0.1. Since it is reasonable to assume that the precision of the CSD extraction computational pipeline is the same as for the non-phosphorylated peptides, the wider distribution might indicate a lower data quality of the phosphopeptide data sets. The latter can stem from greater challenges in the identification of phosphorylated peptides (and specifically localization of phosphate moiety) compared to non-phosphorylated ones. The distribution of the average observed charge is however quite like the non-phosphorylated dataset, indicating that it is mostly influenced by the tryptic nature of the peptides in the dataset.

As mentioned, one of the common challenges in phosphopeptide characterization is the proper assignment of the position of the phosphoric acid moiety. It is, however, expected that phosphopeptide isoforms (phosphopeptides with the same sequence, but different positions of phosphorylation) can obtain different gas-phase structures resulting in different representations of poly-protonated ions). We could identify 1695 groups of phosphosite isomers. For the members of each of these groups, we calculated the largest observed average charge difference and smallest cosine similarity. The corresponding distributions are presented in Figure 2. While the majority of phosphopeptides demonstrate relatively minor changes in the CSD quantities when the position of the phosphorylation is changed, it is possible to differentiate at least one of the isoforms from the others in about 8.6%, and 25.7% of the cases using the cosine similarity and average observed charge, respectively. The sensitivity is comparable to the case of sequence isomer peptides (see Figure 1c-d). The same properties plotted for peptides having a different number of phosphosites are presented in Supplementary Figure S10.

**Figure 2.**
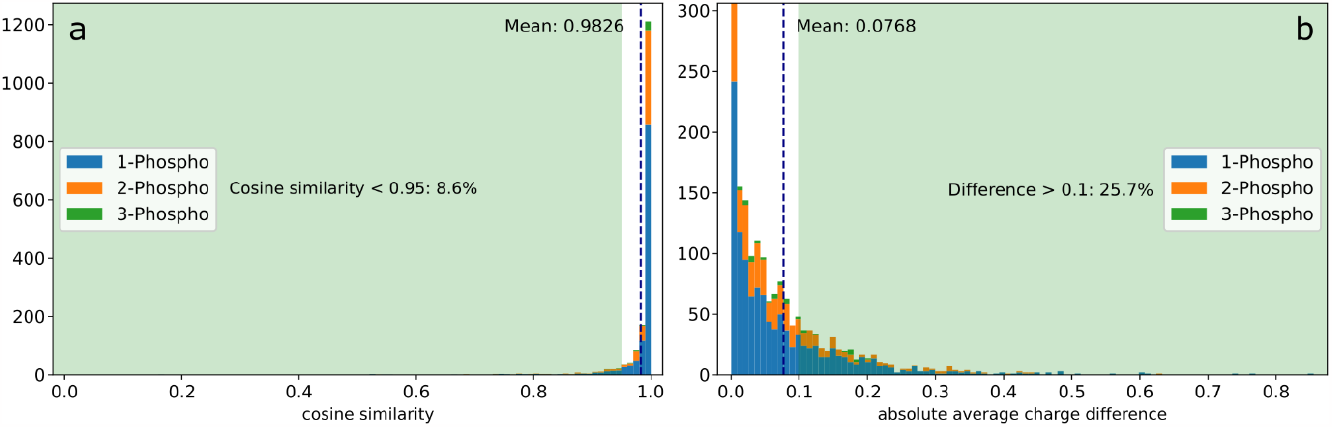
(a) Smallest cosine similarity and (b) largest absolute average observed charge difference between phosphopeptide isoforms. Only peptides with the same number of phosphosites were compared. Parts of the distribution are colored according to the number of phosphosites present, and the combined quantities are calculated for the complete distribution.

To expand the understanding of the impact of peptide phosphorylation, we investigated the changes in CSD quantity between phosphorylated peptides and their non-phosphorylated counterparts. The CSD values for the corresponding non-phosphorylated peptides were extracted from the K562 dataset described above. The pairwise differences between phosphorylated (with all possible positions and number of PTM) and non-phosphorylated peptides are presented in Figure 3a-b. Firstly, approximately 30% of peptides show a cosine similarity below the 0.95 threshold, confirming that phosphorylation substantially influences the CSD of peptides. Upon comparing the average charge state changes between phosphorylated and their corresponding non-phosphorylated counterparts, three distinct groups emerge. One, comprising 18.3% of peptides, exhibits an increase in charge (> 0.1 units) post-phosphorylation. Conversely, a second one representing 29.5% of peptides, manifests a decrease (< 0.1 units) in average charge. The remaining group does not demonstrate any substantial change in the average charge state upon phosphorylation.

**Figure 3.**
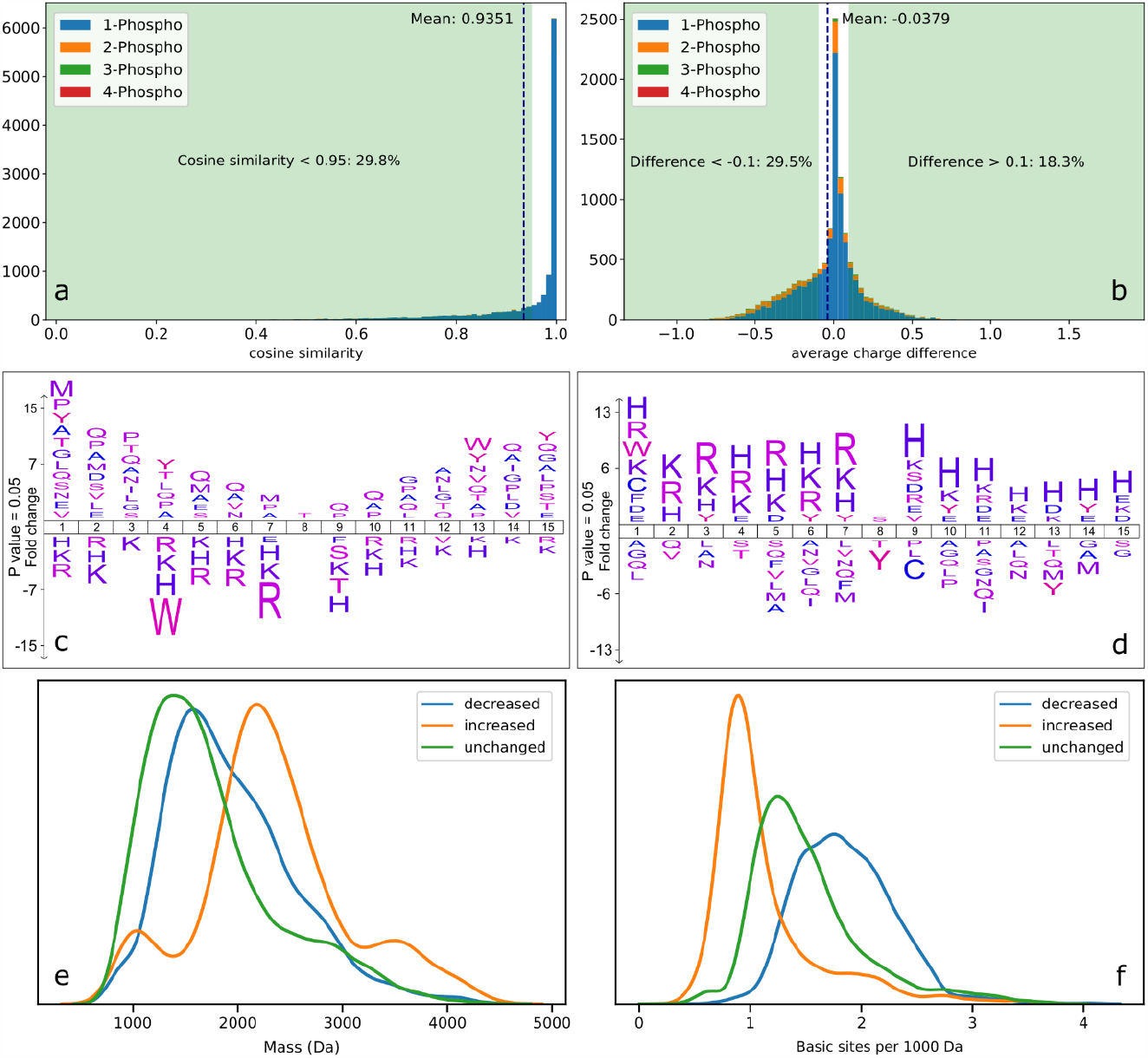
(a) cosine similarity and (b) average observed charge difference between pairs of non-phosphorylated and phosphorylated peptides with the same sequence. Parts of the distribution are colored according to the total number of phosphosites present. IceLogo (seven amino acids up- and downstream from phosphorylation site) for peptides demonstrating increase (c) and decrease (d) in average observed charge state upon phosphorylation. Distribution of peptide mass (e) and basic site density (f) for the same peptide classes. An unchanged group is provided as a reference. Only singly-phosphorylated peptides have been employed for the analysis.

To investigate this phenomenon further, we extracted sequence motifs (up to seven amino acids surrounding the phosphorylation site) for the two diverging phosphopeptide populations and analyzed the over/underrepresentation of specific amino acids (Figure 3 c,d). In this analysis, our focus was solely on singly phosphorylated peptides to eliminate potential ambiguity in the phosphosite influence. Phosphorylated peptides with a higher average charge were found to lack basic residues near the phosphorylation site, whereas those with a reduced average charge had an increased presence of basic residues in proximity to the site. We examined the size, basic site (peptide N-terminus, arginine, lysine, and histidine side chains) density (Figure 3 e,f), and several other properties (Supplementary Figure S11) for all peptide populations. Notably, the three groups differ markedly in peptide size and the density of basic residues. Compared to the group without charge state changes, the group experiencing an increase in charge state upon phosphorylation is significantly larger and possesses a lower density of basic residues. Conversely, the group with a decrease in charge upon phosphorylation is similar in size to the unaffected group but is markedly richer in basic residue density.

Recently, Xu et al^15^ introduced a concept of *under- and overcharging* that is similar to the basic site density described here. Peptides with less than three basic sites were preferably observed in the charge state above the number of basic sites (overcharging), while peptides with more than three basic sites were preferably observed in charge states below the number of basic sites (undercharging). Based on our analysis, the notion of peptides being categorized strictly into undercharged or overcharged classes seems overly simplistic. This distinction appears to stem from a fundamental relationship between the peptide’s mass and its charge-bearing capacity. The correlation of mass, average charge, and the number of basic sites align with the trend outlined by Xu et al. both for regular and phosphorylated peptides (Supplementary Figures S12 and S13).

These physicochemical characteristics and the unique responses to phosphorylation suggest distinct modes of action for each group. The population exhibiting an increased charge state post-phosphorylation has a lower initial charge, particularly relative to their larger size. The average charge increase upon phosphorylation is presumably due to the phosphate motif facilitating charge stabilization through new ionic-hydrogen interactions (between a protonated site and the neutral phosphate group). The low density of basic residues in this group suggests that additional charging is less likely if the phosphate group is involved in a salt-bridge formation. This is because, to increase the overall charge state, it would require at least two additional sites to be protonated to compensate for the deprotonation of the phosphate group. For peptides achieving favorable structural arrangements, the charge-solvating property of the phosphate group can stabilize existing charges, thereby promoting a greater portion of the ion population to occupy higher charge states in the charge state envelope. For peptides where such ionic-hydrogen structures necessitate bringing protonated sites closer together, the increase in Coulomb repulsion will inhibit this intramolecular interaction, and no gain in average charge is observed. In some cases, charge solvation and conformational changes in larger peptides can even enhance the gas-phase basicity of less basic sites in peptides, leading to further protonation^3,24^. Similarly, the phosphate group may alter the conformational space of the peptide, thereby promoting the protonation of certain ionizable sites and adding a positive shift to the distribution of charges in the gas phase.

For the group displaying a decrease in average charge state post-phosphorylation, it is suggested that this occurs due to these peptides’ high propensity for intramolecular interactions between the phosphate motif and basic sites, likely in the form of a salt bridge. These peptides are of similar size as the unaffected group but exhibit a substantially higher density of basic residues. The formation of salt bridges upon phosphorylation will effectively reduce the average charge state of the peptide unless compensation via further protonation occurs. A compensation that seems less likely since this group of peptides already has a higher initial charge. As these peptides have more protonation sites per mass, there are greater possibilities for stabilizing the phosphate motif’s negative charge with one or more positive charges. Such arrangements, involving one negative and one or two positive charges in salt bridges of gas phase peptides and proteins, have been documented in the literature^7,25–27^. Therefore, the most probable intramolecular interactions for these peptides are argued to be salt bridges, which is consistent with the observed negative shift in the average charge state for this group of peptides.

For the last group of phosphopeptides displaying no or limited change in average charge state upon phosphorylation, the interpretation lies in the notion that the intermediate basic site density of such peptides is insufficient to efficiently support salt bridge formation. Additionally, their smaller size limits the possibility of additional charging through newly formed ionic-hydrogen interactions. This is rationalized by the Coulomb repulsion within these peptides; any additional charging through potential ionic-hydrogen interactions is less likely, as the induced increase in gas-phase basicity of a site in the peptide cannot overcome the Coulomb repulsion that prohibits further protonation.

Our findings and the proposed mechanisms for changes in average charge states of peptides upon phosphorylation underscore the significant role of peptide size and basic site density in determining whether intramolecular interactions predominate. As demonstrated, a shift in charge state will occur only in a subset of the entire phosphopeptide population. This is similar to the results of IM-MS studies that attribute the negative shift in collisional cross-section (CCS) to enhanced intramolecular interactions^10^. Furthermore, recent larger-scale IM-MS studies have indicated that this ion spatial compression occurs in a substantial portion of phosphorylated peptides and is most pronounced among peptides with high charge states and extended structures^28^. While compaction in ion structures is recognized as a sign of intramolecular interactions, molecular modeling is necessary to assess the nature of these interactions – salt bridges or ionic-hydrogen bonding.

In this connection, the above observations made from the CSD data are very intriguing as they present a complementary approach to classify and characterize molecular structures of phosphorylated peptides on a large scale. The consistency and robustness of the CSD as a feature justify its use for differentiation, for instance, distinguishing phosphopeptides dominated by intramolecular interactions from those that are not. Such classification has previously been demonstrated experimentally using IM-MS^10,13,28^ and electron capture/transfer dissociation in the gas phase^25,29,30^. In addition to these established applications, it is plausible that CSD also reflects the type of intramolecular interactions involved, with salt bridge formation leading to a reduction in average charge state, while ionic-hydrogen interaction results in a gain. Such differentiation now appears achievable using CSD, but further validation is necessary to verify this potential.

A direct comparison between the CSD data and IM-MS data is intriguing when assessing potential complementarity. However, aligning them is complicated by the fact that the two methods have different scales. The CSD dimension considers the impact of phosphorylation or any other modification on the entire envelope of charge states and its conformers, while IM-MS techniques study single charge states at a time. In this context, reviewing a recent large-scale study by Ogata et al.^28^ (PXD019746) reveals that for the part of the dataset where more than one charge state for the same phosphorylated and corresponding non-phosphorylated peptides are recorded, a significant portion (>60%) of these peptides demonstrate opposite directions of CCS shifts post-phosphorylation for different charge states (Supplementary Figure 14a). This means that for the same peptide, compression is observed for one charge state while expansion is observed for another one. The latter can be the reason for a rather weak correlation between the changes in CCS and average charge (Supplementary Figure 14b).

### The potential of the CSD dimension in proteomics

For a more precise evaluation of the discriminative power of the CSDs, we categorized the data from the K562 dataset into bins based on peptide properties, including theoretical mass, retention time (apex of the elution profile), and average observed charge or CSD cosine distance. We employed various mass intervals (2, 5, and 10 ppm), retention times (0.5, 1, and 2 min), and CSD bins (0.1, 0.2, 0.4 in average charge or 0.025 and 0.05 in cosine distance). The width of the bins has been chosen based on a typical performance of the proteomics experiment, the expected width of the peptide elution profile (dataset uses 63 min gradient), and the reproducibility of CSD determined earlier. Figure 4 shows the distribution of the number of unique peptides falling into a single bin, depending on the subset of criteria used (distributions with all possible combinations of parameters are provided as supplementary material). Using the accurate mass alone with a narrow 2 ppm bin width allows having a single peptide in a bin in 28% of the cases, such a high mass accuracy is difficult to reach under routine LC-MS analysis while using more common 5 ppm and 10 ppm binning lowers the number of single peptide bins to 13% and 6%, respectively. Employing CSD information together with mass, i.e. a bin is determined in mass and CSD dimension, provides a substantial increase in the number of single peptide bins (58% and 47% single peptide bins for 5 and 10 ppm, respectively) highlighting the orthogonality of this dimension. Using retention time and mass bins dramatically improves partitioning of the peptide search space – over 95% of peptides can be separated using 2 ppm mass bins with 0.5 min retention time bin. Such a level of orthogonality between retention time and mass is well-known and, thus, the retention time dimension is currently actively employed in the state-of-the-art proteomics data analysis pipelines^31–33^. While direct infusion was among the early techniques used in proteomics^34^, it has gained renewed attention in recent research^35,36^. Our analysis shows that utilizing CSD information substantially aids in the discrimination of peptide candidates, although not to the same degree as the retention time dimension. While retention time and mass information allow superior partitioning of the peptide space, including the CSD dimension further adds small but consistent improvement in discrimination across all investigated conditions (red bars in Figure 4). The effect of charge information is (as expected) more pronounced in cases when retention time and mass cannot provide almost complete separation (for example in the 10-ppm case). Similar trends can be observed if cosine distance (0.05) is used to determine the bin size in the charge-state dimension, the discriminative power is, however, lower (Supplementary Figure S15). Our analysis suggests that CSD information is a useful addition and should allow for improved discrimination of peptides in LC-MS data analysis in general.

**Figure 4.**
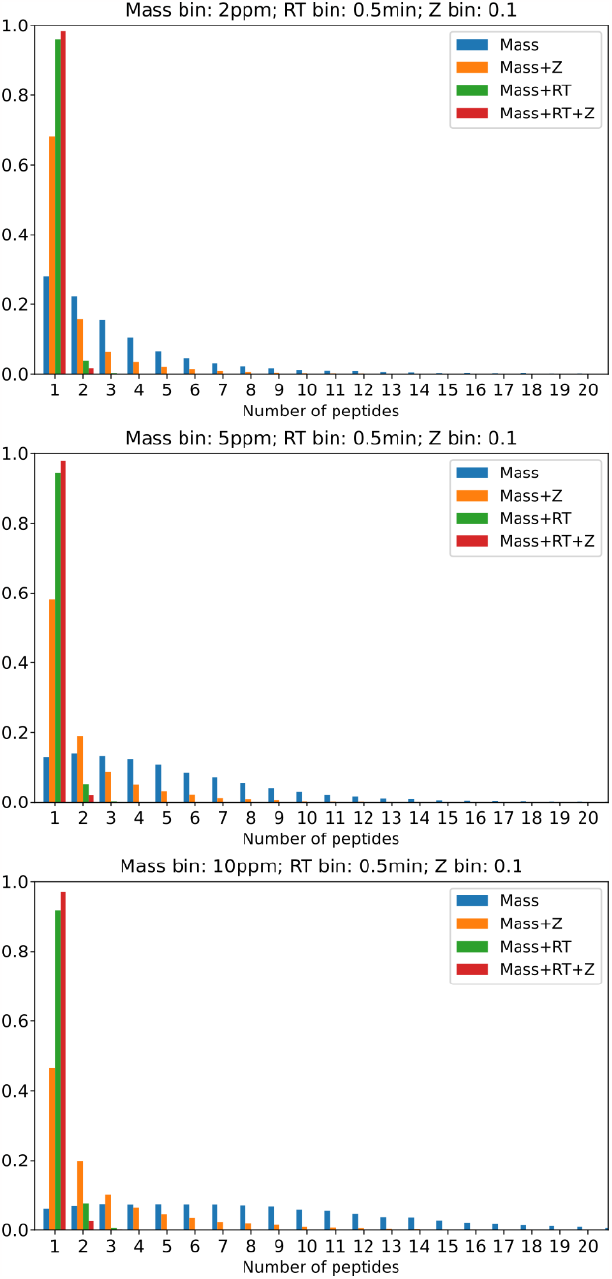
Distribution of the number of peptides in a single bin. Mass bin: top – 2ppm, middle – 5ppm, bottom – 10ppm; retention time bin 0.5min; average charge bin 0.1. Binning by mass (blue), mass and charge (orange), mass and retention time (green), mass, retention time, and charge (red).

### Using CSD for DirectMS1 and MS/MS approach

Next, we included the CSD dimension in two proteomics data analysis approaches. In this regard, we have trained deep-learning CSD predictor showing excellent performance – cosine similarity of 0.989 on the separate test set (additional details are provided in Supplementary Text 1). First, we utilized a recently developed DirectMS1 approach. DirectMS1 is a rapid proteomics screening method relying solely on the survey scan data (no fragmentation) and allowing more than 2000 proteins to be identified from 200 ng of cell lysate in approximately eight minutes per sample (180 samples per day)^21^. In this iteration, we integrate our CSD model into the DirectMS1 workflow by extending the set of peptide features used in the last machine-learning-based assessment step with the predicted CSD features. The following three approaches were compared: 1) standard DirectMS1, i.e., no CSD features added; 2) reduced CSD, only the features denoting the number of observed charge states and the number of times a particular peptide has been observed in the experiment added; 3) full CSD, features denoting the correlation between predicted and experimental CSD added. The reduced CSD approach served as additional control, to assess the importance of CSD prediction features. We have used several LC-MS experiments acquired with the DirectMS1 approach (dataset description provided in Supplementary Information) focusing on the number of proteins identified with 1% FDR for every tested experiment (Figure 5a). All experiments except the ones utilizing field asymmetric ion mobility spectrometry (FAIMS) separation demonstrate an increase (although very moderate) in the number of identified proteins. The reduction in performance by the CSD dimension in FAIMS-enabled experiments was expected, since ion mobility separation is known to separate different peptide charge states and, thus, distort CSD, making the predictions inaccurate. While the reduced CSD feature set allows for detecting more proteins, the complete CSD feature set typically excels even further in performance.

**Figure 5.**
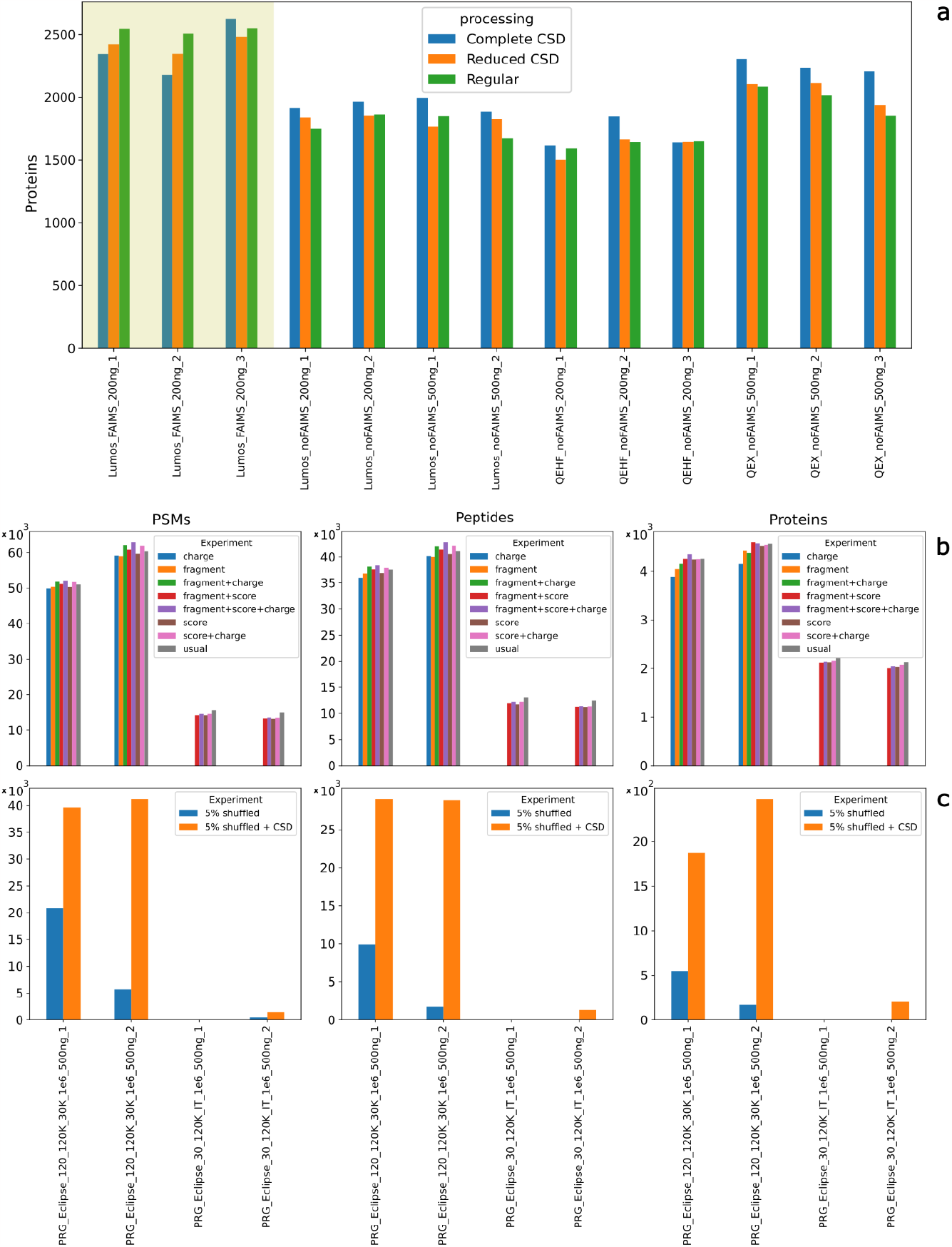
(a) The number of proteins identified at 1% FDR with the DirectMS1 approach with different feature sets used in the machine-learning step. Shaded region – LC-MS runs employing online FAIMS separation. Identified PSMs, peptides, and proteins, using different feature sets in Percolator (b), and randomly shuffling 5% of scores (c).

Finally, we integrated CSD features into the conventional MS/MS shotgun proteomics framework. Several LC-MS/MS experiments (dataset description provided in Supplementary Information) were analyzed using MSGF+ and Percolator using either a standard feature set or a few other feature sets. The default features used by Percolator were split into several classes: score – features related to MSGF+ scores; fragment – features corresponding to fragmentation, and all other features. The charge feature set contains extra features calculated from the predicted CSDs (the complete list of the features and their classification can be found in Supplementary Table 2). Lastly, we always included a simple matching score (the absolute and relative to peptide length number of matched theoretical fragments). Identification performance on PSM, peptide, and protein levels, when different sets of features were employed, are presented in Figure 5b. The observations differ for experiments employing ion trap (low resolution) and Orbitrap (high resolution) for fragmentation spectra acquisition. For low-resolution experiments using MSGF+ score features were essential for successful identification, a simplistic score even when supplemented with charge or fragmentation features doesn’t allow sufficient separation of target and decoy populations. With the Percolator input including score features identification performance returns to a level slightly below the regular one. Note that the inclusion of the charge dimension allows for a slight identification improvement. For the high-resolution experiments, on the contrary, simplistic scores in combination with charge or fragment features demonstrate a performance close to the regular feature set. Using charge features results in a more visible increase in identification while using score, fragmentation, and charge features in combination outperforms the regular approach. A similar effect of CSD features has been observed in the analysis of the phosphoproteomics dataset (Supplementary Figure S16).To further investigate the discrimination ability of charge features we randomly shuffled the score features for 5% of entries in the Percolator input and discarded all other features, the remaining 95% were preserved in the correct order, thus, diminishing their ability to serve as a good discriminator, the charge features, however, were included in the correct order (without shuffling). The results of this experiment are presented in Figure 5c. Shuffling of just 5% of the Percolator features compromises identification ability almost completely. Employing charge features allows reviving some of the performance, although even in the best cases it is markedly lower compared to the regular feature set.

The combined results of the MS/MS identification experiments demonstrate the CSD features contain valid information allowing for the discrimination of correct and incorrect peptide candidates. The discriminating power is, however, incomparable with MS/MS-based information, and thus, alone, cannot provide efficient separation of target and decoy populations.

## Conclusions

We have conducted a detailed investigation of the peptide charge state dimension to explore its analytical utility in proteome research. When comparing datasets from different laboratories, time points, and even mass analyzers, we found that CSDs are highly reproducible for peptides and phosphopeptides. This reproducibility suggests the possibility of using this characteristic to distinguish peptides. While the sensitivity of CSD to *sequence* and *PTM isomers* of a peptide is not particularly high, CSD is quite sensitive to *constitutional isomers*. Clustering peptides by their average observed charge or by the similarity of their CSD offers strong discrimination. Using mass and observed charge alone, close to 70% of peptides from a large experimental dataset can be separated, while retention time and mass allow for over 95% of peptides to be distinguished. As a result, CSDs can be an attractive dimension to utilize in direct infusion proteomics data analysis. Notably, CSD and retention time provide somewhat complementary separation; using both consistently yields deeper proteome coverage. Although the gain is moderate, it does not require additional experiments or equipment. A deep-learning predictor for CSD has been developed, demonstrating excellent performance (cosine similarity on a separate test set: 0.989) based on approximately 270,000 experimental CSDs. The trained model was tested in both MS1-only (DirectMS1) and regular MS/MS (MSGF+ and Percolator) searches, proving to enhance the discriminative power in both contexts.

By studying CSDs of phosphopeptides and their unmodified counterparts we can observe distinct populations of peptides responding differently to the phosphorylation. Foremost, distinguishing phosphopeptides that suggest intramolecular interactions from those that do not. Results indicate that these intramolecular interactions, especially the formation of salt bridges, reduce the average charge state of phosphopeptides. On the other hand, interactions involving ionic-hydrogen bonds result in a higher average charge state. The use of the CSD metric may now offer a promising method to distinguish between these interactions. However, additional research is required to confirm this capability. Potentially, this newly discovered feature of CSD can assist in the investigation of gas-phase structures of phosphorylated peptides.

Finally, we have demonstrated that the CSD dimension, which is currently underutilized in proteomics despite its readily available nature, holds the potential for improving proteomics analyses and efficiently characterizing different phosphopeptide structures.

## Supporting information

Description for used datasets

Intermediate plots

Scripts for data analysis and generation of results and intermediate plots

Extended experimental details and additional results

## Acknowledgment

This study was supported by the Lundbeck Foundation (Grant R346-2020-1215 to F.K.). Proteomics and mass spectrometry infrastructure at the University of Southern Denmark SDU were supported by generous grants to the VILLUM Center for Bioanalytical Sciences (VILLUM Foundation grant no. 7292), PRO-MS: Danish National Mass Spectrometry Platform for Functional Proteomics (grant no. 5072-00007B), and the Novo Nordisk Foundation (INTEGRA, NNF20OC0061575). The authors would like to thank Dr. Mark Ivanov for his valuable help in implementing and testing the DirectMS1-CSD approach.

## Supporting Information

The following supporting information is available along with the manuscript

- Supplementary figures and tables (PDF)
- Extended experimental details and supplementary text (PDF)
- Description of all datasets used in the manuscript (XLSX)
- Python scripts for data analysis and generating results and intermediate figures (ZIP)
- Intermediate figures (ZIP)

## Notes

### Competing Interest Statement

The authors have declared no competing interest.

### Summary of Updates

- Analysis in the "Charge-state distribution of phosphorylated peptides" section was significantly extended and improved - Minor data processing error for the K562 dataset was corrected - Comparison with timsTOF platform was added - Further smaller changes to improve readability and clarity of the presentation - Supplementary files updated

