## Supplementary figures and tables for "Exploiting Charge State Distribution to Probe Intramolecular Interactions in Gas-Phase Phosphopeptides and Enhance Proteomics Analyses"

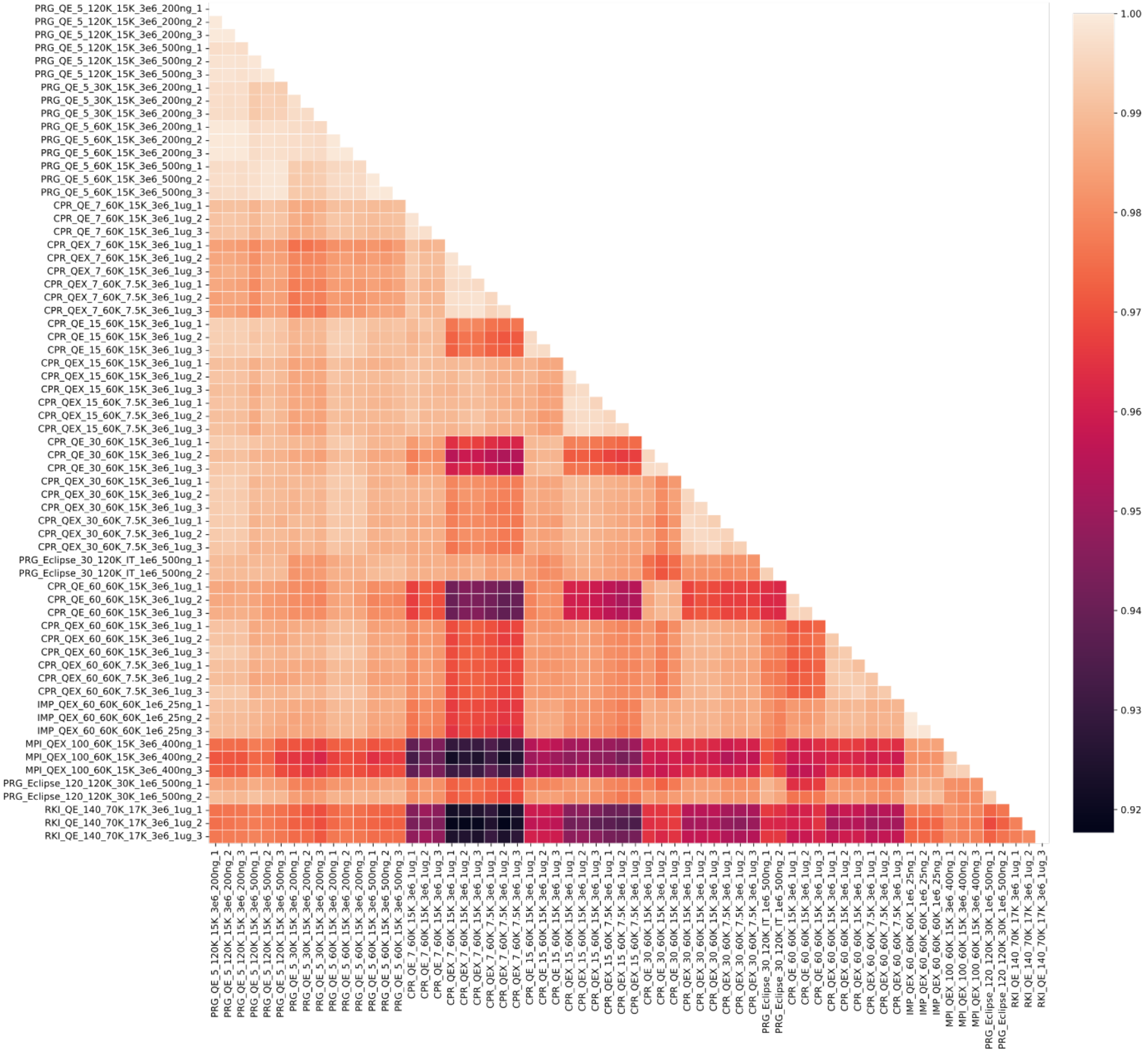


Supplementary Figure S1. Mean cosine similarity for CSDs of the same peptide in different experiments. LC-MS experiments are ordered according to gradient length; the label shows laboratory, instrument, gradient length, MS1, and MS2 resolutions, MS1 AGC target, sample load, and replicate (see supplementary table for details).

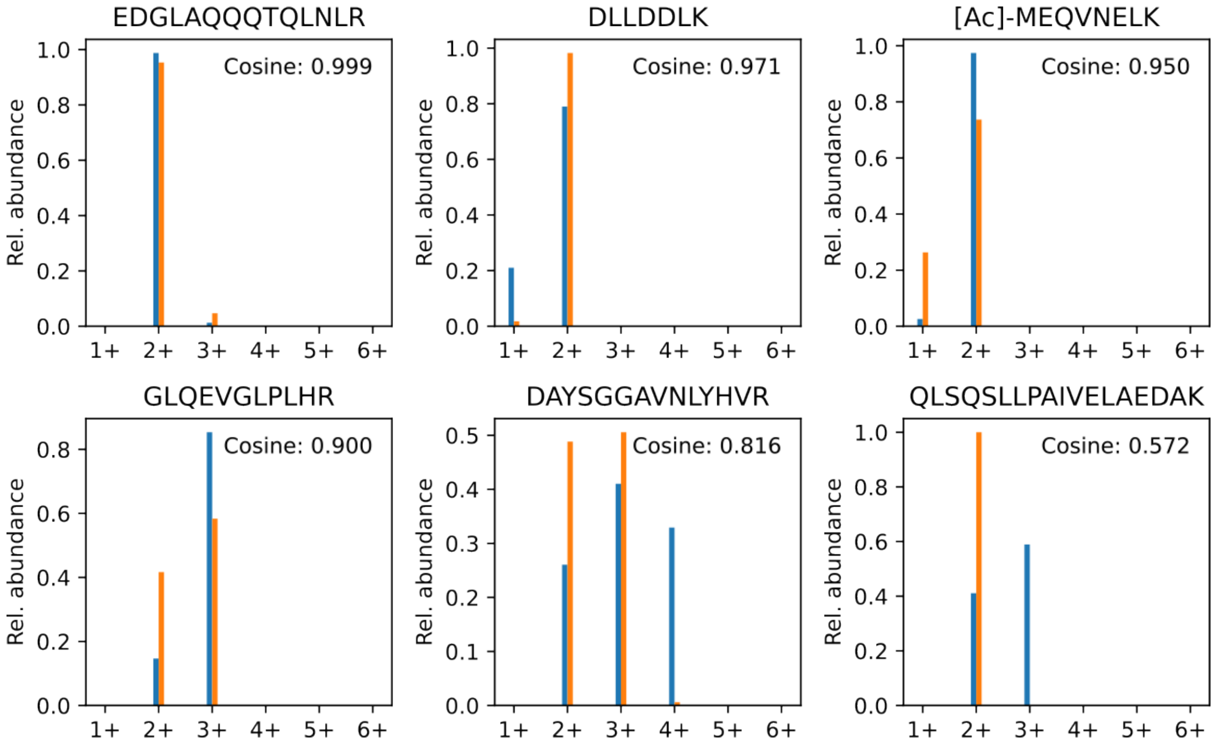


Supplementary Figure S2. Representative examples of peptide CSD similarity from several cosine similarity ranges – 0.999, 0.97, 0.95, 0.9, 0.8, and 0.5. The title shows the peptide sequence.

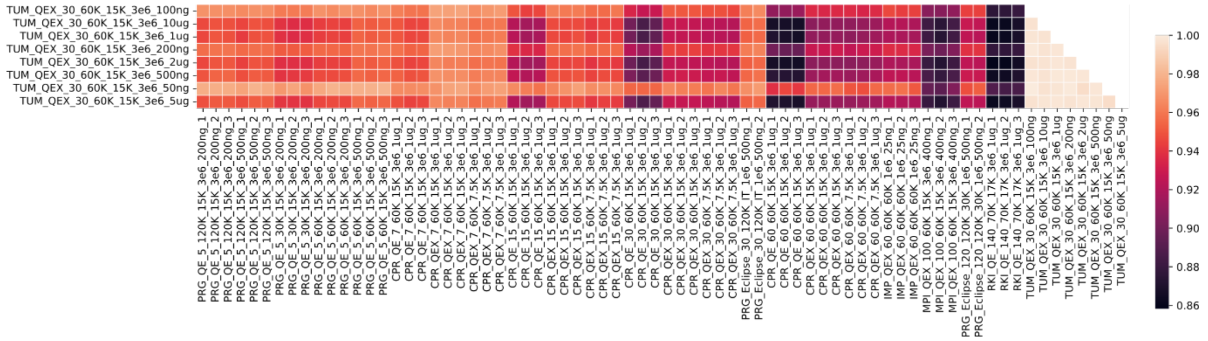


Supplementary Figure S3. Cosine similarity between CSDs in LC-MS experiments with micro- and nanoflow.

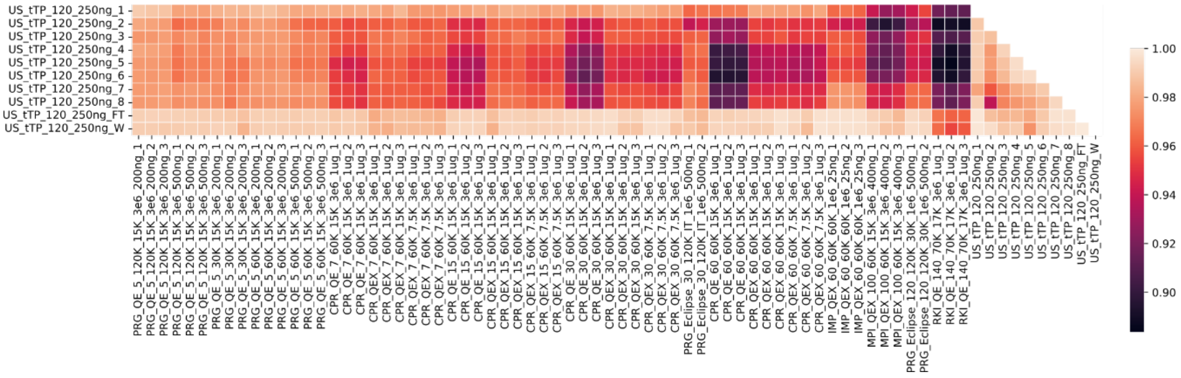


Supplementary Figure S4. Cosine similarity between CSDs in LC-MS experiments with timsTOF and Orbitrap datasets (both datasets employed nanoflow).

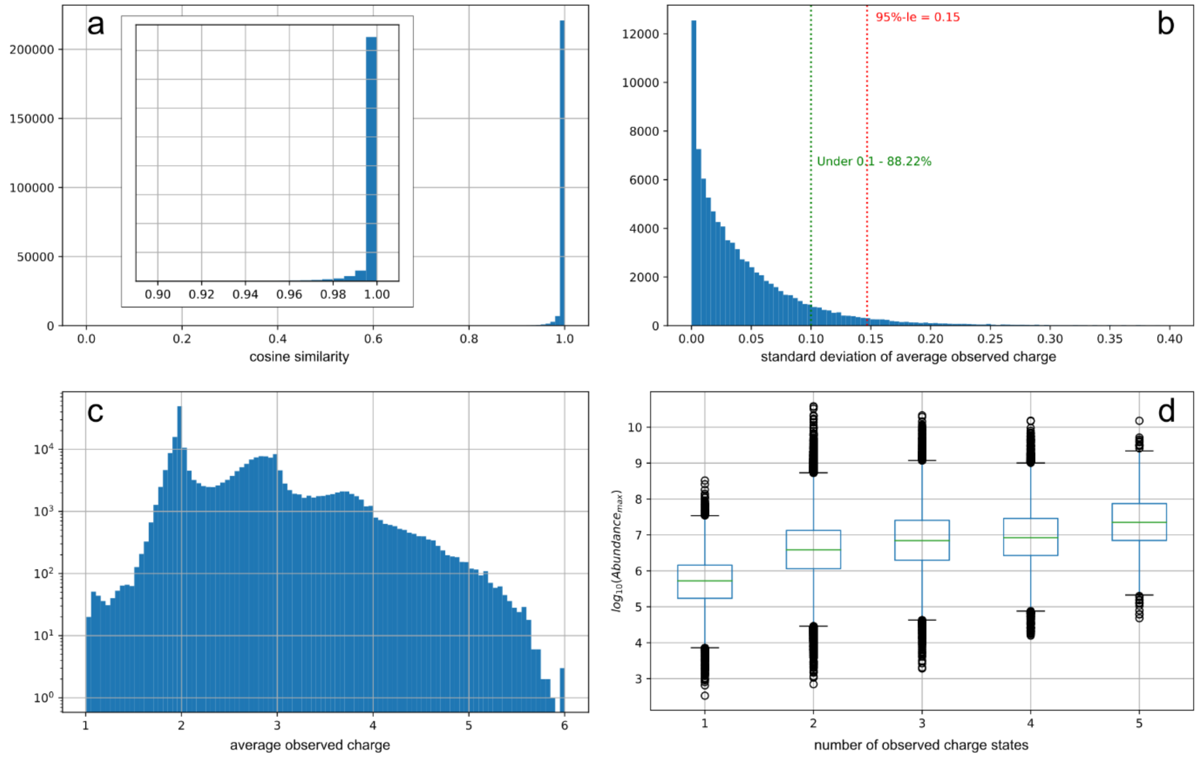


Supplementary Figure S5: (a) cosine similarity between multiple measurements of the same peptide sequence, including modifications – inset shows the cosine similarity region from 0.9 to 1.0; (b) standard deviation of average observed peptide charge for multiple measurements of the same peptide, including modifications; (c) distribution of peptides by the average observed charge (logarithmic scale of the y-axis); (d) relation between peptide observed abundance (in the most abundant charge state) and the number of observed charge states.

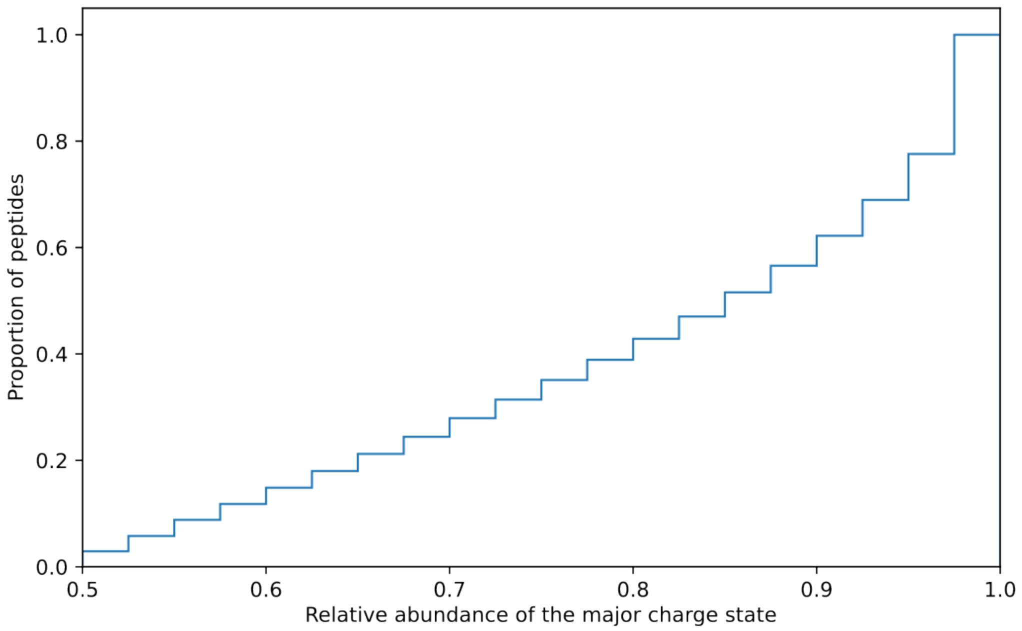


Supplementary Figure S6. Cumulative proportion of peptides according to the relative abundance of the major charge state

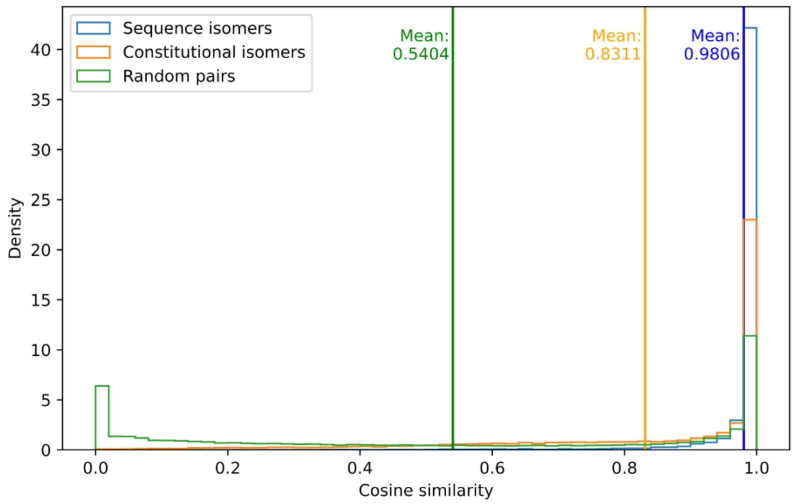


Supplementary Figure S7. Complete cosine similarity distribution between sequence isomers, constitutional isomers, and 25000 random peptide pairs. Please note, that sequence isomers are a subclass of constitutional isomers in this representation.

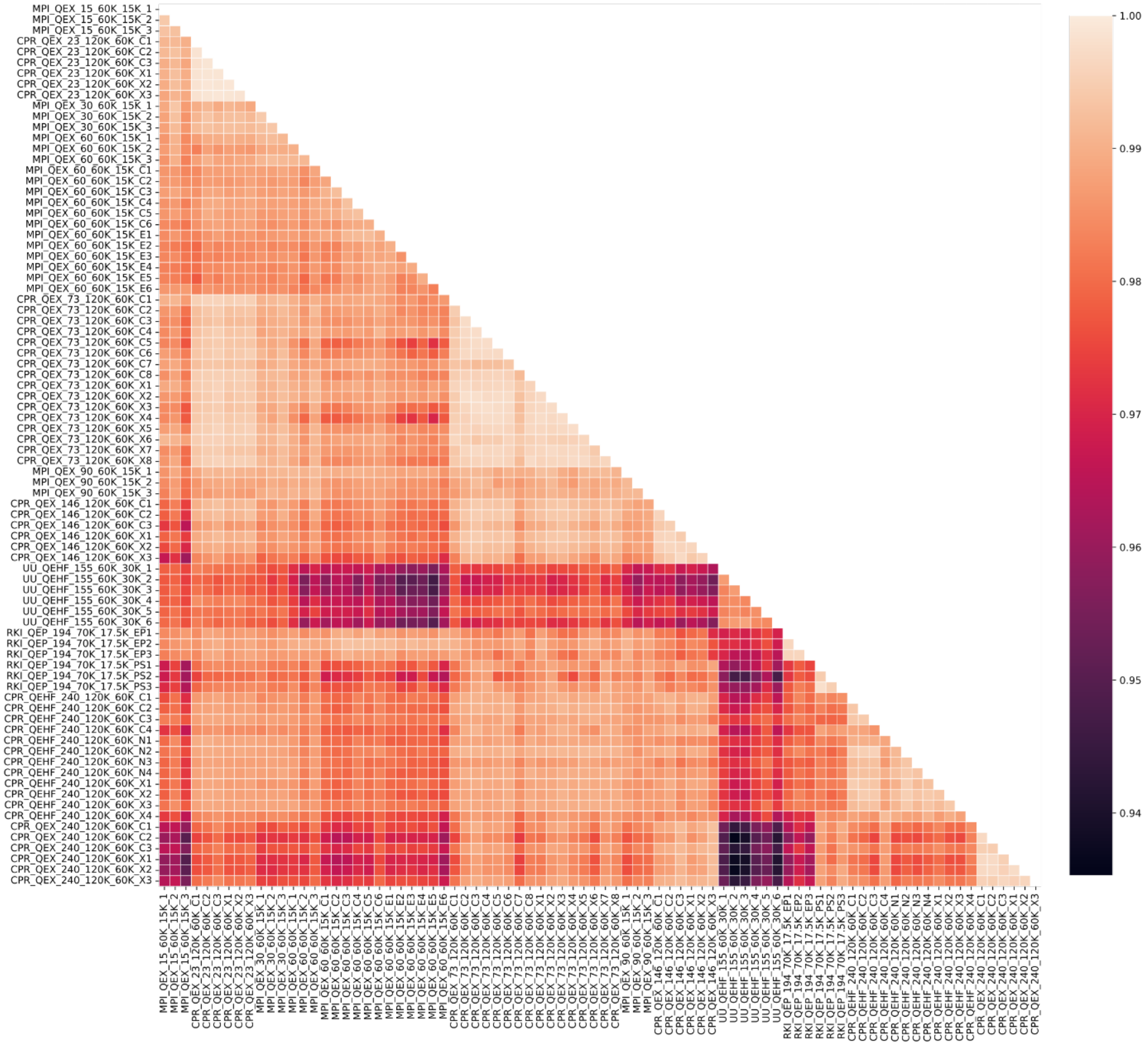


Supplementary Figure S8. Mean cosine similarity for CSDs of the same phosphopeptides in different experiments. LC-MS/MS experiments are ordered according to gradient length; the label shows laboratory, instrument, gradient length, MS1, and MS2 resolutions, and replicate (see supplementary table for details).

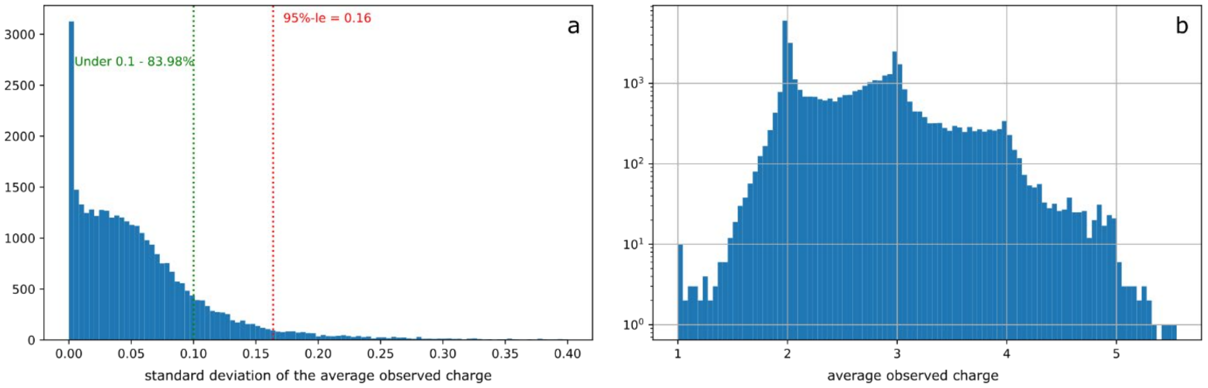


Supplementary Figure S9: (a) distribution of the standard deviation of the average observed charge for different measurements of the same phosphopeptides; (b) distribution of observed average charge of phosphopeptides

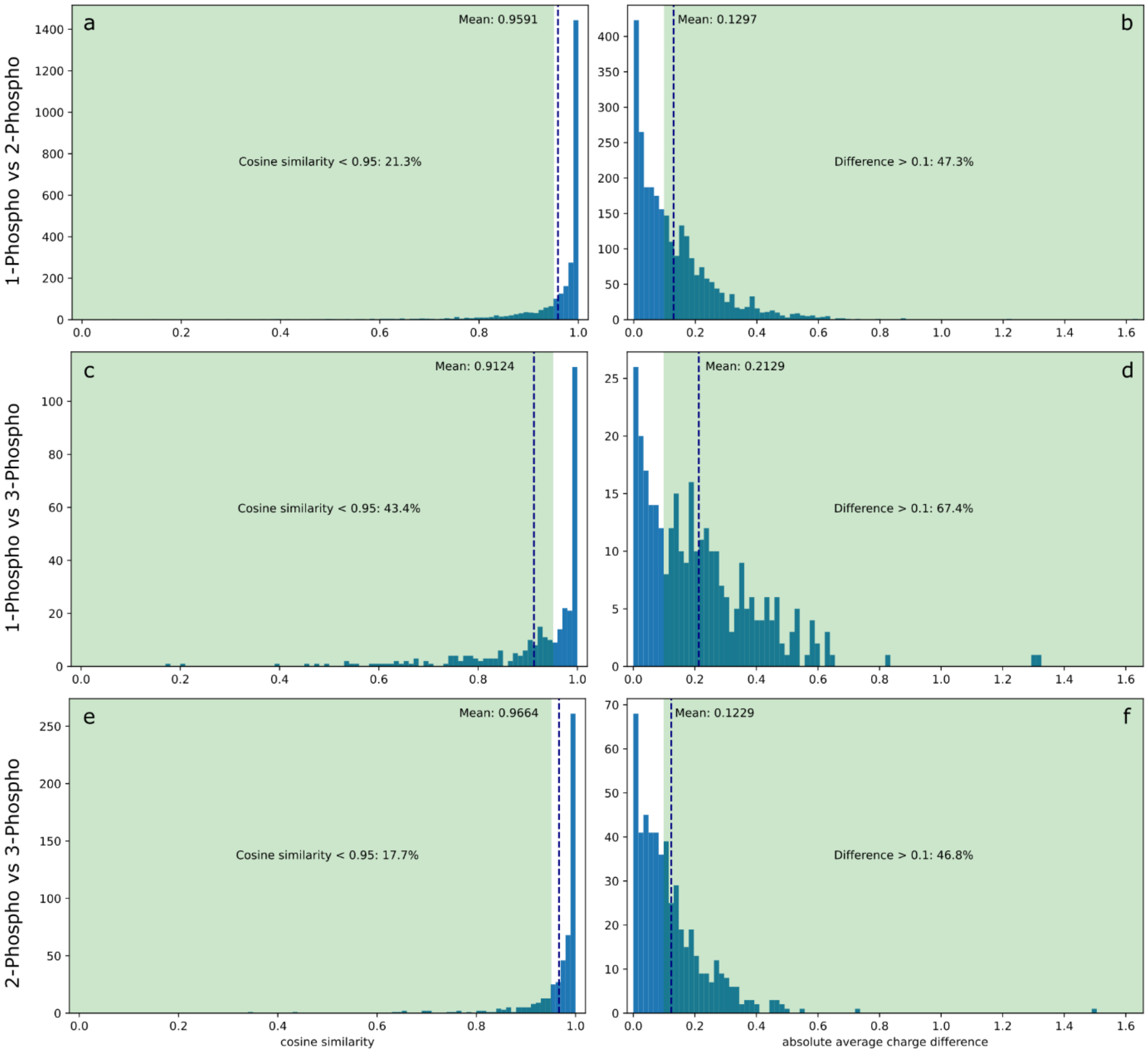


Supplementary Figure S10: Minimal cosine similarity (a, c, e) and maximal absolute average observed charge difference (b, d, f) for peptides with the same sequence except for the number of phosphorylations. Singly-phosphorylated vs doubly-phosphorylated peptides (a,b); singly-phosphorylated vs triply-phosphorylated peptides (c, d); doubly-phosphorylated vs triply-phosphorylated peptides (e, f). Other comparisons are not displayed due to the small number of cases.

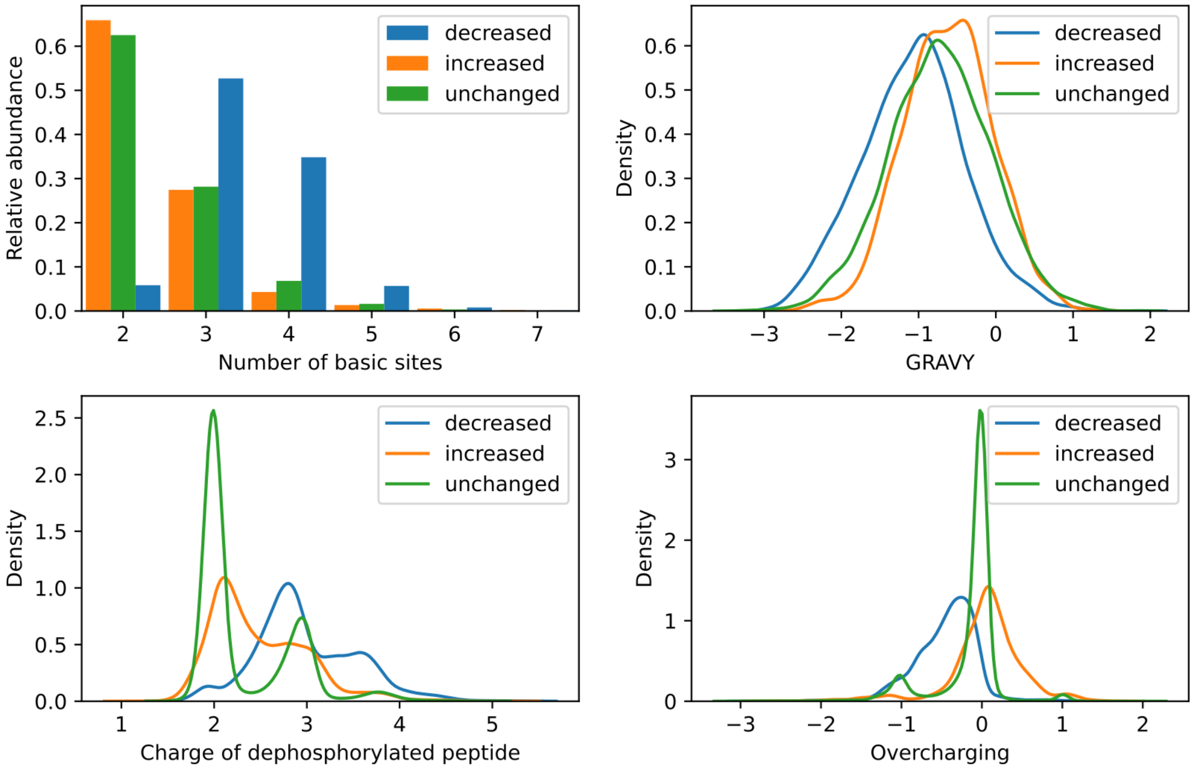


Supplementary Figure S11: Distribution of properties of phosphopeptides demonstrating increase (orange), decrease (blue), and no change (green) in average observed charge upon phosphorylation.

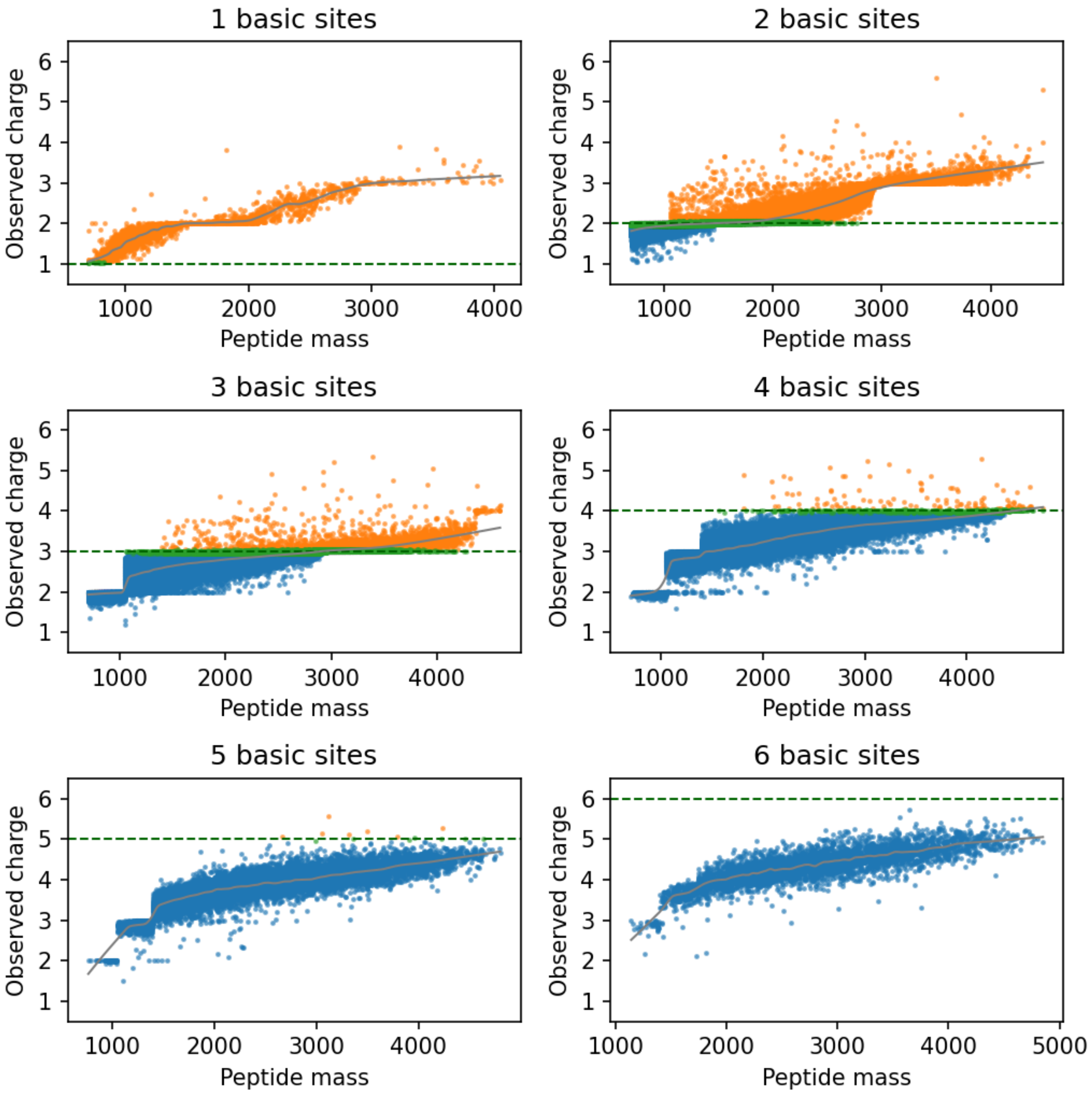


Supplementary Figure S12: Average observed charge as a function of the number of basic sites and peptide mass for the K562 dataset (only peptides with 1 – 6 basic sites are shown). Orange – overcharging (observed charge is higher than the number of basic sites), green – no charging (observed charge is close to the number of basic sites, i.e., ±0.05), blue – undercharging (observed charge is lower than the number of basic sites). The dashed green line denotes the number of basic sites, and the grey line shows the LOWESS fit of the data.

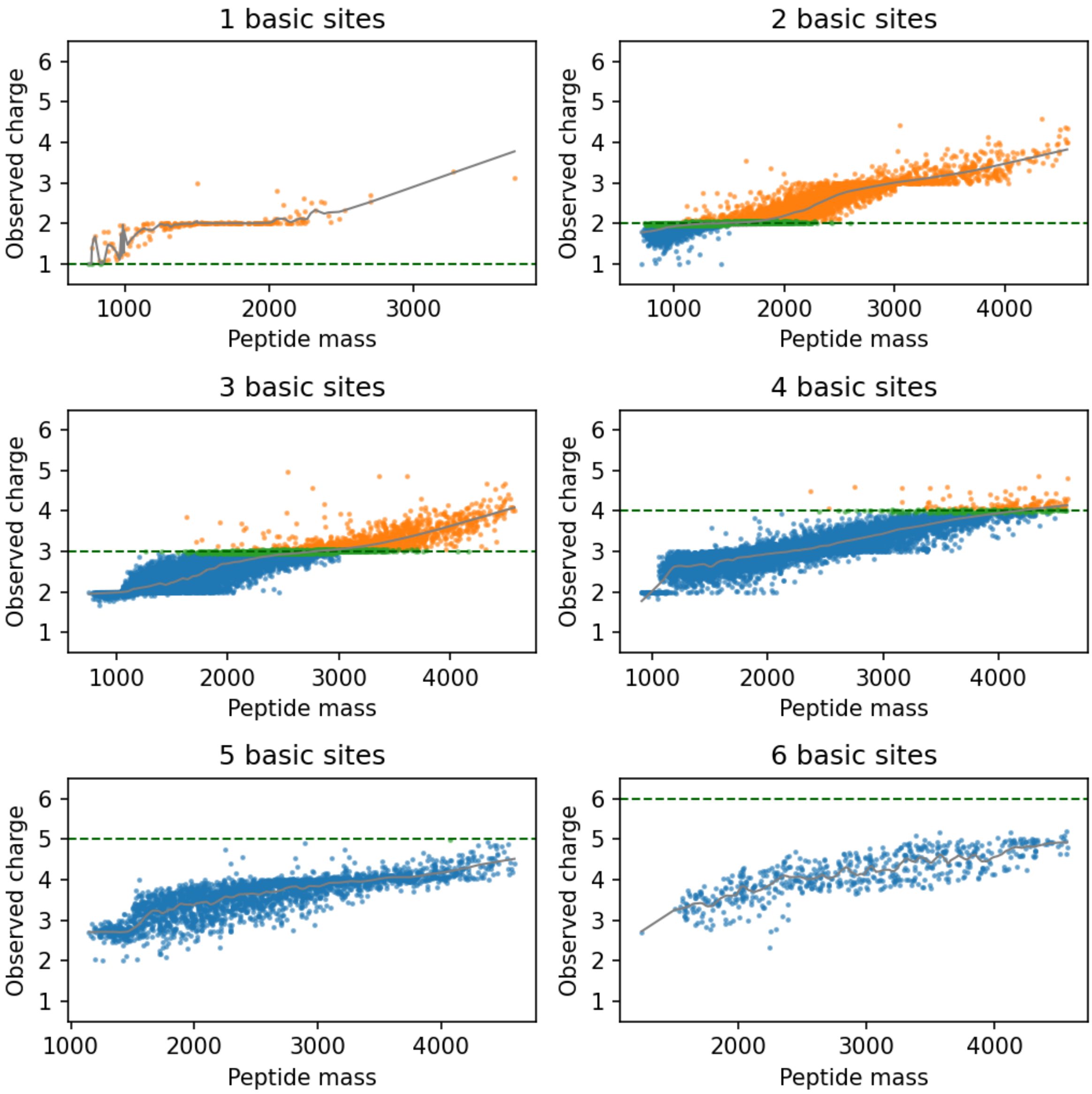


Supplementary Figure S13: Average observed charge as a function of the number of basic sites and peptide mass for the phosphopeptides dataset (only peptides with 1 – 6 basic sites are shown). Orange – overcharging (observed charge is higher than the number of basic sites), green – no charging (observed charge is close to the number of basic sites, i.e., ±0.05), blue – undercharging (observed charge is lower than the number of basic sites). The dashed green line denotes the number of basic sites, and the grey line shows the LOWESS fit of the data.

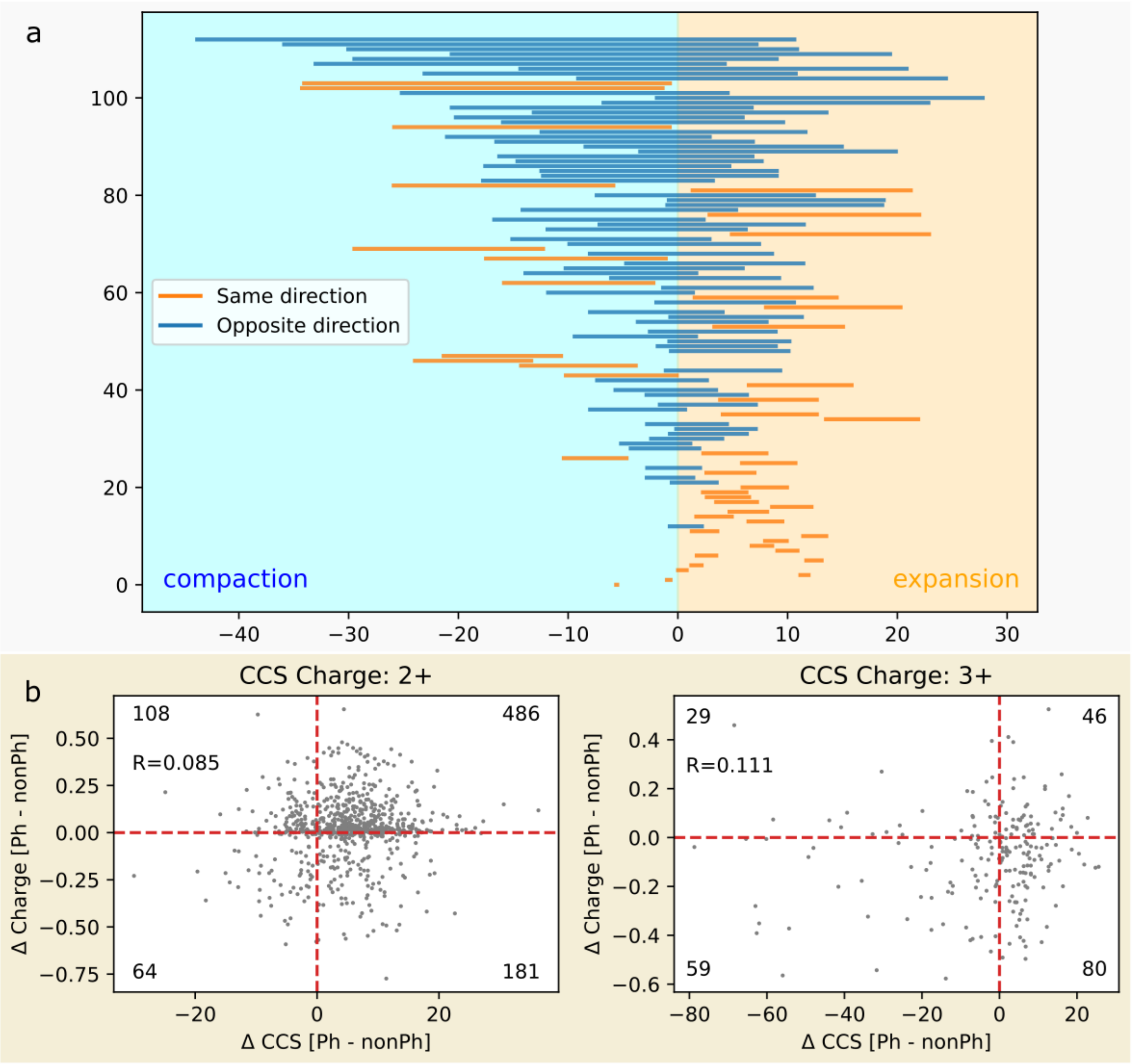


Supplementary Figure S14: (a) Size modulation upon phosphorylation, expressed as CCS change, for the same peptide in different charge states; (b) Scatter plot showing the change in CCS and average charge upon phosphorylation. The number of points in each quadrant and the Pearson correlation coefficient are shown. CCS data from Ogata et al. (PXD019746).

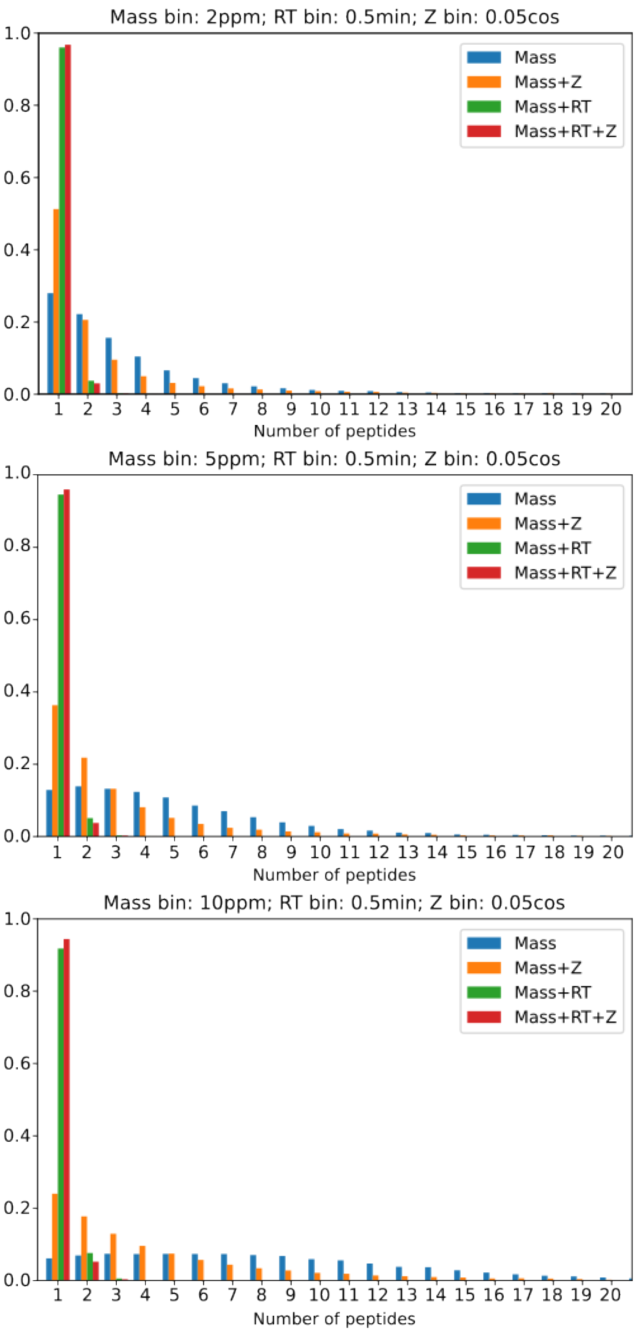


Supplementary Figure S15: Distribution of the number of peptides in a single bin in the K562 dataset. Mass bin: top – 2ppm, middle – 5ppm, bottom – 10ppm; RT bin 0.5min; charge-state bin cosine distance 0.05. Binning by mass (blue), mass and CSD (orange), mass and retention time (green), mass, retention time, and CSD (red).

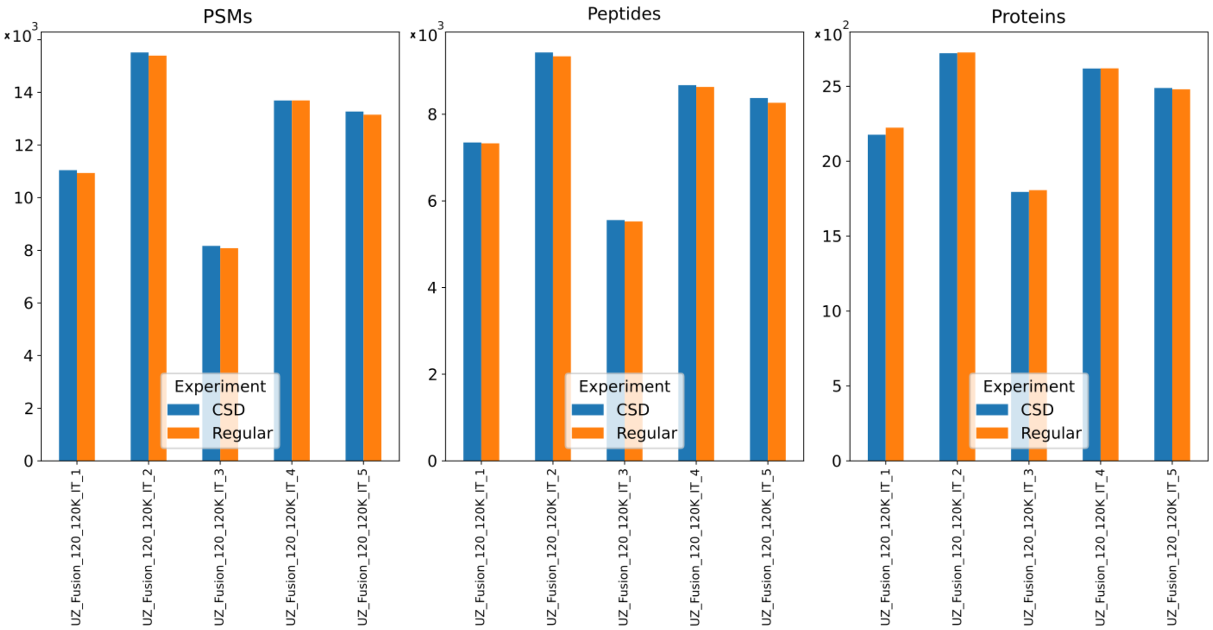


Supplementary Figure S16: PSMs, peptides, and proteins identified in the phosphoproteomics dataset with the usual workflow (MSGF+ and Percolator) and workflow employing CSD features.

### Supplementary Tables

Supplementary Table 1. Number of observed charge states for peptides in the K562 dataset

| Nr charge states | 1 | 2 | 3 | 4 | 5 | 6 |
|---|---|---|---|---|---|---|
| Nr peptides | 19268 | 136599 | 63680 | 13032 | 1596 | 0 |
| Percent | 8.23% | 58.33% | 27.19% | 5.57% | 0.68% | 0.00% |

Supplementary Table 2: Classification of Percolator features

| Percolator features | Set |
|---|---|
| RawScore, DeNovoScore, ScoreRatio, Energy, lnEValue | Score |
| MeanErrorTop7, sqMeanErrorTop7, StdevErrorTop7, lnExplainedIonCurrentRatio, lnNTermIonCurrentRatio, lnCTermIonCurrentRatio, lnMS2IonCurrent | Fragment |
| ExpMass, CalcMass | Other |
| nMatch, cover | Simple score |
| 1, 2, 3, 4, 5, 6, z | Charge |
