## Supplementary material for "Exploiting Charge State Distribution to Probe Intramolecular Interactions in Gas-Phase Phosphopeptides and Enhance Proteomics Analyses": Intermediate plots: description.txt

Folder description:

digest - plots related to complete human proteome digest
K562 - plots related to K562 dataset
ML - plots related to ML predictors comparison
phospho - plots related to phosphopeptide dataset and also comparison of phosphopeptides with their non-phosphorylated counterparts
run_correlation - heatmaps for CSD comparison accross laboratories and instruments
shotgun - plots related to employing CSD dimension in shotgun proteomics context
