## Supplementary figures and images for "Exploiting Charge State Distribution to Probe Intramolecular Interactions in Gas-Phase Phosphopeptides and Enhance Proteomics Analyses"

### decision_tree_regression.png

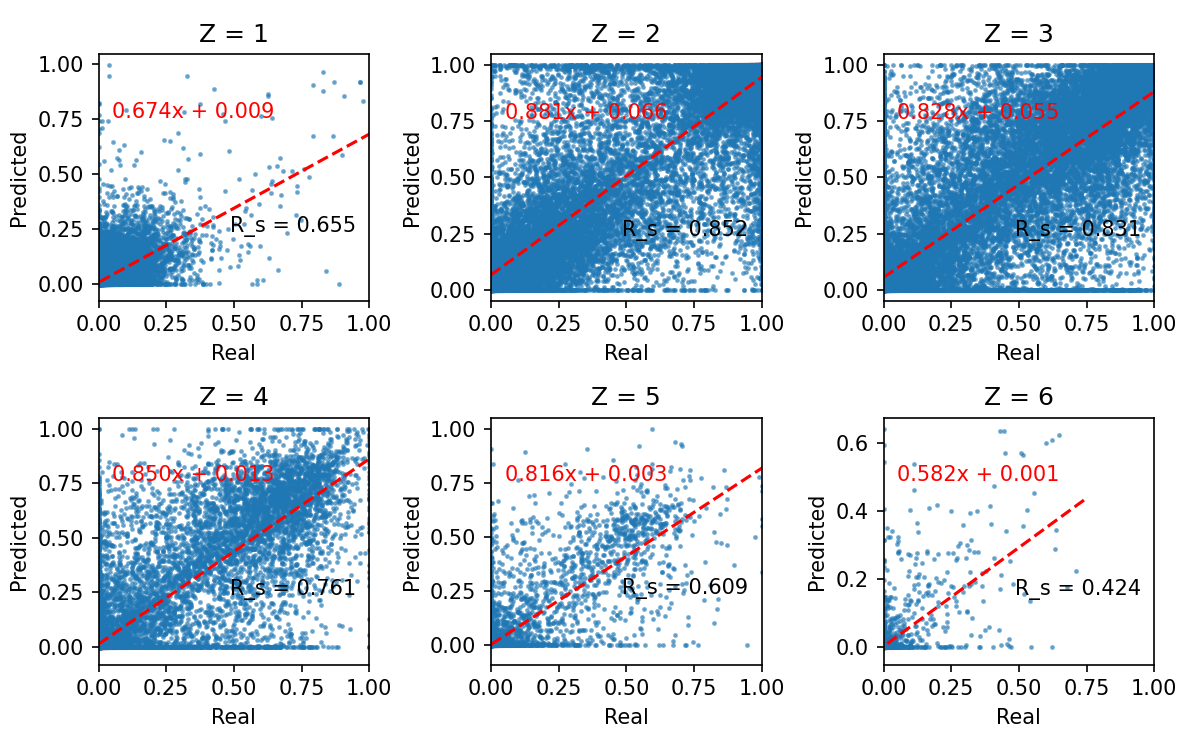

### icelogo_neg_nonneg.pdf

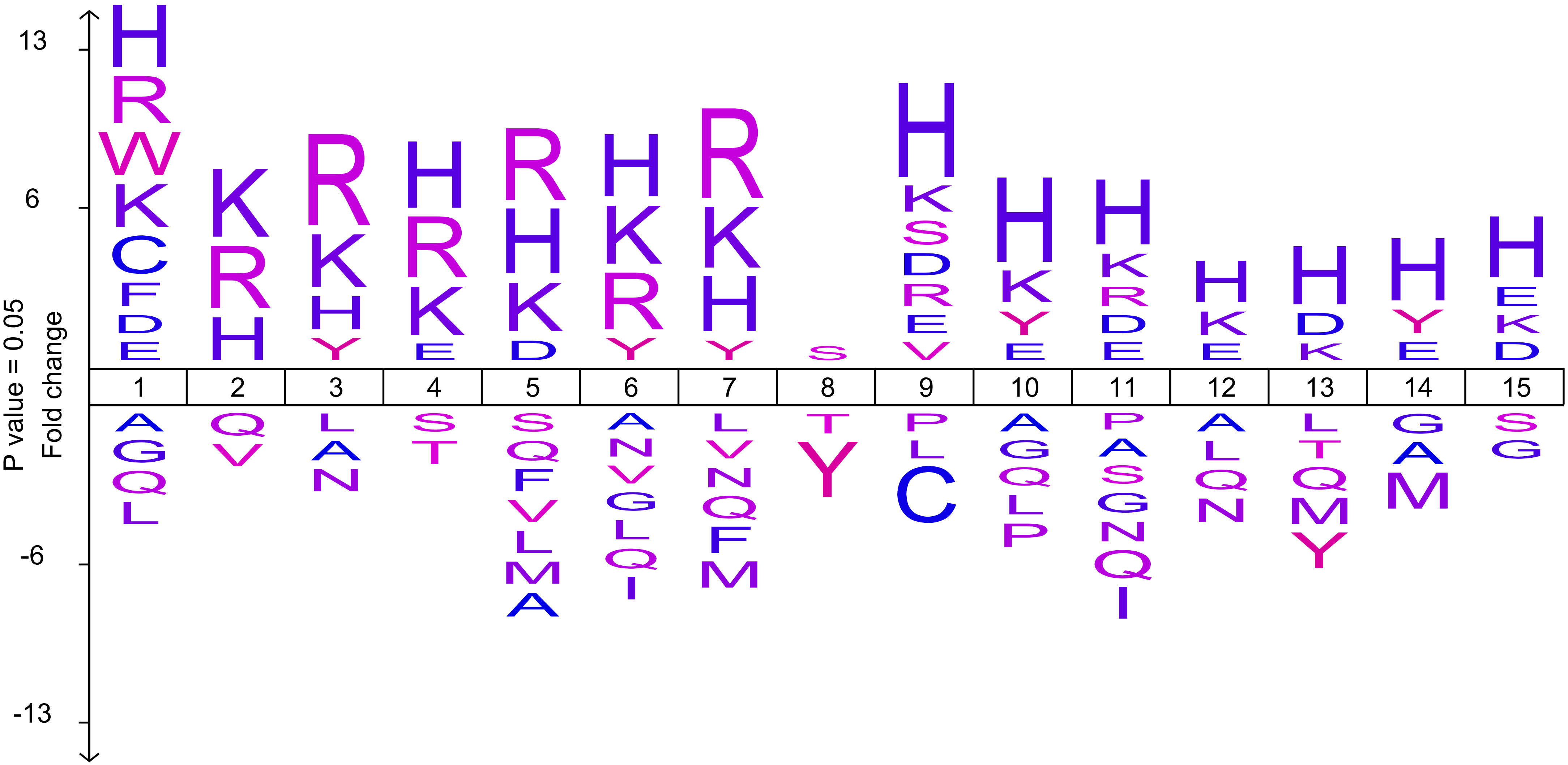

### icelogo_pos_nonpos.pdf

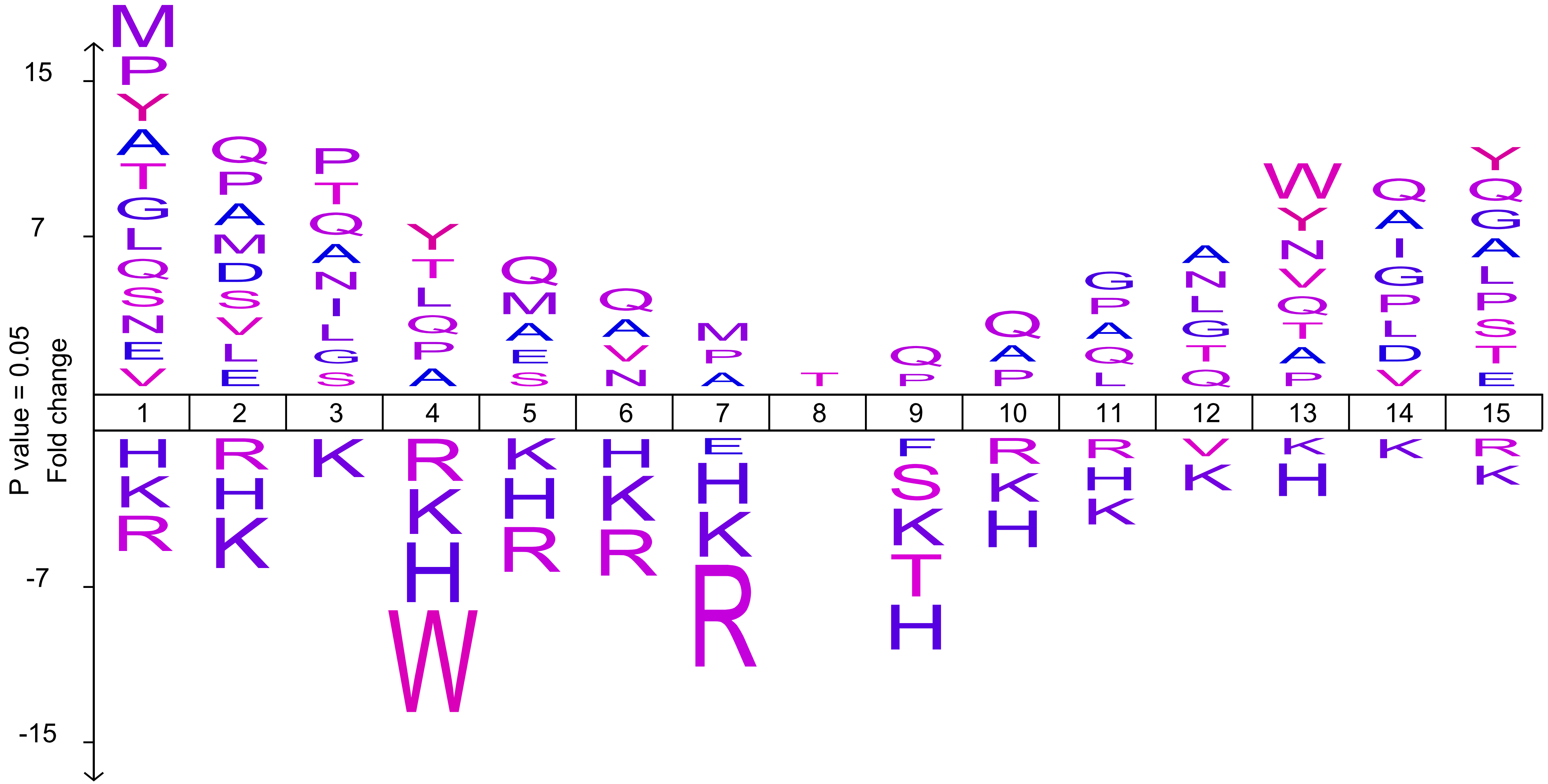

### K562_isomeric_cos_distnace_vs_mass.png

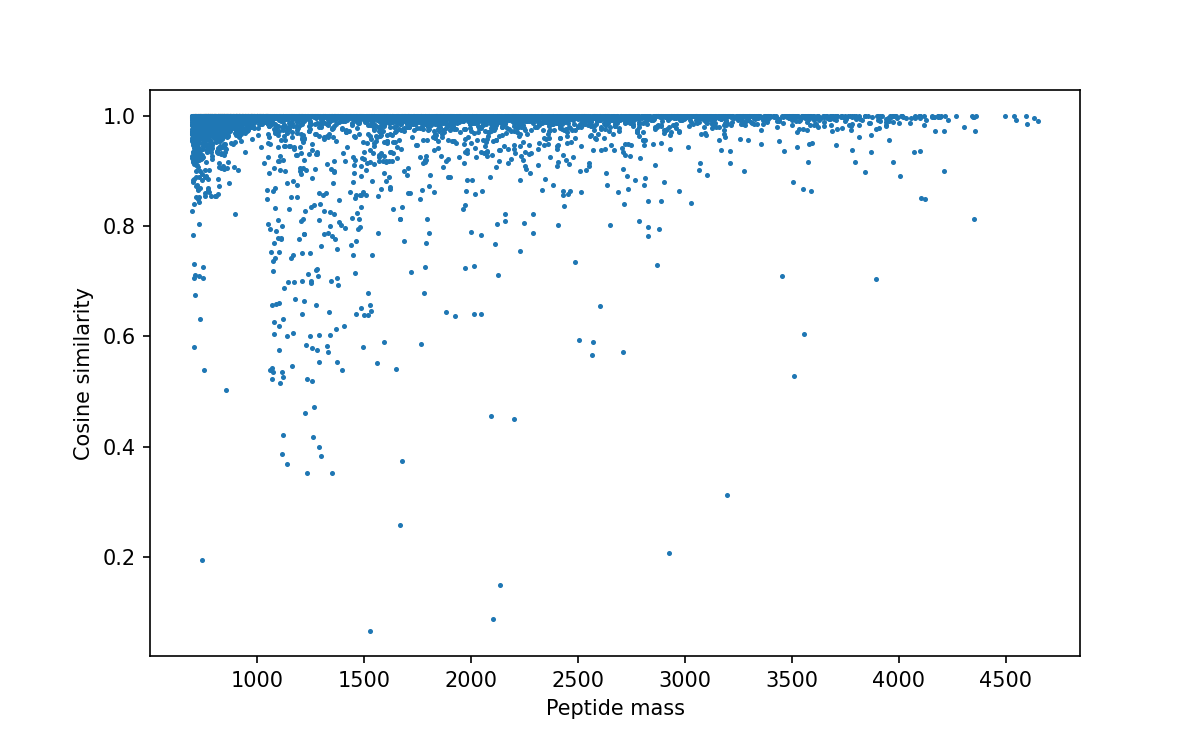

### K562_under_overcharging.png

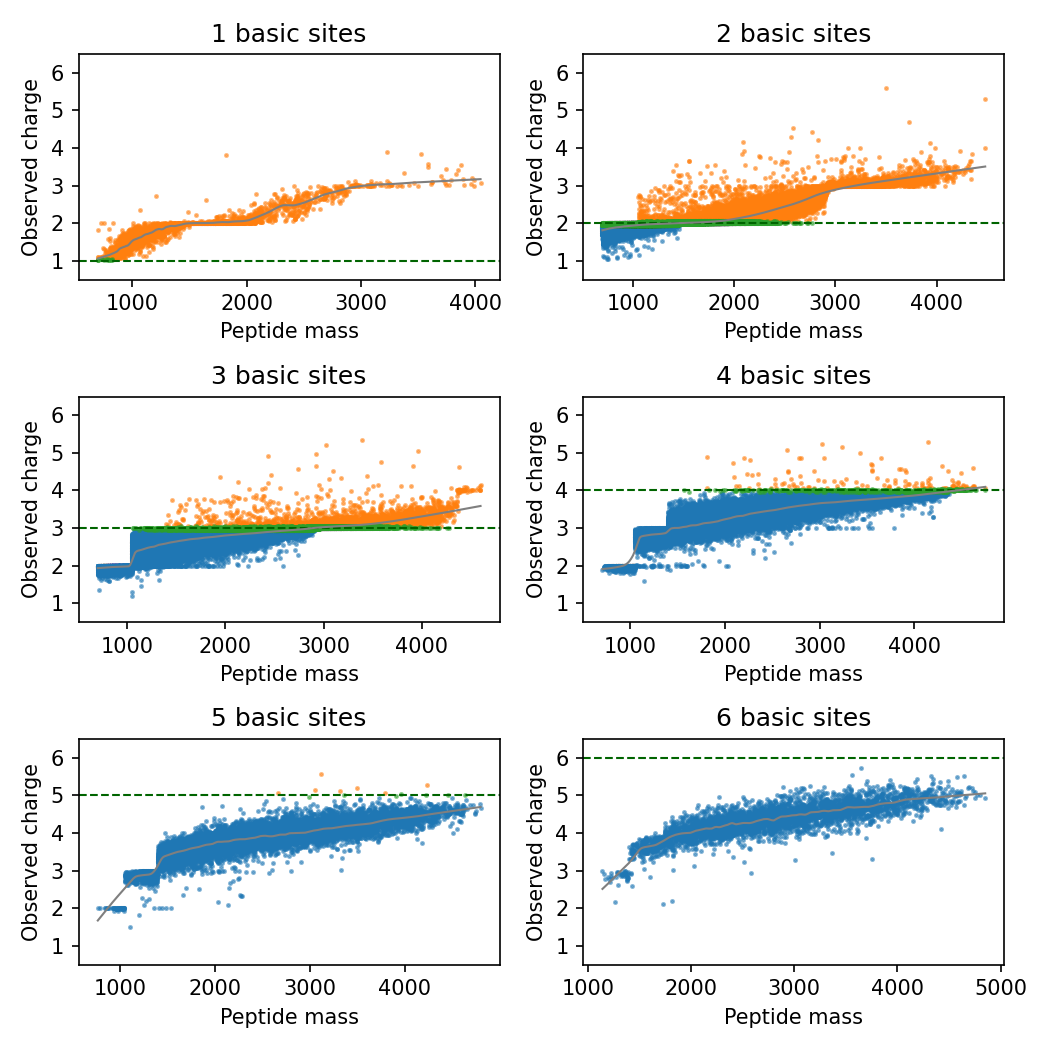

### KNN_regression.png

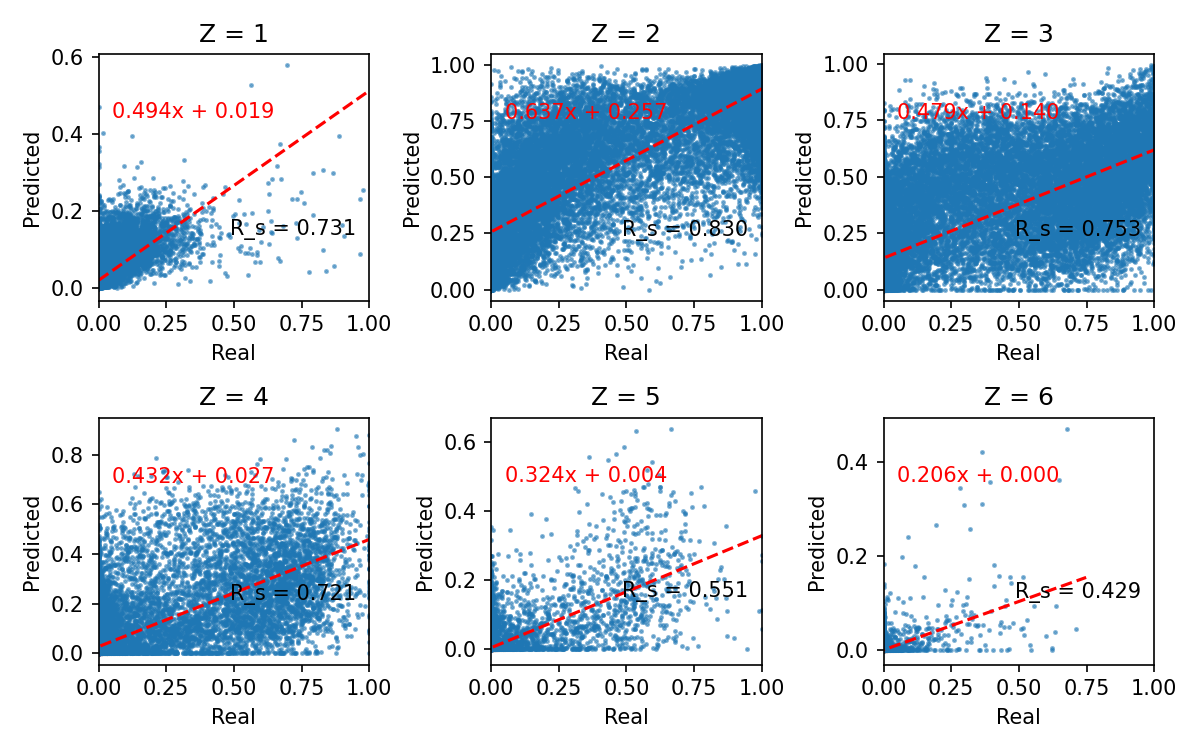

### linear_regression.png

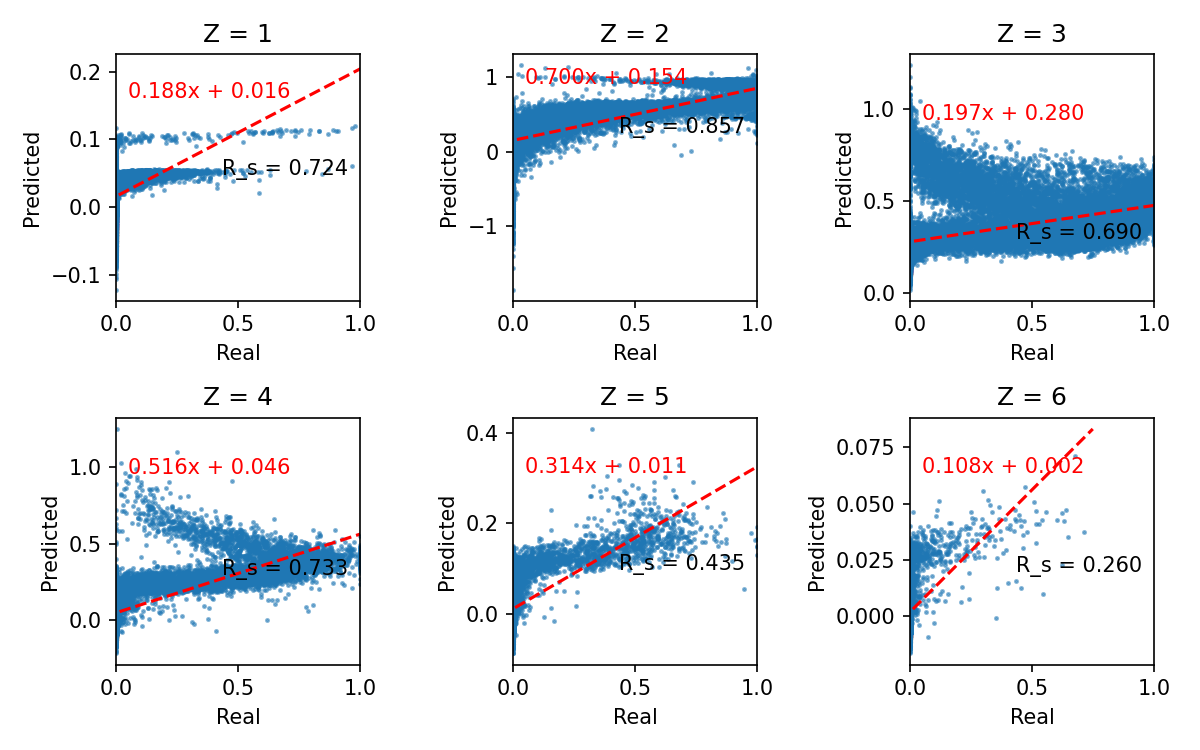

### LSTM_regression.png

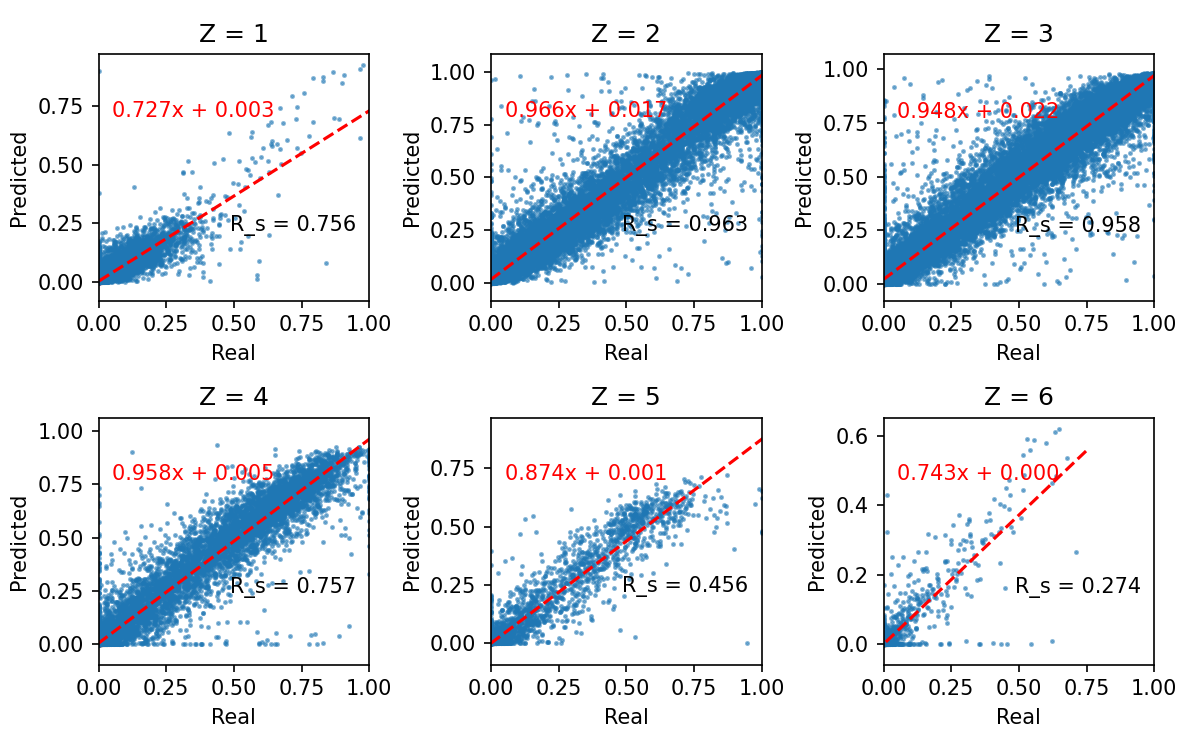

### phospho_under_overcharging.png

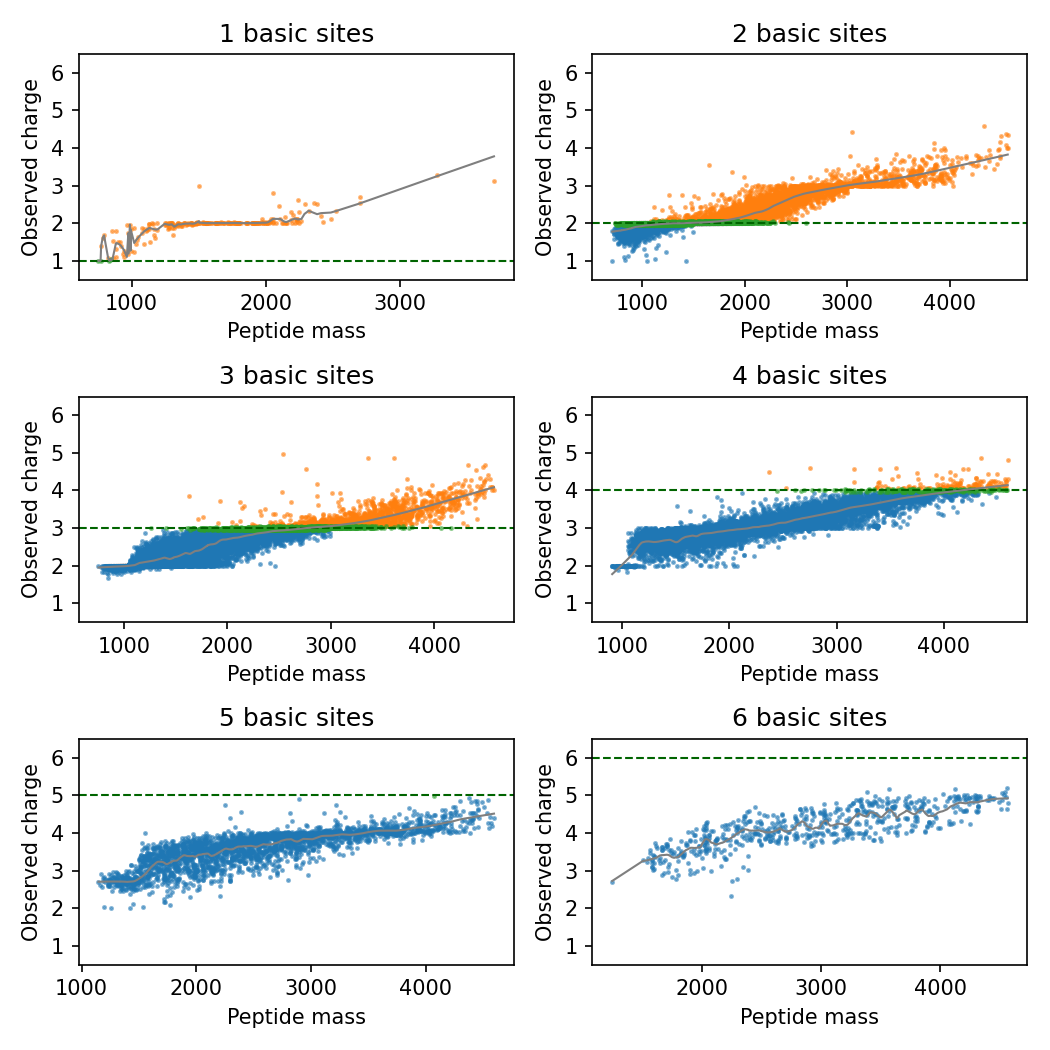

### SVR_multi_regression.png

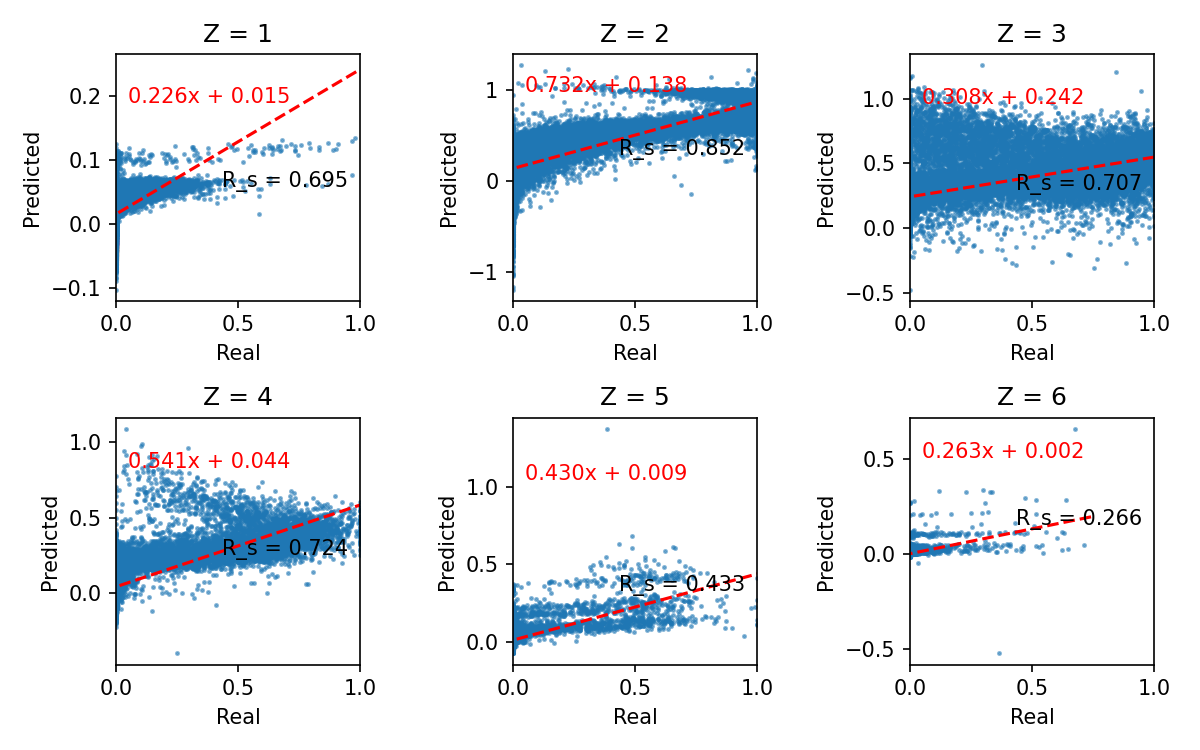
