## Supplementary material for "Exploiting Charge State Distribution to Probe Intramolecular Interactions in Gas-Phase Phosphopeptides and Enhance Proteomics Analyses": Scripts for data analysis and generation of results and intermediate plots: CCS_Z.py

```
# -*- coding: utf-8 -*-
"""
Analysis of CCS and charge state dimemsions of phosphopeptides
Need:
    evidence file from Ogata et al for CCS
    complete (phospho)peptides CSD table

Output:
    Plots and graphs

Usage:
    plain script (not to be executed from command line)

@author: Vladimir
"""

import re
import pandas as pd
import numpy as np
import matplotlib.pyplot as plt
from pyteomics.proforma import ProForma

modification_mapping = {'(Acetyl (Protein N-term))': '[Acetylation]-',
                        '(Phospho (STY))': '[Phosphorylation]',
                        '(Oxidation (M))': '[Oxidation]'}

evidence_path = 'D:/RawData/PhosphoTims/evidence.txt'
min_loc_prob = 0.95

def convert_to_proforma(string, mod_map):
    string = string.strip('_')
    for (_to, _from) in mod_map.items():
        string = string.replace(_to, _from)

    string = re.sub('C', 'C[Carbamidomethyl]', string)

    return string

def parse_loc_prob(l, min_loc_prob):
    if type(l) == list:
        return int(np.sum([float(x) >= min_loc_prob for x in l]))
    else:
        return 0

def get_ccs_shifts(frame):
    return pd.DataFrame({'proforma': frame.loc[frame['nr_phospho'] > 0, 'proforma'],
                         'shift': (frame.loc[frame['nr_phospho'] > 0, 'CCS'].values -\
                                   frame.loc[frame['nr_phospho'] == 0, 'CCS'].values),
                         'CCS': frame.loc[frame['nr_phospho'] == 0, 'CCS'].values[0]})

#charge shift by proforma
def get_z_shifts(frame):
    return pd.DataFrame({'proforma': frame.loc[frame['Phospho_count'] > 0, 'proforma'],
                         'shift': (frame.loc[frame['Phospho_count'] > 0, 'charge'].values -\
                                   frame.loc[frame['Phospho_count'] == 0, 'charge'].values),
                         'charge': frame.loc[frame['Phospho_count'] == 0, 'charge'].values[0]})

data = pd.read_csv(evidence_path, sep='\t')

#extracting data from the evidence file
data['proforma'] = data['Modified sequence'].apply(convert_to_proforma, args=( modification_mapping,))
data['nr_phospho'] = data['proforma'].str.count('\[Phosphorylation\]')
data['nr_sites'] = data['Sequence'].str.count('[STY]')
data['nr_located_sites'] = data['Phospho (STY) Probabilities'].str.findall('\(([0-9\.]+)\)')\
.apply(parse_loc_prob, args=(min_loc_prob,)) #parsing located sites

peptides = data[['proforma', 'nr_phospho', 'nr_sites', 'nr_located_sites', 'Charge', 'Mass', 'CCS']].copy()
peptides.rename({'Charge': 'charge'}, axis=1, inplace=True)

#removing multiphosphorylated
peptides = peptides[np.logical_and(peptides['nr_phospho']  == peptides['nr_located_sites'],
                                   peptides['nr_phospho'] < 2)].copy()

#removing replicate peptides if their CCS values has more than 5% difference
peptides = peptides.groupby(['proforma', 'charge'])\
.filter(lambda f: np.max(f['CCS'].values - f['CCS'].mean()) / f['CCS'].mean() < 0.05)
#averaging properties for peptides and charge-states: CCSes etc
peptides = peptides.groupby(['proforma', 'charge']).mean().reset_index()

peptides['dephosphorylated'] = peptides['proforma'].str.replace('[Phosphorylation]', '', regex=False)

paired  = peptides.groupby(['dephosphorylated', 'charge']).filter(lambda f: np.logical_and(np.any(f['nr_phospho'] > 0),
                                                                                      np.any(f['nr_phospho'] == 0)))

ccs_shifts = paired.groupby(['dephosphorylated', 'charge']).apply(get_ccs_shifts).reset_index().drop('level_2', axis=1)

ccs_shifts.to_csv('D:/RawData/PhosphoTims/ccs_shift.csv', index=False)

#loading charge data
peptides = pd.read_csv('D:/RawData/CSD/peptides+phospho_more.csv')
peptides['dephosphorylated'] = peptides['proforma'].str.replace('[Phosphorylation]', '', regex=False)

#find groups of peptides with the same dephosphorylated sequence and at least one plain and one phosphorylated peptide
phospho_paired = peptides.groupby('dephosphorylated').filter(lambda f: np.logical_and(np.any(f['Phospho_count'] > 0),
                                                                                      np.any(f['Phospho_count'] == 0)))

z_shifts = phospho_paired.groupby('dephosphorylated').apply(get_z_shifts)
z_shifts['Phospho_count'] = z_shifts['proforma'].str.count('\[Phosphorylation\]')
summed_shifts = ccs_shifts.merge(z_shifts, on='proforma', suffixes=('_ccs', '_z'))
summed_shifts['mass'] = summed_shifts['dephosphorylated'].apply(lambda s: ProForma.parse(s).mass)
summed_shifts.to_csv('D:/RawData/PhosphoTims/ccs+z_shift.csv', index=False)

multi_shift = summed_shifts.groupby('proforma').filter(lambda f: f.shape[0] > 1)
single_shift = summed_shifts.groupby('proforma').filter(lambda f: f.shape[0] == 1)

print('CCS Charge distribution:\n', single_shift.groupby('charge_ccs').size())

plt.figure(figsize=(8, 3))
n = 1
for z in [2, 3]:
    frame = single_shift[single_shift['charge_ccs'] == z]
    plt.subplot(1, 2, n)
    x = frame['shift_ccs']
    y = frame['shift_z']

    plt.scatter(x, y, s=1, c='tab:gray')

    plt.axvline(0, ls='--', color='tab:red')
    plt.axhline(0, ls='--', color='tab:red')

    p = np.logical_and(x >= 0, y >= 0).sum()
    plt.text(plt.xlim()[1] - 3, plt.ylim()[1] - 0.05, f'{p}', color='k',
             va='top', ha='right')
    p = np.logical_and(x < 0, y < 0).sum()
    plt.text(plt.xlim()[0] + 3, plt.ylim()[0] + 0.05, f'{p}', color='k',
             va='bottom', ha='left')
    p = np.logical_and(x >= 0, y < 0).sum()
    plt.text(plt.xlim()[1] - 3, plt.ylim()[0] + 0.05, f'{p}', color='k',
             va='bottom', ha='right')
    p = np.logical_and(x < 0, y >= 0).sum()
    plt.text(plt.xlim()[0] + 3, plt.ylim()[1] - 0.05, f'{p}', color='k',
             va='top', ha='left')

    plt.text(plt.xlim()[0] + 3, plt.ylim()[1] / 2, f'R={np.corrcoef(x, y)[0, 1]:.3f}', color='k',
             va='center', ha='left')

    plt.xlabel('$\Delta$ CCS [Ph - nonPh]')
    plt.ylabel('$\Delta$ Charge [Ph - nonPh]')
    plt.title(f'CCS Charge: {z}+')
    n += 1

    plt.tight_layout()
plt.savefig('D:/RawData/CSD/pub/plots/phospho/ccs+z_scatter.svg')


changes = []
for s, g in multi_shift.groupby('proforma'):
    sg = g.sort_values('charge_ccs')
    for i1, i2 in zip(*np.triu_indices(sg.shape[0], 1)):
        changes.append({'ccs_1': sg.iloc[i1,:]['shift_ccs'],
                        'ccs_2': sg.iloc[i2,:]['shift_ccs'],
                        'charges': '{} - {}'.format(sg.iloc[i1,:]['charge_ccs'], sg.iloc[i2,:]['charge_ccs'])})

changes = pd.DataFrame(changes)

changes['same_direction'] = changes['ccs_1'] * changes['ccs_2'] > 0
changes['delta'] = changes['ccs_1'] - changes['ccs_2']
changes['abs_delta'] = np.abs(changes['ccs_1'] - changes['ccs_2'])
changes['color'] = changes['same_direction'].apply(lambda s: 'tab:orange' if s else 'tab:blue')

plt.figure(figsize=(8, 5.5))
for n, row in changes.sort_values('abs_delta').reset_index().iterrows():
    plt.plot([row['ccs_1'], row['ccs_2']], [n, n], color=row['color'], lw=2, alpha=0.8)

plt.axvspan(changes['ccs_2'].min() - 5, 0, color='cyan', alpha=0.2)
plt.text(changes['ccs_2'].min() - 2, 0, 'compaction', ha='left', va='bottom', color='blue')
plt.axvspan(0, changes['ccs_2'].max() + 5, color='orange', alpha=0.2)
plt.text(changes['ccs_2'].max() + 2, 0, 'expansion', ha='right', va='bottom', color='orange')

custom_lines = [plt.Line2D([0], [0], color='tab:orange', lw=2),
                plt.Line2D([0], [0], color='tab:blue', lw=2)]

descriptions = ['Same direction',
                'Opposite direction']

plt.xlim(changes['ccs_2'].min() - 5, changes['ccs_2'].max() + 5)
plt.legend(custom_lines, descriptions, loc='center left')

plt.savefig('D:/RawData/CSD/pub/plots/phospho/compaction_CCS_trends.svg')
```
