## Supplementary material for "Exploiting Charge State Distribution to Probe Intramolecular Interactions in Gas-Phase Phosphopeptides and Enhance Proteomics Analyses": Scripts for data analysis and generation of results and intermediate plots: CSD_Learning_Phospho_cos.ipynb

```
# %% [code]
import numpy as np
import pandas as pd
from sklearn.model_selection import train_test_split
from keras import models, layers, callbacks
from keras.metrics import cosine_similarity, mean_squared_error

# %% [code]
psi_to_single_ptm = {'[Acetyl]-': 'B',
                     '[Carbamidomethyl]': '',
                     'M[Oxidation]': 'J',
                     'S[Phosphorylation]': 'X',
                     'T[Phosphorylation]': 'O',
                     'Y[Phosphorylation]': 'U'}

def reshapeOneHot(X):
    X = np.dstack(X)
    X = np.swapaxes(X, 1, 2)
    X = np.swapaxes(X, 0, 1)
    return X

def get_single_ptm_code(psi_sequence):
    sequence = psi_sequence
    for ptm in psi_to_single_ptm:
        sequence = sequence.replace(ptm, psi_to_single_ptm[ptm])
    return sequence

def one_hot_encode_peptide(psi_sequence, MAX_LENGTH = 40):
    peptide = get_single_ptm_code(psi_sequence)
    if len(peptide) > MAX_LENGTH:
        print('Peptide length is larger than maximal length of ', str(MAX_LENGTH))
        return ['']
    else:
        AA_vocabulary = 'KRPTNAQVSGILCMJHFYWEDBXOU'#B: acetyl; J: oxidized Met; O: PhosphoT; X: PhosphoS; U: PhosphoY
        
        one_hot_peptide = np.zeros((len(peptide), len(AA_vocabulary)))

        for j in range(0, len(peptide)):
            try:
                aa = peptide[j]
                one_hot_peptide[j, AA_vocabulary.index(aa)] = 1
            except:
                print(f'WARN: "{aa}" is not in vocabulary; it will be skipped')

        no_front_paddings = int((MAX_LENGTH - len(peptide))/2)
        peptide_front_paddings = np.zeros((no_front_paddings, one_hot_peptide.shape[1]))

        no_back_paddings = MAX_LENGTH - len(peptide) - no_front_paddings
        peptide_back_paddings = np.zeros((no_back_paddings, one_hot_peptide.shape[1]))

        full_one_hot_peptide = np.vstack((peptide_front_paddings, one_hot_peptide, peptide_back_paddings))

        return peptide, full_one_hot_peptide

# %% [code]
#preparing the data

data = pd.read_csv('peptides+phospho.csv')
data.fillna(0, inplace=True)
data['parsed_sequence'] = data['proforma'].apply(lambda s: one_hot_encode_peptide(s)[0])

print((data['parsed_sequence'] == '').sum(), ' sequences were not parsed')

# %% [code]
#slicing initial dataset in different sizes
from os import mkdir

try:
    mkdir('splits')
except Exception:
    pass

index = list(data.index)
np.random.shuffle(index)

for size in [0.1, 0.3, 1.0]:
    data_slice = data.loc[index[:int(len(index) * size)], :]
    train, test = train_test_split(data_slice, test_size=0.1)
    train.to_csv(f'splits/peptides+phospho_{size}_train.csv', index=False)
    test.to_csv(f'splits/peptides+phospho_{size}_test.csv', index=False)

# %% [code]
for size in [0.1, 0.3, 1.0]:
    print(f'Working with set size {size}')
    data = pd.read_csv(f'splits/peptides+phospho_{size}_train.csv', na_filter=False, low_memory=False)
    train_X = reshapeOneHot(data['proforma'].apply(lambda s: one_hot_encode_peptide(s)[1]).values)
    train_Y = data.loc[:, '1':'6'].values
    print(f'Train size {data.shape[0]}')

    data = pd.read_csv(f'splits/peptides+phospho_{size}_test.csv', na_filter=False, low_memory=False)
    test_X = reshapeOneHot(data['proforma'].apply(lambda s: one_hot_encode_peptide(s)[1]).values)
    test_Y = data.loc[:, '1':'6'].values
    print(f'Test size {data.shape[0]}')

    #Model
    network = models.Sequential()
    network.add(layers.Masking(mask_value=0., input_shape=(None, train_X.shape[2])))
    network.add(layers.Bidirectional(layers.LSTM(256,
                             dropout=0.1,
                             recurrent_dropout=0.5,
                             return_sequences=True)))
    network.add(layers.Bidirectional(layers.LSTM(256,
                             dropout=0.1,
                             recurrent_dropout=0.5)))
    network.add(layers.Dense(256, activation='tanh'))
    network.add(layers.Dense(6, activation='softmax'))
    print(network.summary())

    network.compile(optimizer='adam', loss='cosine_similarity')
    
    #fitting
    print('Training...')
    earlystop = callbacks.EarlyStopping(monitor='val_loss', patience=5, mode='min', restore_best_weights=True)
    history = network.fit(train_X,
                  train_Y,
                  epochs=100,
                  batch_size=512,
                  validation_split=0.2,
                  callbacks=(earlystop,))

    #Testing
    cost = network.evaluate(test_X, test_Y, batch_size=1024)
    print(f'Final cost: {cost:.6f}')
    
    #only phospho
    peptides = data.loc[data['proforma'].str.count('\[Phosphorylation\]') > 0, :].copy()

    X = reshapeOneHot(peptides['proforma'].apply(lambda s: one_hot_encode_peptide(s)[1]).values)
    Y_pred = network.predict(X, batch_size=2048)
    Y_true = peptides.loc[:, '1':'6'].fillna(0).values.astype('float32')
    print(f'Error on {peptides.shape[0]} phosphopeptides: {np.mean(cosine_similarity(Y_true, Y_pred)):.6f}')

    #only nonphospho
    peptides = data.loc[data['proforma'].str.count('\[Phosphorylation\]') == 0, :].copy()

    X = reshapeOneHot(peptides['proforma'].apply(lambda s: one_hot_encode_peptide(s)[1]).values)
    Y_pred = network.predict(X, batch_size=2048)
    Y_true = peptides.loc[:, '1':'6'].fillna(0).values.astype('float32')
    print(f'Error on {peptides.shape[0]} non-phosphopeptides: {np.mean(cosine_similarity(Y_true, Y_pred)):.6f}')

    #save model
    network.save(f'splits/CSD_model_LCMSMS_{size}.h5', include_optimizer=False)
```
