## Supplementary material for "Exploiting Charge State Distribution to Probe Intramolecular Interactions in Gas-Phase Phosphopeptides and Enhance Proteomics Analyses": Scripts for data analysis and generation of results and intermediate plots: digest_exploration.py

```
# -*- coding: utf-8 -*-
"""
Investigating CSDs properties of the complete human proteome digest
Need:
    trained CSD model
    DeepLC

Output:
    Plots and graphs

Usage:
    plain script (not to be executed from command line)

@author: Vladimir
"""
import pandas as pd
import numpy as np
import matplotlib.pyplot as plt
from pyteomics.proforma import ProForma
from pyteomics import parser, fasta
from pyteomics.mass import Composition, calculate_mass
from functools import reduce
from collections import Counter, defaultdict
from sklearn.metrics.pairwise import cosine_similarity
from sklearn.cluster import AgglomerativeClustering
from keras import models

#mapping of PTMs in CSD model
psi_to_single_ptm = {'[Acetyl]-': 'B',
                     '[Carbamidomethyl]': '',
                     'M[Oxidation]': 'J',
                     'S[Phosphorylation]': 'X',
                     'T[Phosphorylation]': 'O',
                     'Y[Phosphorylation]': 'U'}


def proforma_to_composition(proforma_string):
    p = ProForma.parse(proforma_string)
    mods = []
    sequence = []

    for aa, mod in p.sequence:
        sequence.append(aa)

        if not mod is None:
            for m in mod:
                mods.append(m.composition)

    if not p.n_term is None:
        for m in p.n_term:
            mods.append(m.composition)

    if not p.c_term is None:
        for m in p.c_term:
            mods.append(m.composition)

    composition = Composition(sequence=''.join(sequence)) + reduce(lambda a,b: a + b, mods, Composition())
    return ''.join([f'{k}{composition[k]}' for k in sorted(composition.keys())])

def cosine(data):
    if data.shape[0] > 1:
        splits = 1
        intermediate = []
        while (splits < 10):
            try:
                borders = np.round(np.linspace(0, data.shape[0], splits + 1), 0).astype(int)
                for i in range(1, splits + 1):
                    for j in range(i, splits + 1):
                        if i == j:
                            intermediate.append(cosine_similarity(data.loc[:, 'z=1':'z=6'].values[borders[i - 1]:borders[i], :])[np.triu_indices(borders[i] - borders[i - 1], 1)].mean())
                        else:
                            intermediate.append(cosine_similarity(data.loc[:, 'z=1':'z=6'].values[borders[i - 1]:borders[i], :],
                                                                  data.loc[:, 'z=1':'z=6'].values[borders[j - 1]:borders[j], :]).mean())

                return np.mean(intermediate)
            except MemoryError:
                print(f'Cannot calculate cosine cross-cimilarity for {data.shape[0]} elements with {splits} splits')
                splits += 1

        raise MemoryError(f'Cannot calculate cosine cross-cimilarity for {data.shape[0]} elements with {splits} splits')

    else:
        return np.nan

def cluster_size(data, metric='cosine', linkage='average', threshold=0.05):
    if data.shape[0] > 1:
        AC = AgglomerativeClustering(n_clusters=None, affinity=metric, linkage=linkage, distance_threshold=threshold)
        AC.fit(data.loc[:, 'z=1':'z=6'])
        return Counter(Counter(AC.labels_).values())
    else:
        return {1: 1}

#this version returns if we can separate anything at all
def cluster_size_c(data, metric='cosine', linkage='average', threshold=0.05):
    if data.shape[0] > 1:
        AC = AgglomerativeClustering(n_clusters=None, affinity=metric, linkage=linkage, distance_threshold=threshold)
        AC.fit(data.loc[:, 'z=1':'z=6'])
        return max(Counter(Counter(AC.labels_).values()).keys()) < data.shape[0]
    else:
        return np.nan

def count_letters(sequence, relative=False):
    counts = Counter(sequence)
    if relative:
        length = len(sequence)
        return {f'rel_{k}': counts[k]/length for k in sorted(counts.keys())}
    else:
        return ''.join([f'{k}{counts[k]}' for k in sorted(counts.keys())])


def in_silico_digestion(fasta_file, enzyme, missed_cleavages, min_length, max_length):
    # Initialize empty list to store peptides
    peptides = []

    # Use pyteomics to perform in silico digestion
    for header, sequence in fasta.read(fasta_file):
        # Perform in silico digestion
        digest = parser.cleave(sequence, enzyme, missed_cleavages=missed_cleavages,
                               min_length=min_length, max_length=max_length)
        # Add the filtered peptides to the list
        peptides.extend(digest)

    # Convert the list of peptides to a pandas DataFrame
    df = pd.DataFrame({'sequence': peptides})

    return df

def composition_to_string(composition):
    return ''.join([f'{element}{composition[element]}' for element in sorted(composition.keys())])

def sequence_to_composition(sequence):
    return composition_to_string(Composition(sequence=sequence))

def get_peprec_modifications(proforma_string):
    result = []
    p = ProForma.parse(proforma_string)

    if not p.n_term is None:
        for mod in p.n_term:
            result.append(f'0|{mod}')

    for i, (aa, mods) in enumerate(p.sequence):
        if not mods is None:
            for mod in mods:
                result.append(f'{i + 1}|{mod}')

    if not p.c_term is None:
        for mod in p.c_term:
            result.append(f'-1|{mod}')

    return '|'.join(result)

#feature preapration functions for CSD model
def reshapeOneHot(X):
    X = np.dstack(X)
    X = np.swapaxes(X, 1, 2)
    X = np.swapaxes(X, 0, 1)
    return X

def get_single_ptm_code(psi_sequence):
    sequence = psi_sequence
    for ptm in psi_to_single_ptm:
        sequence = sequence.replace(ptm, psi_to_single_ptm[ptm])
    return sequence

def one_hot_encode_peptide(psi_sequence, MAX_LENGTH = 40):
    peptide = get_single_ptm_code(psi_sequence)
    if len(peptide) > MAX_LENGTH:
        print('Peptide length is larger than maximal length of ', str(MAX_LENGTH))
        return ['', None]
    else:
        AA_vocabulary = 'KRPTNAQVSGILCMJHFYWEDBXOU'

        one_hot_peptide = np.zeros((len(peptide), len(AA_vocabulary)))

        for j in range(0, len(peptide)):
            try:
                aa = peptide[j]
                one_hot_peptide[j, AA_vocabulary.index(aa)] = 1
            except:
                print(f'WARN: "{aa}" is not in vocabulary; it will be skipped')

        no_front_paddings = int((MAX_LENGTH - len(peptide))/2)
        peptide_front_paddings = np.zeros((no_front_paddings, one_hot_peptide.shape[1]))

        no_back_paddings = MAX_LENGTH - len(peptide) - no_front_paddings
        peptide_back_paddings = np.zeros((no_back_paddings, one_hot_peptide.shape[1]))

        full_one_hot_peptide = np.vstack((peptide_front_paddings, one_hot_peptide, peptide_back_paddings))

        return peptide, full_one_hot_peptide


#in silico digestion 2 missed cleavages 6-35 AA
digest = in_silico_digestion('D:/Fasta/uniprot-human.fasta', 'trypsin', 2, 6, 35)

#removing redundant and strange sequences
digest.drop_duplicates(inplace=True)
digest.drop(digest.index[digest['sequence'].str.find('X') != -1], axis=0, inplace=True)
digest.reset_index(drop=True, inplace=True)

#add CAM-C
digest['ProForma'] = digest['sequence'].str.replace('C', 'C[Carbamidomethyl]')

#calculate elemental compositions
digest['composition'] = digest['ProForma'].apply(proforma_to_composition)

#calculate amino acid compositions
digest['AA_composition'] = digest['sequence'].apply(count_letters)

#grouping by composition
composition_digest = digest.groupby('composition').agg({'sequence': 'size'})

#building linking column sequence <-> composition
composition_digest['compID'] = np.arange(composition_digest.shape[0])
digest['compID'] = composition_digest.loc[digest['composition'], 'compID'].values
composition_digest.reset_index(inplace=True)
composition_digest.set_index('compID', verify_integrity=True, inplace=True)

#grouping by AA composition
aa_digest = digest.groupby('AA_composition').agg({'sequence': 'size'})

#building linking column sequence <-> composition
aa_digest['isoID'] = np.arange(aa_digest.shape[0])
digest['isoID'] = aa_digest.loc[digest['AA_composition'], 'isoID'].values
aa_digest.reset_index(inplace=True)
aa_digest.set_index('isoID', verify_integrity=True, inplace=True)

#calculating masses
composition_digest['mass'] = composition_digest['composition'].apply(lambda s: calculate_mass(formula=s))

#add masses to digest
digest['mass'] = composition_digest.loc[digest['compID'], 'mass'].values

#generating PEPREC for DeepLC
digest['peprec_mod'] = digest['ProForma'].apply(get_peprec_modifications)
digest[['sequence', 'peprec_mod']].rename(columns={'sequence': 'seq', 'peprec_mod': 'modifications'}).to_csv('D:/RawData/CSD/positive_digest.peprec', index=False)

## DeepLC runnned externally

#reading predictions
rt_pred = pd.read_csv('D:/RawData/CSD/positive_digest_deeplc.csv', index_col=0)
digest['rRT'] = rt_pred['predicted_tr'].values

#CSD model
CSD_LCMSMS = models.load_model('D:/RawData/CSD/splits_phospho_cos/CSD_model_LCMSMS_1.0.h5', compile=False)
CSD_LCMSMS.compile()

#making CSD predictions
X = reshapeOneHot(digest['ProForma'].apply(lambda s: one_hot_encode_peptide(s)[1]).values)

Y_pred = CSD_LCMSMS.predict(X, batch_size=2048)

digest[[f'z={z}' for z in '123456']] = Y_pred

digest['charge'] = (digest.loc[:, 'z=1':'z=6'] * np.arange(1, 7, 1).reshape(1, 6)).sum(axis=1)

#saving the data
digest.to_csv('D:/RawData/CSD/positive_digest_peptides.csv', index=False)
composition_digest.to_csv('D:/RawData/CSD/positive_digest_compositions.csv', index=False)
aa_digest.to_csv('D:/RawData/CSD/positive_digest_peptides_aa.csv', index=False)

#binning with average charge
for bin_width in [1e-5, 5e-6, 2e-6]:
    for peaks in [100, 200, 400]:
        for z_width in [0.1, 0.2, 0.4]:
            print(f'MS width: {bin_width}; Peak capacity: {peaks}; Z width: {z_width}')
            if not f'mass_group_{bin_width*1e6:.0f}' in digest.columns:
                min_mass = digest['mass'].min() * (1 - bin_width / 2)
                digest[f'mass_group_{bin_width*1e6:.0f}'] = np.floor((np.log(digest['mass']) - np.log(min_mass)) / np.log(1 + bin_width)).astype(int)

            if not f'rt_group_{peaks}' in digest.columns:
                rt_width = 10 / peaks #relative RT from DeepLC are between 0 and 10
                min_rt = -1 * rt_width / 2
                digest[f'rt_group_{peaks}'] = np.floor((digest['rRT'] - min_rt) / rt_width).astype(int)

            if not f'z_group_{z_width}' in digest.columns:
                min_z = digest['charge'].min() - z_width / 2
                digest[f'z_group_{z_width}'] = np.floor((digest['charge'] - min_z) / z_width).astype(int)

            mass_groups = Counter(digest.groupby(f'mass_group_{bin_width*1e6:.0f}').size())
            mass_rt_groups = Counter(digest.groupby([f'mass_group_{bin_width*1e6:.0f}', f'rt_group_{peaks}']).size())
            mass_z_groups = Counter(digest.groupby([f'mass_group_{bin_width*1e6:.0f}', f'z_group_{z_width}']).size())
            mass_rt_z_groups = Counter(digest.groupby([f'mass_group_{bin_width*1e6:.0f}', f'rt_group_{peaks}', f'z_group_{z_width}']).size())

            plt.bar(np.array(list(mass_groups.keys())) - 0.3,
                    np.array(list(mass_groups.values())) * np.array(list(mass_groups.keys())) / digest.shape[0],
                    width=0.2, label='Mass')
            plt.bar(np.array(list(mass_z_groups.keys())) - 0.1,
                    np.array(list(mass_z_groups.values())) * np.array(list(mass_z_groups.keys())) / digest.shape[0],
                    width=0.2, label='Mass+Z')
            plt.bar(np.array(list(mass_rt_groups.keys())) + 0.1,
                    np.array(list(mass_rt_groups.values())) * np.array(list(mass_rt_groups.keys())) / digest.shape[0],
                    width=0.2, label='Mass+RT')
            plt.bar(np.array(list(mass_rt_z_groups.keys())) + 0.3,
                    np.array(list(mass_rt_z_groups.values())) * np.array(list(mass_rt_z_groups.keys())) / digest.shape[0],
                    width=0.2, label='Mass+RT+Z')

            plt.xticks(np.arange(1, max(mass_groups.keys())))
            plt.legend()
            plt.xlabel('Number of peptides')
            plt.ylabel('Frequency')
            plt.xlim((0.3, 20.7))
            plt.ylim(top=1.0)
            plt.title(f'Mass bin: {bin_width*1e6:.0f}ppm; RT bin: {peaks}peaks; Z bin: {z_width}')
            plt.savefig(f'D:/RawData/CSD/pub/plots/digest/digest_discrimination_{bin_width*1e6:.0f}ppm_{peaks}peaks_{z_width}z.svg')
            plt.close()

#binning with cosine similarity
for bin_width in [1e-5, 5e-6, 2e-6]:
    for peaks in [100, 200, 400]:
        for max_cos in [0.05, 0.025]:
            print(f'MS width: {bin_width}; Peak capacity: {peaks}; Max cos: {max_cos}')
            if not f'mass_group_{bin_width*1e6:.0f}' in digest.columns:
                min_mass = digest['mass'].min() * (1 - bin_width / 2)
                digest[f'mass_group_{bin_width*1e6:.0f}'] = np.floor((np.log(digest['mass']) - np.log(min_mass)) / np.log(1 + bin_width)).astype(int)

            if not f'rt_group_{peaks}' in digest.columns:
                rt_width = 10 / peaks #relative RT from DeepLC are between 0 and 10
                min_rt = -1 * rt_width / 2
                digest[f'rt_group_{peaks}'] = np.floor((digest['rRT'] - min_rt) / rt_width).astype(int)

            mass_groups = Counter(digest.groupby(f'mass_group_{bin_width*1e6:.0f}').size())
            mass_rt_groups = Counter(digest.groupby([f'mass_group_{bin_width*1e6:.0f}', f'rt_group_{peaks}']).size())

            mass_z_count = defaultdict(int)
            for element in digest.groupby([f'mass_group_{bin_width*1e6:.0f}']).apply(cluster_size, threshold=max_cos).tolist():
                for k, v in element.items():
                    mass_z_count[k] += v

            mass_rt_z_count = defaultdict(int)
            for element in digest.groupby([f'mass_group_{bin_width*1e6:.0f}', f'rt_group_{peaks}']).apply(cluster_size, threshold=max_cos).tolist():
                for k, v in element.items():
                    mass_rt_z_count[k] += v

            plt.bar(np.array(list(mass_groups.keys())) - 0.3,
                    np.array(list(mass_groups.values())) * np.array(list(mass_groups.keys())) / digest.shape[0],
                    width=0.2, label='Mass')
            plt.bar(np.array(list(mass_z_count.keys())) - 0.1,
                    np.array(list(mass_z_count.values())) * np.array(list(mass_z_count.keys())) / digest.shape[0],
                    width=0.2, label='Mass+Z')
            plt.bar(np.array(list(mass_rt_groups.keys())) + 0.1,
                    np.array(list(mass_rt_groups.values())) * np.array(list(mass_rt_groups.keys())) / digest.shape[0],
                    width=0.2, label='Mass+RT')
            plt.bar(np.array(list(mass_rt_z_count.keys())) + 0.3,
                    np.array(list(mass_rt_z_count.values())) * np.array(list(mass_rt_z_count.keys())) / digest.shape[0],
                    width=0.2, label='Mass+RT+Z')

            plt.xticks(np.arange(1, max(mass_groups.keys())))
            plt.legend()
            plt.xlabel('Number of peptides')
            plt.ylabel('Frequency')
            plt.xlim((0.3, 20.7))
            plt.ylim(top=1.0)
            plt.title(f'Mass bin: {bin_width*1e6:.0f}ppm; RT bin: {peaks}peaks; Z bin: {max_cos}cos')
            plt.savefig(f'D:/RawData/CSD/pub/plots/digest/digest_discrimination_{bin_width*1e6:.0f}ppm_{peaks}peaks_{max_cos}cos.svg')
            plt.close()

#cosine similarity of isomeric and isoelemental peptides
#preparing random shuffling of peptides
pep_index = list(digest.index)
np.random.shuffle(pep_index)

#cosine correlation of isomeric peptides
isom_cos = digest.groupby(['AA_composition', 'composition']).apply(cosine)
plt.hist(isom_cos, bins=np.linspace(0, 1, 51), histtype='step', label='Sequence isomers', density=True)
plt.axvline(np.nanmean(isom_cos), color='blue')
plt.text(np.nanmean(isom_cos) - 0.005, plt.ylim()[1] * 0.95, f'Mean:\n{np.nanmean(isom_cos):.4f}', color='blue', ha='right', va='top')

#cosine correlation of isoelemental peptides
iso_cos = digest.groupby(['composition']).apply(cosine)
plt.hist(iso_cos, bins=np.linspace(0, 1, 51), histtype='step', label='Constitutional isomers', density=True)
plt.axvline(np.nanmean(iso_cos), color='orange')
plt.text(np.nanmean(iso_cos) - 0.005, plt.ylim()[1] * 0.95, f'Mean:\n{np.nanmean(iso_cos):.4f}', color='orange', ha='right', va='top')

#cosine similarity of random peptides in dataset
cos_vector = cosine_similarity(digest.loc[pep_index[:25000], 'z=1':'z=6'].values,
                               digest.loc[pep_index[25000:50000], 'z=1':'z=6'].values)[np.eye(25000).astype(bool)]
plt.hist(cos_vector, bins=np.linspace(0, 1, 51), histtype='step', label='Random pairs', density=True)
plt.axvline(np.nanmean(cos_vector), color='green')
plt.text(np.nanmean(cos_vector) - 0.005, plt.ylim()[1] * 0.95, f'Mean:\n{np.nanmean(cos_vector):.4f}', color='green', ha='right', va='top')

plt.legend(loc='upper left')
plt.xlabel('Cosine similarity')
plt.ylabel('Density')
plt.savefig('D:/RawData/CSD/pub/plots/digest/digest_cosine_isomers.svg')
plt.close()

#cosine similarity for peptides in the same mass or RT groups
for bin_width in [1e-5, 5e-6, 2e-6]:
    for peaks in [100, 200, 400]:
        min_mass = digest['mass'].min() * (1 - bin_width / 2)
        digest['mass_group'] = np.floor((np.log(digest['mass']) - np.log(min_mass)) / np.log(1 + bin_width)).astype(int)

        rt_width = 10 / peaks
        min_rt = -1 * rt_width / 2
        digest['rt_group'] = np.floor((digest['rRT'] - min_rt) / rt_width).astype(int)

        #similarity of peptides of peptides with similar masses
        mass_cos = digest.groupby(['mass_group']).apply(cosine)
        plt.hist(mass_cos, bins=np.linspace(0, 1, 51), histtype='step', label='Mass', density=True)
        plt.axvline(np.nanmean(mass_cos), color='blue')

        #similarity of peptides with similar RT
        rt_cos = digest.groupby(['rt_group']).apply(cosine)
        plt.hist(rt_cos, bins=np.linspace(0, 1, 51), histtype='step', label='RT', density=True)
        plt.axvline(np.nanmean(rt_cos), color='orange')

        #similarity of peptides with similar Mass and RT
        mass_rt_cos = digest.groupby(['mass_group', 'rt_group']).apply(cosine)
        plt.hist(mass_rt_cos, bins=np.linspace(0, 1, 51), histtype='step', label='Mass+RT', density=True)
        plt.axvline(np.nanmean(mass_rt_cos), color='green')

        plt.legend(loc='upper left')
        plt.xlabel('Cosine similarity')
        plt.ylabel('Density')
        plt.savefig(f'D:/RawData/CSD/human_digest/cosine_discrimination_{bin_width*1e6:.0f}ppm_{peaks}peaks.png')
        plt.close()
```
