## Supplementary material for "Exploiting Charge State Distribution to Probe Intramolecular Interactions in Gas-Phase Phosphopeptides and Enhance Proteomics Analyses": Scripts for data analysis and generation of results and intermediate plots: extract_CSD_msms.py

```
# -*- coding: utf-8 -*-
"""
Extracting CSD from a single MS/MS file
Need:
    RAW file,
    mzML file,
    mzIdentML with annotated q-values
    features.tsv file from biosaur2

    all files has to share the name before the extension (basename)

Output:
    CSDs will be saved to {basename}_csd.csv

Usage:
    <script_name> <path_to_any_of_the_files>

    Some extraction parameters are stored in the #constant section

@author: Vladimir
"""
import sys
import json
import pandas as pd
import numpy as np
from os import path
from pyteomics.proforma import ProForma
from pyteomics.mzid import DataFrame
from pyteomics.mzml import read as read_mzml
from subprocess import check_output, CalledProcessError

#constant
mzTol = 5e-6
index_max = 2
PROTON = 1.00727646677

def mapModifications(psm):
    peptide = [''] + list(psm['PeptideSequence']) + ['']
    mods = psm['Modification']

    if not mods is np.nan:
        for mod in mods:
            loc = int(mod['location'])
            modtype = mod.get('name', mod['monoisotopicMassDelta'])
            peptide[loc] += f'[{modtype}]'

        if peptide[0] != '':
            peptide[0] = peptide[0] + '-'
        if peptide[-1] != '':
            peptide[-1] = '-' + peptide[-1]

    return ''.join(peptide)

def extract_feature_properties(group, faims_mode=False):
    group['Sequence'] = group['Match'].str.replace(r'\/\d+?', '', regex=True)
    sequence_vect = group.loc[group['Match'].astype(str) != '', 'Sequence'].str.split(';', expand=True).values.flatten()
    sequence_vect = np.unique(sequence_vect[sequence_vect != None])
    if len(sequence_vect) > 0:
        if faims_mode:
            feat_vect = [group.agg({'massCalib': np.mean, 'rtApex': np.mean, 'rtStart': np.min, 'rtEnd': np.max, 'FAIMS': np.mean}).values.reshape(1, -1) for _ in sequence_vect]
            frame = pd.DataFrame(np.concatenate(feat_vect), columns=['mass', 'rtApex', 'rtStart', 'rtEnd', 'FAIMS'], index=sequence_vect)
        else:
            feat_vect = [group.agg({'massCalib': np.mean, 'rtApex': np.mean, 'rtStart': np.min, 'rtEnd': np.max}).values.reshape(1, -1) for _ in sequence_vect]
            frame = pd.DataFrame(np.concatenate(feat_vect), columns=['mass', 'rtApex', 'rtStart', 'rtEnd'], index=sequence_vect)
        return frame

def expand_charges(row, min_charge, max_charge, mode):
    charge = np.arange(min_charge, max_charge + 1).reshape(-1, 1)
    mz = row['mass']/charge + PROTON * mode
    index = [f'{row["pepid"]}_{z[0]}' for z in charge]
    result = pd.DataFrame(mz, columns=['mz'], index=index)
    result['z'] = charge
    result['rt_start'] = row['rtStart']
    result['rt_end'] = row['rtEnd']
    result['rt'] = row['rtApex']
    result['Proforma'] = row['Proforma']
    if 'FAIMS' in row.keys():
        result['scan_filter'] = f'ms cv={row["FAIMS"]}'

    return result

def groupPFMs(features, faims_mode=False):
    features['pepid'] = -1
    print(f'Read {features.shape[0]} lines')

    pepid = 1

    for index in features.index[features['Match_count'] > 0]:
        if features.at[index, 'pepid'] == -1: #not assigned so far
                mass = features.at[index, 'massCalib']
                start = features.at[index, 'rtStart']
                end = features.at[index, 'rtEnd']
                cv = features.at[index, 'FAIMS']

                #select features based on similar mass and overlapping elution
                if faims_mode:
                    join_candidates = features[(np.abs(features['massCalib'] - mass) / mass < mzTol).astype(int) +\
                                            (features['rtStart'] <= end).astype(int) +\
                                            (features['rtEnd'] >= start).astype(int) +\
                                            (features['FAIMS'] == cv).astype(int) == 4]
                else:
                    join_candidates = features[(np.abs(features['massCalib'] - mass) / mass < mzTol).astype(int) +\
                                            (features['rtStart'] <= end).astype(int) +\
                                            (features['rtEnd'] >= start).astype(int) == 3]

                #require apex indices to be not more then index_max away
                group_ind = join_candidates.index[np.abs(join_candidates['scanApex'] - join_candidates.at[index, 'scanApex']) <= index_max]

                occupied = features.loc[group_ind, 'pepid'] != -1
                if np.any(occupied):
                    occupied_pepid = np.unique(features.loc[group_ind[occupied], "pepid"])
                    if len(occupied_pepid) > 1:
                        print(f'WARNING Mixed peptide group observed at {index}; candidate IDS: {group_ind}; PEPIDs: {occupied_pepid}')
                        print('Finding group with the smallest mass difference:')
                        best_delta = mass * mzTol
                        best_pepid = -1
                        for i in occupied_pepid:
                            delta = np.abs(mass - np.mean(features.loc[features['pepid'] == i, 'massCalib']))
                            if delta < best_delta:
                                best_pepid = i
                                best_delta = delta

                        print(f'BestID: {best_pepid}; BestDelta: {best_delta}')
                        features.loc[group_ind[~occupied], 'pepid'] = best_pepid

                    else:
                        features.loc[group_ind, 'pepid'] = occupied_pepid[0]

                else:
                    features.loc[group_ind, 'pepid'] = pepid
                    pepid += 1

    features.drop(features.index[features['pepid'] == -1], axis=0, inplace=True)

    print('Solving multiple assignements')
    #try solving multiple assignment cases (in quite a simplistic way)
    #also geting some stats
    counts = {'Cases': 0,
              'Joinable': 0,
              'Disjoint': 0,
              'Multicase': 0}

    for gid, group in features.groupby(['pepid', 'charge']):
        if group.shape[0] > 1:
            counts['Cases'] += 1
            z = gid[1]
            candidates = features[features['pepid'] == gid[0]].copy()
            #more than one double assignment per peptide
            if not np.all(candidates[candidates['charge'] != z].groupby('charge').count()['massCalib'] == 1):
                counts['Multicase'] += 1
                print(f'WARNING multiple cases per peptide:\n{candidates}')

            scans = group['mono_hills_scan_lists'].str.slice(1, -1).apply(np.fromstring, sep=',')
            sorted_index = (scans.apply(np.min).sort_values()).index
            #feature can be merged - all scan numbers are consequtive with max 1 gap
            if np.all(scans[sorted_index[1:]].apply(np.min).values - scans[sorted_index[0:-1]].apply(np.max).values <= 2):
                counts['Joinable'] += 1
                remove = group.index.drop(group.index[np.argmax(group['intensityApex'])])
                #mark as useless
                features.loc[remove, 'charge'] = 0
            else:
                counts['Disjoint'] += 1
                print(f'WARNING Disjoint features:\n{candidates}')
                #mark all as suspicious
                features.loc[group.index, 'charge'] = 0

    #drop useless features
    features.drop(features.index[features['charge'] == 0], inplace=True)
    print(f'Solving done:\n{counts}')

def AUC(x, y):
    return (0.5 * (y[1:] + y[:-1]) * (x[1:] - x[:-1])).sum()

def aggregate_csd(g):
    result = {k: v for k,v in zip(g['z'], g['area'])}
    result['ProForma'] = g['Proforma'].iloc[0]
    result['rt_start'] = g['rt_start'].iloc[0]
    result['rt_end'] = g['rt_end'].iloc[0]
    result['rt'] = g['rt'].iloc[0]
    result['pepid'] = g['pepid'].iloc[0]
    return result

def processFile(base_name):
    faims_mode = False
    def matchPSM(psm, faims_mode=False):
        if faims_mode:
            match_index = (features['rtStart'] < psm['RT']).astype(int) +\
                        (features['rtEnd'] > psm['RT']).astype(int) +\
                        (features['charge'] == psm['chargeState']).astype(int) +\
                        (np.abs((features['massCalib'] - psm['Mass'])/psm['Mass']) < mzTol).astype(int) +\
                        (features['FAIMS'] == psm['CV']).astype(int) == 5
        else:
            match_index = (features['rtStart'] < psm['RT']).astype(int) +\
                        (features['rtEnd'] > psm['RT']).astype(int) +\
                        (features['charge'] == psm['chargeState']).astype(int) +\
                        (np.abs((features['massCalib'] - psm['Mass'])/psm['Mass']) < mzTol).astype(int) == 4

        if sum(match_index) > 1:
            print('Warning: multiple PSM match\n', features.loc[match_index])

        elif sum(match_index) == 1:
            features.loc[match_index, 'Match'] += f';{psm["ProForma"]}'

    print(f'Starting with {base_name}')

    #reading features
    print(f'Reading features from {base_name}.features.tsv')
    features = pd.read_csv(f'{base_name}.features.tsv', sep='\t')
    print(f'Read {features.shape[0]} features')

    if np.sum(features['FAIMS'] != 0) > 0:
        print('FAIMS detected')
        faims_mode = True
        print(f'Reading scans from {base_name}.mzML')
        faims_map = {scan['id']: scan['FAIMS compensation voltage'] for scan in read_mzml(f'{base_name}.mzML')}
        print(f'Read {len(faims_map)} scans')

    #reading psms from mzid and filtering
    print(f'Reading identifications from {base_name}.mzid')
    identifications = DataFrame(f'{base_name}.mzid')
    identifications = identifications.query('`percolator:Q value` < 0.01').copy()
    identifications['ProForma'] = identifications.apply(mapModifications, axis=1)
    identifications['Mass'] = identifications['ProForma'].apply(lambda p: ProForma.parse(p).mass)
    identifications.rename({'scan start time': 'RT'}, axis=1, inplace=True)
    if faims_mode:
        identifications['CV'] = identifications['spectrumID'].apply(lambda x: faims_map[x])
    print(f'Read {identifications.shape[0]} PSMs')

    #mapping PSMs
    print('Mapping PSMs to features')
    features['Match'] = ''
    identifications.apply(matchPSM, axis=1, faims_mode=faims_mode)
    features['Match'] = features['Match'].apply(lambda s: ';'.join(set(s.strip(';').split(';'))))
    features['Match_count'] = features['Match'].str.count(';') + 1
    features.loc[features['Match'] == '', 'Match_count'] = 0
    print(f'Found PFMs:  {sum(features["Match_count"] > 0)}')

    #grouping PFM to peptides
    print('Grouping PFMs')
    groupPFMs(features, faims_mode=faims_mode)

    #merging PFM groups
    peptides = features.groupby('pepid').apply(extract_feature_properties, faims_mode=faims_mode)
    peptides.index = peptides.index.droplevel(0)
    peptides.index.names = ['Proforma']
    peptides.reset_index(inplace=True)
    peptides['pepid'] = np.arange(peptides.shape[0])
    print(f'Total peptides: {peptides.shape[0]}')

    #making mz-table
    mz_table = pd.concat(peptides.apply(expand_charges, axis=1, args=(1, 6, 1)).tolist())
    mzmin = np.floor(features['mz'].min())
    mzmax = np.ceil(features['mz'].max())
    mz_table = mz_table.query('mz >= @mzmin & mz <= @mzmax').copy()

    #writing JSON input for TRFP
    mz_table['tolerance'] = 5
    mz_table['tolerance_unit'] = 'ppm'
    mz_table['comment'] = mz_table.index
    with open(f'{base_name}_mz.json', 'w') as out:
        json.dump(mz_table.drop(['z', 'Proforma', 'rt'], axis=1).apply(lambda row: {k:v for k,v in zip(row.index, row)}, axis=1).tolist(), out)

    trfp_params = ['thermoparser', 'xic', '-i', f'{base_name}.raw', '-j', f'{base_name}_mz.json']

    try:
        print(f'Using TRFP to extract {mz_table.shape[0]} XICs')
        check_output(trfp_params, shell=True)
    except CalledProcessError as ex:
        print('TRFP execution failed\nOutput:\n{}', ex.output)
        raise Exception('TRFP XIC extraction failed')

    #reading TRFP output
    with open(f'{base_name}.json', 'r') as xic_input:
        chromatograms = json.load(xic_input)

    #calculating areas
    mz_table['area'] = [AUC(np.array(cr['RetentionTimes']), np.array(cr['Intensities'])) for cr in chromatograms['Content']]
    mz_table['pepid'] = mz_table['comment'].str.split('_', expand=True).iloc[:, 0].astype(int)

    #calculating CSD table
    csd_table = pd.DataFrame(mz_table.groupby('pepid').apply(aggregate_csd).tolist())
    csd_table.rename(columns={k:v for k,v in zip(csd_table.columns, csd_table.columns.astype(str))}, inplace=True)
    csd_table = csd_table.reindex(columns=sorted(csd_table.columns))
    csd_table.to_csv(f'{base_name}_csd.csv', index=False)
    print(f'{base_name} completed\n\n')

if __name__ == '__main__':
    processFile(path.splitext(sys.argv[1])[0])
```
