## Supplementary material for "Exploiting Charge State Distribution to Probe Intramolecular Interactions in Gas-Phase Phosphopeptides and Enhance Proteomics Analyses": Scripts for data analysis and generation of results and intermediate plots: extract_CSD_phospho.py

```
# -*- coding: utf-8 -*-
"""
Extracting CSD from a single MS/MS file
Need:
    MaxQuant msms.txt file,
    MaxQuant evidence.txt file (should share the name with msms.txt file),
    corresponding RAW files (should be named as provided in msms.txt file),
    and features.tsv files from biosaur2 for all of them

Output:
    CSDs will be saved to {rawfile_name}_csd.csv

Usage:
    <script_name> -f <path to msms.txt> -f <experiment format> -l <optinal path to library file>
                  -p <optional minimal site localization probability>

    experiment format is one of:
        MQ - regular MaxQuant (site probability and score threshold will be used)
        MQsynthetic - MaxQuant results validated against synthetic library as regexes (PXD000138)
        MQverify - MaxQuant results validated against synthetic library as reference sequences (PXD009449)

    Some extraction parameters are stored in the #constant section

@author: Vladimir
"""

import re
import logging
import json
import argparse
import pandas as pd
import numpy as np
from psm_utils.io import read_file
from os import path
from subprocess import check_output, CalledProcessError
from collections import Counter

#constant
mzTol = 5e-6
index_max = 2
PROTON = 1.00727646677

logging.basicConfig(format='%(asctime)s %(message)s', datefmt='%H:%M:%S', level=logging.INFO)

modification_mapping = {'ac': 'Acetylation',
                        'Acetyl': 'Acetylation',
                        'ph': 'Phosphorylation',
                        'Phospho (STY)': 'Phosphorylation',
                        'Phospho': 'Phosphorylation',
                        'ox': 'Oxidation',
                        'Oxidation (M)': 'Oxidation'}

def mq_to_dict(p):
    return {'proforma': re.sub(r'/\d+$', '', p.peptidoform.proforma),
            'sequence': p.peptidoform.sequence,
            'phospho_prob': p.metadata['Phospho (STY) Probabilities'],
            'theor_mz': p.peptidoform.theoretical_mz,
            'theor_mass': p.peptidoform.theoretical_mass,
            'evidence': int(p.metadata['Evidence ID']),
            'charge': p.peptidoform.precursor_charge,
            'run': p.run,
            'score': p.score,
            'rt': p.retention_time,
            'decoy': p.is_decoy}

def matchPSM(psm, features, faims_mode=False):
    if faims_mode:
        match_index = (features['rtStart'] < psm['rt']).astype(int) +\
                    (features['rtEnd'] > psm['rt']).astype(int) +\
                    (features['charge'] == psm['charge']).astype(int) +\
                    (np.abs((features['massCalib'] - psm['mass'])/psm['mass']) < mzTol).astype(int) +\
                    (features['FAIMS'] == psm['CV']).astype(int) == 5
    else:
        match_index = (features['rtStart'] < psm['rt']).astype(int) +\
                    (features['rtEnd'] > psm['rt']).astype(int) +\
                    (features['charge'] == psm['charge']).astype(int) +\
                    (np.abs((features['massCalib'] - psm['mass'])/psm['mass']) < mzTol).astype(int) == 4

    if sum(match_index) > 1:
        print('Warning: multiple PSM match\n', features.loc[match_index])

    elif sum(match_index) == 1:
        features.loc[match_index, 'match'] += f';{psm["proforma"]}'

def groupPFMs(features, faims_mode=False):
    features['pepid'] = -1
    pepid = 1

    for index in features.index[features['match_count'] > 0]:
        if features.at[index, 'pepid'] == -1: #not assigned so far
                mass = features.at[index, 'massCalib']
                start = features.at[index, 'rtStart']
                end = features.at[index, 'rtEnd']
                cv = features.at[index, 'FAIMS']

                #select features based on similar mass and overlapping elution
                if faims_mode:
                    join_candidates = features[(np.abs(features['massCalib'] - mass) / mass < mzTol).astype(int) +\
                                            (features['rtStart'] <= end).astype(int) +\
                                            (features['rtEnd'] >= start).astype(int) +\
                                            (features['FAIMS'] == cv).astype(int) == 4]
                else:
                    join_candidates = features[(np.abs(features['massCalib'] - mass) / mass < mzTol).astype(int) +\
                                            (features['rtStart'] <= end).astype(int) +\
                                            (features['rtEnd'] >= start).astype(int) == 3]

                #require apex indices to be not more then index_max away
                group_ind = join_candidates.index[np.abs(join_candidates['scanApex'] - join_candidates.at[index, 'scanApex']) <= index_max]

                occupied = features.loc[group_ind, 'pepid'] != -1
                if np.any(occupied):
                    occupied_pepid = np.unique(features.loc[group_ind[occupied], "pepid"])
                    if len(occupied_pepid) > 1:
                        logging.warning(f'WARNING Mixed peptide group observed at {index}; candidate IDS: {group_ind}; PEPIDs: {occupied_pepid}')
                        logging.info('Finding group with the smallest mass difference:')
                        best_delta = mass * mzTol
                        best_pepid = -1
                        for i in occupied_pepid:
                            delta = np.abs(mass - np.mean(features.loc[features['pepid'] == i, 'massCalib']))
                            if delta < best_delta:
                                best_pepid = i
                                best_delta = delta

                        logging.info(f'BestID: {best_pepid}; BestDelta: {best_delta}')
                        features.loc[group_ind[~occupied], 'pepid'] = best_pepid

                    else:
                        features.loc[group_ind, 'pepid'] = occupied_pepid[0]

                else:
                    features.loc[group_ind, 'pepid'] = pepid
                    pepid += 1

    features.drop(features.index[features['pepid'] == -1], axis=0, inplace=True)

    logging.info('Solving multiple assignements')
    #try solving multiple assignment cases (in quite a simple way)
    #also geting some stats
    counts = {'Cases': 0,
              'Joinable': 0,
              'Disjoint': 0,
              'Multicase': 0}

    for gid, group in features.groupby(['pepid', 'charge']):
        if group.shape[0] > 1:
            counts['Cases'] += 1
            z = gid[1]
            candidates = features[features['pepid'] == gid[0]].copy()
            #more than one double assignment per peptide
            if not np.all(candidates[candidates['charge'] != z].groupby('charge').count()['massCalib'] == 1):
                counts['Multicase'] += 1
                logging.warning(f'WARNING multiple cases per peptide:\n{candidates}')

            scans = group['mono_hills_scan_lists'].str.slice(1, -1).apply(np.fromstring, sep=',')
            sorted_index = (scans.apply(np.min).sort_values()).index
            #feature can be merged - all scan numbers are consequtive with max 1 gap
            if np.all(scans[sorted_index[1:]].apply(np.min).values - scans[sorted_index[0:-1]].apply(np.max).values <= 2):
                counts['Joinable'] += 1
                remove = group.index.drop(group.index[np.argmax(group['intensityApex'])])
                #mark as useless
                features.loc[remove, 'charge'] = 0
            else:
                counts['Disjoint'] += 1
                logging.warning(f'WARNING Disjoint features:\n{candidates}')
                #mark all as suspicious
                features.loc[group.index, 'charge'] = 0

    #drop useless features
    features.drop(features.index[features['charge'] == 0], inplace=True)
    logging.info(f'Solving done:\n{counts}')

def extract_feature_properties(group, faims_mode=False):
    group['sequence'] = group['match'].str.replace(r'\/\d+?', '', regex=True)
    sequence_vect = group.loc[group['match'].astype(str) != '', 'sequence'].str.split(';', expand=True).values.flatten()
    sequence_vect = np.unique(sequence_vect[sequence_vect != None])
    if len(sequence_vect) > 0:
        if faims_mode:
            feat_vect = [group.agg({'massCalib': np.mean, 'rtApex': np.mean, 'rtStart': np.min, 'rtEnd': np.max, 'FAIMS': np.mean}).values.reshape(1, -1) for _ in sequence_vect]
            frame = pd.DataFrame(np.concatenate(feat_vect), columns=['mass', 'rtApex', 'rtStart', 'rtEnd', 'FAIMS'], index=sequence_vect)
        else:
            feat_vect = [group.agg({'massCalib': np.mean, 'rtApex': np.mean, 'rtStart': np.min, 'rtEnd': np.max}).values.reshape(1, -1) for _ in sequence_vect]
            frame = pd.DataFrame(np.concatenate(feat_vect), columns=['mass', 'rtApex', 'rtStart', 'rtEnd'], index=sequence_vect)
        return frame

def AUC(x, y):
    return (0.5 * (y[1:] + y[:-1]) * (x[1:] - x[:-1])).sum()

def aggregate_csd(g):
    result = {k: v for k,v in zip(g['z'], g['area'])}
    result['proforma'] = g['proforma'].iloc[0]
    result['rt_start'] = g['rt_start'].iloc[0]
    result['rt_end'] = g['rt_end'].iloc[0]
    result['rt'] = g['rt'].iloc[0]
    result['pepid'] = g['pepid'].iloc[0]
    return result

def expand_charges(row, min_charge, max_charge, mode):
    charge = np.arange(min_charge, max_charge + 1).reshape(-1, 1)
    mz = row['mass']/charge + PROTON * mode
    index = [f'{row["pepid"]}_{z[0]}' for z in charge]
    result = pd.DataFrame(mz, columns=['mz'], index=index)
    result['z'] = charge
    result['rt_start'] = row['rtStart']
    result['rt_end'] = row['rtEnd']
    result['rt'] = row['rtApex']
    result['proforma'] = row['proforma']
    if 'FAIMS' in row.keys():
        result['scan_filter'] = f'ms cv={row["FAIMS"]}'

    return result

def mapModifications(psm):
    peptide = [''] + list(psm['sequence']) + ['']
    mods = psm['Modification']

    if not mods is np.nan:
        for mod in mods:
            loc = int(mod['location'])
            modtype = mod.get('name', mod['monoisotopicMassDelta'])
            peptide[loc] += f'[{modtype}]'

        if peptide[0] != '':
            peptide[0] = peptide[0] + '-'
        if peptide[-1] != '':
            peptide[-1] = '-' + peptide[-1]

    return ''.join(peptide)

#Read MaxQuant search results and filter with location probability
def read_MQ(msms_file, min_loc_prob=0.95):
    logging.info(f'Reading PSMs from {msms_file}..')
    psms = read_file(msms_file)
    psms.rename_modifications(modification_mapping)
    psms.add_fixed_modifications([('Carbamidomethyl', ['C'])])
    psms.apply_fixed_modifications()

    logging.info('Converting to DataFrame..')
    psm_frame = pd.DataFrame([mq_to_dict(psm) for psm in psms])
    psm_frame['nr_phospho'] = psm_frame['proforma'].str.count('\[Phosphorylation\]')
    psm_frame['nr_sites'] = psm_frame['sequence'].str.count('[STY]')
    psm_frame['nr_located_sites'] = psm_frame['phospho_prob'].str.findall('\(([0-9\.]+)\)').apply(lambda l: np.sum([float(x) >= min_loc_prob for x in l])).astype(int) #parsing located sites
    psm_frame['proforma'] = psm_frame['proforma'].str.replace('-[', '[', regex=False) #converting c-term modifications to modifications of the last AA

    logging.info(f'Read {psm_frame.shape[0]} PSMs')

    #correcting masses to match non calibrated features
    evidence = pd.read_csv(msms_file.replace('msms.txt', 'evidence.txt'), sep='\t', index_col='id', low_memory=False)
    ppm_shift = evidence.rename(lambda s: s.lower(), axis=1).reindex(psm_frame['evidence'])['uncalibrated mass error [ppm]'].values
    ppm_shift = np.nan_to_num(ppm_shift, np.nanmedian(ppm_shift))
    logging.info(f'Median mass shift: {np.median(ppm_shift):.3f} ppm')
    psm_frame['mass'] = psm_frame['theor_mass'] * (1 + ppm_shift * 1e-6)

    #high quality index
    hq = np.logical_and(
            np.logical_and(psm_frame['nr_phospho'] == psm_frame['nr_located_sites'], ~psm_frame['decoy']),
            np.logical_and(psm_frame['score'] > 50, psm_frame['nr_phospho'] > 0))

    logging.info(f'{hq.sum()} PSMs pass the quality filter')

    return psm_frame[hq].copy()

#Read MaxQuant results and filter with known library (PXD000138)
def read_MQ_library(msms_file, library_file):
    logging.info(f'Reading PSMs from {msms_file}..')
    psms = read_file(msms_file)
    psms.rename_modifications(modification_mapping)

    logging.info('Converting to DataFrame..')
    psm_frame = pd.DataFrame([mq_to_dict(psm) for psm in psms])
    psm_frame['nr_phospho'] = psm_frame['proforma'].str.count('\[Phosphorylation\]')
    psm_frame['nr_sites'] = psm_frame['sequence'].str.count('[STY]')

    logging.info(f'Read {psm_frame.shape[0]} PSMs')

    #correcting masses to match non calibrated features
    evidence = pd.read_csv(msms_file.replace('msms.txt', 'evidence.txt'), sep='\t', index_col='id', low_memory=False)
    ppm_shift = evidence.rename(lambda s: s.lower(), axis=1).reindex(psm_frame['evidence'])['uncalibrated mass error [ppm]'].values
    ppm_shift = np.nan_to_num(ppm_shift, np.nanmedian(ppm_shift))
    logging.info(f'Median mass shift: {np.median(ppm_shift):.3f} ppm')
    psm_frame['mass'] = psm_frame['theor_mass'] * (1 + ppm_shift * 1e-6)

    logging.info('Filtering input according to library..')
    library = pd.read_csv(library_file, index_col=0)

    invalid_indices = []
    for file, frame in psm_frame.groupby('run'):
        regex = library.at[int(file), 'Regex']
        invalid_indices.extend(frame.index[~frame['sequence'].str.match(regex)])

    psm_frame.drop(invalid_indices, axis=0, inplace=True)
    logging.info(f'Valid library PSMs: {psm_frame.shape[0]}')

    #high quality index
    hq = np.logical_and(
            np.logical_and(psm_frame['nr_phospho'] == psm_frame['nr_sites'], ~psm_frame['decoy']),
            np.logical_and(psm_frame['score'] > 50, psm_frame['nr_phospho'] > 0))

    logging.info(f'PSMs pass the quality filter: {hq.sum()}')

    return psm_frame[hq].copy()

#Read MaxQuant results and filter with known library (PXD009449)
def read_MQ_verify(msms_file, library_file):
    logging.info(f'Reading PSMs from {msms_file}..')
    psms = read_file(msms_file)
    psms.rename_modifications(modification_mapping)

    logging.info('Converting to DataFrame..')
    psm_frame = pd.DataFrame([mq_to_dict(psm) for psm in psms])

    logging.info(f'Read {psm_frame.shape[0]} PSMs')

    #correcting masses to match non calibrated features
    evidence = pd.read_csv(msms_file.replace('msms.txt', 'evidence.txt'), sep='\t', index_col='id', low_memory=False)
    ppm_shift = evidence.rename(lambda s: s.lower(), axis=1).reindex(psm_frame['evidence'])['uncalibrated mass error [ppm]'].values
    ppm_shift = np.nan_to_num(ppm_shift, np.nanmedian(ppm_shift))
    logging.info(f'Median mass shift: {np.median(ppm_shift):.3f} ppm')
    psm_frame['mass'] = psm_frame['theor_mass'] * (1 + ppm_shift * 1e-6)

    logging.info('Filtering input according to library..')
    library = set(pd.read_csv(library_file)['mod_sequence'])

    psm_frame = psm_frame.loc[psm_frame['proforma'].apply(lambda s: s in library), :].copy()
    logging.info(f'Valid library PSMs: {psm_frame.shape[0]}')

    #high quality index
    hq = np.logical_and(~psm_frame['decoy'], psm_frame['score'] > 50)

    logging.info(f'PSMs pass the quality filter: {hq.sum()}')

    return psm_frame[hq].copy()

def main(params):
    msms_file = params.msms_file
    input_type = params.input_type
    library_file = params.library_file
    minprob = params.minprob
    logging.info(f'Parameters: {params}')

    if input_type == 'MQ':
        psm_frame = read_MQ(msms_file, minprob)
    elif input_type == 'MQsynthetic':
        psm_frame = read_MQ_library(msms_file, library_file)
    elif input_type == 'MQverify':
        psm_frame = read_MQ_verify(msms_file, library_file)
    else:
        logging.critical(f'Unknown input format: {input_type}')
        raise Exception('Unknown input type')

    for filename, psms in psm_frame.groupby('run'):
        logging.info(f'Processing {filename}')

        if not path.exists(f'{path.split(msms_file)[0]}/{filename}.features.tsv'):
            logging.warning(f'Feature file not found for {filename}, the rest will be skipped')
            logging.info(f'Processing of {filename} completed\n')
            continue

        features = pd.read_csv(f'{path.split(msms_file)[0]}/{filename}.features.tsv', sep='\t')
        logging.info(f'Read {features.shape[0]} features')

        logging.info(f'Mapping {psms.shape[0]} PSMs..')
        features['match'] = ''
        psms.apply(matchPSM, axis=1, features=features)
        features['match'] = features['match'].apply(lambda s: ';'.join(set(s.strip(';').split(';'))))
        features['match_count'] = features['match'].str.count(';') + 1
        features.loc[features['match'] == '', 'match_count'] = 0
        logging.info(f'Found PFMs: {Counter(features["match_count"])}')

        logging.info('Grouping PFMs')
        groupPFMs(features)

        if features.shape[0] == 0:
            logging.warning(f'{filename} - no features left after grouping, the rest will be skipped')
            logging.info(f'Processing of {filename} completed\n')
            continue

        #merging PFM groups
        peptides = features.groupby('pepid').apply(extract_feature_properties)
        peptides.index = peptides.index.droplevel(0)
        peptides.index.names = ['proforma']
        peptides.reset_index(inplace=True)
        peptides['pepid'] = np.arange(peptides.shape[0])
        logging.info(f'Total peptides: {peptides.shape[0]}')

        #making mz-table
        mz_table = pd.concat(peptides.apply(expand_charges, axis=1, args=(1, 6, 1)).tolist())
        mzmin = np.floor(features['mz'].min())
        mzmax = np.ceil(features['mz'].max())
        mz_table = mz_table.query('mz >= @mzmin & mz <= @mzmax').copy()

        #writing JSON input for TRFP
        mz_table['tolerance'] = 5
        mz_table['tolerance_unit'] = 'ppm'
        mz_table['comment'] = mz_table.index
        with open(f'{path.split(msms_file)[0]}/{filename}_mz.json', 'w') as out:
            json.dump(mz_table.drop(['z', 'proforma', 'rt'], axis=1).apply(lambda row: {k:v for k,v in zip(row.index, row)}, axis=1).tolist(), out)

        trfp_params = ['thermoparser', 'xic', '-i', f'{path.split(msms_file)[0]}/{filename}.raw', '-j', f'{path.split(msms_file)[0]}/{filename}_mz.json']

        try:
            logging.info(f'Using TRFP to extract {mz_table.shape[0]} XICs')
            check_output(trfp_params, shell=True)
        except CalledProcessError as ex:
            logging.critical('TRFP execution failed\nOutput:\n{}', ex.output)
            raise Exception('TRFP XIC extraction failed')

        #reading TRFP output
        with open(f'{path.split(msms_file)[0]}/{filename}.json', 'r') as xic_input:
            chromatograms = json.load(xic_input)

        #integrity check
        if not np.all(mz_table['comment'] == [cr['Meta']['Comment'] for cr in chromatograms['Content']]):
            logging.critical('Misalignment between XICs and mz_table!')
            raise Exception('Misalignment; processing failed')
        #calculating areas
        mz_table['area'] = [AUC(np.array(cr['RetentionTimes']), np.array(cr['Intensities'])) for cr in chromatograms['Content']]
        mz_table['pepid'] = mz_table['comment'].str.split('_', expand=True).iloc[:, 0].astype(int)

        #calculating CSD table
        csd_table = pd.DataFrame(mz_table.groupby('pepid').apply(aggregate_csd).tolist())
        csd_table.rename(columns={k:v for k,v in zip(csd_table.columns, csd_table.columns.astype(str))}, inplace=True)
        csd_table = csd_table.reindex(columns=sorted(csd_table.columns))
        csd_table.to_csv(f'{path.split(msms_file)[0]}/{filename}_csd.csv', index=False)
        logging.info(f'Processing of {filename} completed\n')

if __name__ == '__main__':
    parser = argparse.ArgumentParser(description='Create CSD tables from DDA results')
    parser.add_argument('-f', '--msms-file', help='PSM file', required=True)
    parser.add_argument('-t', '--input-type', help='Format of PSM file', required=True)
    parser.add_argument('-l', '--library-file', help='Path to synthetic library')
    parser.add_argument('-p', '--minprob', help='Minimal probability of phospho-site', default=0.95, type=float)

    args = parser.parse_args()
    main(args)
```
