## Supplementary material for "Exploiting Charge State Distribution to Probe Intramolecular Interactions in Gas-Phase Phosphopeptides and Enhance Proteomics Analyses": Scripts for data analysis and generation of results and intermediate plots: file_description.txt

Provided scripts

pairwise_correlation.py - investigating reproducibility of the CSD measurements across laboratories and instruments

extract_CSD_msms.py - extracting CSDs from a single MS/MS (MSGF+ & Percolator) experiment

K562_merging.py - assembling complete K562 dataset from invidual files extracted using extract script

K562_exploration.py - investigaing CSD properties on K562 dataset

extract_CSD_phospho.py - extracting CSDs from a single msms.txt MaxQuant file (and several corresponding mass spectrometer files); support verification against standard library

phospho_merging.py - assembling complete phosphopeptides dataset from invidual files extracted using extract script

phospho_exploration.py - investigating CSD properties of phosphopeptides and their non-phosphorylated counterparts

digest_exploration.py - investigating CSD properties of the complete human proteome digest

CCS_Z.py - analysis of CCS and charge state dimemsions of phosphopeptides

ML_models.py - comparing differnt ML predictors for CSD

CSD_Learning_Phospho_cos.ipynb - training of deep learning predictor with several data splits

PercolatorCSD.py - adding CSD dimension to Percolator input

shotgun.py - using CSD in shotgun context
