## Supplementary material for "Exploiting Charge State Distribution to Probe Intramolecular Interactions in Gas-Phase Phosphopeptides and Enhance Proteomics Analyses": Scripts for data analysis and generation of results and intermediate plots: K562_exploration.py

```
# -*- coding: utf-8 -*-
"""
Investigating CSDs properties
Need:
    peptide-level CSD table

Output:
    Plots and graphs

Usage:
    plain script (not to be executed from command line)

@author: Vladimir
"""

import re
import pandas as pd
import numpy as np
import matplotlib.pyplot as plt
from pyteomics.proforma import ProForma
from pyteomics.mass import Composition, calculate_mass
from functools import reduce
from collections import Counter, defaultdict
from sklearn.metrics.pairwise import cosine_similarity
from sklearn.cluster import AgglomerativeClustering
from statsmodels.nonparametric.smoothers_lowess import lowess

def proforma_to_composition(proforma_string):
    p = ProForma.parse(proforma_string)
    mods = []
    sequence = []

    for aa, mod in p.sequence:
        sequence.append(aa)

        if not mod is None:
            for m in mod:
                mods.append(m.composition)

    if not p.n_term is None:
        for m in p.n_term:
            mods.append(m.composition)

    if not p.c_term is None:
        for m in p.c_term:
            mods.append(m.composition)

    composition = Composition(sequence=''.join(sequence)) + reduce(lambda a,b: a + b, mods, Composition())
    return ''.join([f'{k}{composition[k]}' for k in sorted(composition.keys())])

def cosine(data):
    if data.shape[0] > 1:
        splits = 1
        intermediate = []
        while (splits < 10):
            try:
                borders = np.round(np.linspace(0, data.shape[0], splits + 1), 0).astype(int)
                for i in range(1, splits + 1):
                    for j in range(i, splits + 1):
                        if i == j:
                            intermediate.append(cosine_similarity(data.loc[:, 'z=1':'z=6'].values[borders[i - 1]:borders[i], :])[np.triu_indices(borders[i] - borders[i - 1], 1)].mean())
                        else:
                            intermediate.append(cosine_similarity(data.loc[:, 'z=1':'z=6'].values[borders[i - 1]:borders[i], :],
                                                                  data.loc[:, 'z=1':'z=6'].values[borders[j - 1]:borders[j], :]).mean())

                return np.mean(intermediate)
            except MemoryError:
                print(f'Cannot calculate cosine cross-cimilarity for {data.shape[0]} elements with {splits} splits')
                splits += 1

        raise MemoryError(f'Cannot calculate cosine cross-cimilarity for {data.shape[0]} elements with {splits} splits')

    else:
        return np.nan

def cluster_size(data, metric='cosine', linkage='average', threshold=0.05):
    if data.shape[0] > 1:
        AC = AgglomerativeClustering(n_clusters=None, affinity=metric, linkage=linkage, distance_threshold=threshold)
        AC.fit(data.loc[:, 'z=1':'z=6'])
        return Counter(Counter(AC.labels_).values())
    else:
        return {1: 1}

def count_letters(sequence, relative=False):
    counts = Counter(sequence)
    if relative:
        length = len(sequence)
        return {f'rel_{k}': counts[k]/length for k in sorted(counts.keys())}
    else:
        return ''.join([f'{k}{counts[k]}' for k in sorted(counts.keys())])

def max_z_distance_g(frame):#the largest difference in average charge for unique sequences in a group
    gframe = frame.groupby('AA_composition').mean()
    if gframe.shape[0] > 1:
        return max_z_distance_i(gframe)
    else:
        return np.nan

def max_cos_distance_g(frame):#the smallest cosine similarity for unique sequences in a group
    gframe = frame.groupby('AA_composition').mean()
    if gframe.shape[0] > 1:
        return max_cos_distance_i(gframe)
    else:
        return np.nan

def max_z_distance_i(frame):#the largest difference in average charge for a group
    return np.max(frame.loc[:, 'charge'].values.reshape(-1, 1) - frame.loc[:, 'charge'].values.reshape(1, -1))

def max_cos_distance_i(frame):#the smallest cosine similarity for a group
    return cosine_similarity(frame.loc[:, 'z=1':'z=6']).min()

#load K562 dataset
peptides = pd.read_csv('D:/RawData/CSD/PXD024364/positive_peptides.csv')

#add composition information
peptides['sequence'] = [re.sub(r'\[.+?\]|\-', '', seq) for seq in peptides['ProForma']]
peptides['AA_composition'] = peptides['sequence'].apply(count_letters)
peptides['M'] = peptides['ProForma'].apply(lambda s: ProForma.parse(s).mass)
peptides['composition'] = peptides['ProForma'].apply(proforma_to_composition)

#save extra data for later
peptides.to_csv('D:/RawData/CSD/PXD024364/positive_peptides_more.csv', index=False)

#some renaming
peptides.rename(columns={z: f'z={z}' for z in '123456'}, inplace=True)
peptides.rename(columns={'av_charge': 'charge'}, inplace=True)

#major charge plot
plt.hist(peptides.loc[:, 'z=1':'z=6'].max(axis=1), bins=np.linspace(0.5, 1.0, 21),
         cumulative=True, histtype='step', density=True)
plt.xlim((0.5, 1))
plt.xlabel('Relative abundance of the major charge state')
plt.ylabel('Proportion of peptides')
plt.savefig('D:/RawData/CSD/pub/plots/K562/K562_major_charge.svg')

#binning with average charge
for bin_width in [1e-5, 5e-6, 2e-6]:
    for rt_width in [0.5, 1, 2]:
        for z_width in [0.1, 0.2, 0.4]:
            if not f'mass_group_{bin_width*1e6:.0f}' in peptides.columns:
                min_mass = peptides['M'].min() * (1 - bin_width / 2)
                peptides[f'mass_group_{bin_width*1e6:.0f}'] = np.floor((np.log(peptides['M']) - np.log(min_mass)) / np.log(1 + bin_width)).astype(int)

            if not f'rt_group_{rt_width}' in peptides.columns:
                min_rt = peptides['rt'].min() - rt_width / 2
                peptides[f'rt_group_{rt_width}'] = np.floor((peptides['rt'] - min_rt) / rt_width).astype(int)

            if not f'z_group_{z_width}' in peptides.columns:
                min_z = peptides['charge'].min() - z_width / 2
                peptides[f'z_group_{z_width}'] = np.floor((peptides['charge'] - min_z) / z_width).astype(int)

            mass_groups = Counter(peptides.groupby(f'mass_group_{bin_width*1e6:.0f}').size())
            mass_rt_groups = Counter(peptides.groupby([f'mass_group_{bin_width*1e6:.0f}', f'rt_group_{rt_width}']).size())
            mass_z_groups = Counter(peptides.groupby([f'mass_group_{bin_width*1e6:.0f}', f'z_group_{z_width}']).size())
            mass_rt_z_groups = Counter(peptides.groupby([f'mass_group_{bin_width*1e6:.0f}', f'rt_group_{rt_width}', f'z_group_{z_width}']).size())

            plt.bar(np.array(list(mass_groups.keys())) - 0.3,
                    np.array(list(mass_groups.values())) * np.array(list(mass_groups.keys())) / peptides.shape[0],
                    width=0.2, label='Mass')
            plt.bar(np.array(list(mass_z_groups.keys())) - 0.1,
                    np.array(list(mass_z_groups.values())) * np.array(list(mass_z_groups.keys())) / peptides.shape[0],
                    width=0.2, label='Mass+Z')
            plt.bar(np.array(list(mass_rt_groups.keys())) + 0.1,
                    np.array(list(mass_rt_groups.values())) * np.array(list(mass_rt_groups.keys())) / peptides.shape[0],
                    width=0.2, label='Mass+RT')
            plt.bar(np.array(list(mass_rt_z_groups.keys())) + 0.3,
                    np.array(list(mass_rt_z_groups.values())) * np.array(list(mass_rt_z_groups.keys())) / peptides.shape[0],
                    width=0.2, label='Mass+RT+Z')

            plt.xticks(np.arange(1, max(mass_groups.keys())))
            plt.legend()
            plt.xlabel('Number of peptides')
            plt.ylabel('Frequency')
            plt.xlim((0.3, 20.7))
            plt.ylim(top=1.0)
            plt.title(f'Mass bin: {bin_width*1e6:.0f}ppm; RT bin: {rt_width}min; Z bin: {z_width}')
            plt.savefig(f'D:/RawData/CSD/pub/plots/K562/K562_discrimination_{bin_width*1e6:.0f}ppm_{rt_width}min_{z_width}z.svg')
            plt.close()

#binning with cosine similarity
for bin_width in [1e-5, 5e-6, 2e-6]:
    for rt_width in [0.5, 1, 2]:
        for max_cos in [0.05, 0.025]:
            if not f'mass_group_{bin_width*1e6:.0f}' in peptides.columns:
                min_mass = peptides['M'].min() * (1 - bin_width / 2)
                peptides[f'mass_group_{bin_width*1e6:.0f}'] = np.floor((np.log(peptides['M']) - np.log(min_mass)) / np.log(1 + bin_width)).astype(int)

            if not f'rt_group_{rt_width}' in peptides.columns:
                min_rt = peptides['rt'].min() - rt_width / 2
                peptides[f'rt_group_{rt_width}'] = np.floor((peptides['rt'] - min_rt) / rt_width).astype(int)

            mass_groups = Counter(peptides.groupby(f'mass_group_{bin_width*1e6:.0f}').size())
            mass_rt_groups = Counter(peptides.groupby([f'mass_group_{bin_width*1e6:.0f}', f'rt_group_{rt_width}']).size())

            mass_z_count = defaultdict(int)
            for element in peptides.groupby([f'mass_group_{bin_width*1e6:.0f}']).apply(cluster_size, threshold=max_cos).tolist():
                for k, v in element.items():
                    mass_z_count[k] += v

            mass_rt_z_count = defaultdict(int)
            for element in peptides.groupby([f'mass_group_{bin_width*1e6:.0f}', f'rt_group_{rt_width}']).apply(cluster_size, threshold=max_cos).tolist():
                for k, v in element.items():
                    mass_rt_z_count[k] += v

            plt.bar(np.array(list(mass_groups.keys())) - 0.3,
                    np.array(list(mass_groups.values())) * np.array(list(mass_groups.keys())) / peptides.shape[0],
                    width=0.2, label='Mass')
            plt.bar(np.array(list(mass_z_count.keys())) - 0.1,
                    np.array(list(mass_z_count.values())) * np.array(list(mass_z_count.keys())) / peptides.shape[0],
                    width=0.2, label='Mass+Z')
            plt.bar(np.array(list(mass_rt_groups.keys())) + 0.1,
                    np.array(list(mass_rt_groups.values())) * np.array(list(mass_rt_groups.keys())) / peptides.shape[0],
                    width=0.2, label='Mass+RT')
            plt.bar(np.array(list(mass_rt_z_count.keys())) + 0.3,
                    np.array(list(mass_rt_z_count.values())) * np.array(list(mass_rt_z_count.keys())) / peptides.shape[0],
                    width=0.2, label='Mass+RT+Z')

            plt.xticks(np.arange(1, max(mass_groups.keys())))
            plt.legend()
            plt.xlabel('Number of peptides')
            plt.ylabel('Frequency')
            plt.xlim((0.3, 20.7))
            plt.ylim(top=1.0)
            plt.title(f'Mass bin: {bin_width*1e6:.0f}ppm; RT bin: {rt_width}min; Z bin: {max_cos}cos')
            plt.savefig(f'D:/RawData/CSD/pub/plots/K562/K562_discrimination_{bin_width*1e6:.0f}ppm_{rt_width}min_{max_cos}cos.svg')
            plt.close()

#cosine similarity of isomeric and isoelemental peptides
#preparing random shuffling of peptides
pep_index = list(peptides.index)
np.random.shuffle(pep_index)

#cosine correlation of isomeric peptides
isom_cos = peptides.groupby(['AA_composition', 'composition']).apply(cosine)#both AA content and elemental composition to account for modifications
plt.hist(isom_cos, bins=np.linspace(0, 1, 51), histtype='step', label='Sequence isomers', density=True)
plt.axvline(np.nanmean(isom_cos), color='blue')
plt.text(np.nanmean(isom_cos) - 0.005, plt.ylim()[1] * 0.95, f'Mean:\n{np.nanmean(isom_cos):.4f}', color='blue', ha='right', va='top')

#cosine correlation of isoelemental peptides
iso_cos = peptides.groupby(['composition']).apply(cosine)
plt.hist(iso_cos, bins=np.linspace(0, 1, 51), histtype='step', label='Constitutional isomers', density=True)
plt.axvline(np.nanmean(iso_cos), color='orange')
plt.text(np.nanmean(iso_cos) - 0.005, plt.ylim()[1] * 0.95, f'Mean:\n{np.nanmean(iso_cos):.4f}', color='orange', ha='right', va='top')

#cosine similarity of 25K random peptides in dataset
cos_vector = cosine_similarity(peptides.loc[pep_index[:25000], 'z=1':'z=6'].values,
                               peptides.loc[pep_index[25000:50000], 'z=1':'z=6'].values)[np.eye(25000).astype(bool)]
plt.hist(cos_vector, bins=np.linspace(0, 1, 51), histtype='step', label='Random pairs', density=True)
plt.axvline(np.nanmean(cos_vector), color='green')
plt.text(np.nanmean(cos_vector) - 0.005, plt.ylim()[1] * 0.95, f'Mean:\n{np.nanmean(cos_vector):.4f}', color='green', ha='right', va='top')

plt.legend(loc='upper left')
plt.xlabel('Cosine similarity')
plt.ylabel('Density')
plt.savefig('D:/RawData/CSD/pub/plots/K562/K562_cosine_iso_classes.svg')
plt.close()

#cosine similarity for peptides in the same mass or RT groups
for bin_width in [1e-5, 5e-6, 2e-6]:
    for rt_width in [0.5, 1, 2]:
        min_mass = peptides['M'].min() * (1 - bin_width / 2)
        peptides['mass_group'] = np.floor((np.log(peptides['M']) - np.log(min_mass)) / np.log(1 + bin_width)).astype(int)

        min_rt = peptides['rt'].min() - rt_width / 2
        peptides['rt_group'] = np.floor((peptides['rt'] - min_rt) / rt_width).astype(int)

        #similarity of peptides of peptides with similar masses
        mass_cos = peptides.groupby(['mass_group']).apply(cosine)
        plt.hist(mass_cos, bins=np.linspace(0, 1, 51), histtype='step', label='Mass', density=True)
        plt.axvline(np.nanmean(mass_cos), color='blue')

        #similarity of peptides with similar RT
        rt_cos = peptides.groupby(['rt_group']).apply(cosine)
        plt.hist(rt_cos, bins=np.linspace(0, 1, 51), histtype='step', label='RT', density=True)
        plt.axvline(np.nanmean(rt_cos), color='orange')

        #similarity of peptides with similar Mass and RT
        mass_rt_cos = peptides.groupby(['mass_group', 'rt_group']).apply(cosine)
        plt.hist(mass_rt_cos, bins=np.linspace(0, 1, 51), histtype='step', label='Mass+RT', density=True)
        plt.axvline(np.nanmean(mass_rt_cos), color='green')

        plt.legend(loc='upper left')
        plt.xlabel('Cosine similarity')
        plt.ylabel('Density')
        plt.savefig(f'D:/RawData/CSD/pub/plots/K562/K562_cosine_discrimination_{bin_width*1e6:.0f}ppm_{rt_width}min.svg')
        plt.close()

# charge shifts for isomeric (same AA) and isoelemental (same formula) peptides
isomer_peptides = peptides.groupby(['AA_composition', 'composition']).filter(lambda f: f.shape[0] > 1)
isoel_peptides = peptides.groupby('composition').filter(lambda f: f.shape[0] > 1)

#isomeric peptides
group_size = Counter(isomer_peptides.groupby('AA_composition').size())
plt.bar(group_size.keys(), group_size.values(), align='center')
plt.xticks(sorted(group_size.keys()), sorted(group_size.keys()))
plt.text(plt.xlim()[1] * 0.95, plt.ylim()[1] * 0.95,
         f'Total groups: {isomer_peptides.groupby("AA_composition").size().shape[0]}',
         ha='right', va='top')
plt.xlabel('Group size')
plt.ylabel('Nr cases')
plt.title('Size of isomeric (same AA) peptide groups')
plt.savefig('D:/RawData/CSD/pub/plots/K562/K562_isomeric_group_size.svg')

#distances in average charge and cosine
z_dist = isomer_peptides.groupby(['AA_composition', 'composition']).apply(max_z_distance_i)
cos_dist = isomer_peptides.groupby(['AA_composition', 'composition']).apply(max_cos_distance_i)

plt.hist(z_dist, bins=100)
plt.axvline(np.mean(z_dist), ls='--', color='navy')
plt.xlim(left=-.02)
plt.text(np.mean(z_dist) * 1.1 , plt.ylim()[1] * 0.97, f'Mean: {np.mean(z_dist):.4f}', ha='left', va='top')
plt.axvspan(0.1, plt.xlim()[1], color='green', alpha=0.2)
plt.text((plt.xlim()[1] + 0.1) / 2 , plt.ylim()[1] * 0.5, f'Difference > 0.1: {np.mean(z_dist > 0.1) * 100:.1f}%', ha='center', va='center')
plt.xlabel('absoulute average charge difference')
plt.title('Average charge difference [max of isomeric peptides]')
plt.savefig('D:/RawData/CSD/pub/plots/K562/K562_isomeric_z_distance.svg')
plt.close()

plt.hist(cos_dist, bins=100)
plt.axvline(np.mean(cos_dist), ls='--', color='navy')
plt.xlim(-.01, 1.01)
plt.text(np.mean(cos_dist) * 0.9 , plt.ylim()[1] * 0.97, f'Mean: {np.mean(cos_dist):.4f}', ha='right', va='top')
plt.axvspan(plt.xlim()[0], 0.95, color='green', alpha=0.2)
plt.text((plt.xlim()[1] + plt.xlim()[0]) / 2 , plt.ylim()[1] * 0.5, f'Cosine similarity < 0.95: {np.mean(cos_dist < 0.95) * 100:.1f}%', ha='center', va='center')
plt.xlabel('cosine similarity')
plt.title('Cosine similarity [max of isomeric peptides]')
plt.savefig('D:/RawData/CSD/pub/plots/K562/K562_isomeric_cos_distance.svg')

#cosine similarity vs mass
plt.scatter([calculate_mass(formula=comp) for comp in cos_dist.index.get_level_values('composition')],
            cos_dist, s=2)
plt.xlabel('Peptide mass')
plt.ylabel('Cosine similarity')
plt.savefig('D:/RawData/CSD/pub/plots/K562/K562_isomeric_cos_distnace_vs_mass.png', dpi=150)

#isoelemental peptides
group_size = Counter(isoel_peptides.groupby('composition').size())
plt.bar(group_size.keys(), group_size.values(), align='center')
plt.xticks(sorted(group_size.keys()), sorted(group_size.keys()))
plt.xlim(left=1, right=31)
plt.text(plt.xlim()[1] * 0.95, plt.ylim()[1] * 0.95,
         f'Total groups: {isoel_peptides.groupby("composition").size().shape[0]}',
         ha='right', va='top')
plt.xlabel('Group size')
plt.ylabel('Nr cases')
plt.title('Size of isoelemental (same CHNOS) peptide groups')
plt.savefig('D:/RawData/CSD/pub/plots/K562/K562_isoelement_group_size.svg')

#distances in average charge and cosine
z_dist = isoel_peptides.groupby('composition').apply(max_z_distance_i)
cos_dist = isoel_peptides.groupby('composition').apply(max_cos_distance_i)

plt.hist(z_dist, bins=100)
plt.axvline(np.mean(z_dist), ls='--', color='navy')
plt.xlim(left=-.02)
plt.text(np.mean(z_dist) * 1.1 , plt.ylim()[1] * 0.97, f'Mean: {np.mean(z_dist):.4f}', ha='left', va='top')
plt.axvspan(0.1, plt.xlim()[1], color='green', alpha=0.2)
plt.text((plt.xlim()[1] + 0.1) / 2 , plt.ylim()[1] * 0.5, f'Difference > 0.1: {np.mean(z_dist > 0.1) * 100:.1f}%', ha='center', va='center')
plt.xlabel('absoulute average charge difference')
plt.title('Average charge difference [max of isoelemental peptides]')
plt.savefig('D:/RawData/CSD/pub/plots/K562/K562_isoelement_z_distanсe.svg')
plt.close()

plt.hist(cos_dist, bins=100)
plt.axvline(np.mean(cos_dist), ls='--', color='navy')
plt.xlim(-.01, 1.01)
plt.text(np.mean(cos_dist) * 0.9 , plt.ylim()[1] * 0.97, f'Mean: {np.mean(cos_dist):.4f}', ha='right', va='top')
plt.axvspan(plt.xlim()[0], 0.95, color='green', alpha=0.2)
plt.text((plt.xlim()[1] + plt.xlim()[0]) / 2 , plt.ylim()[1] * 0.5, f'Cosine similarity < 0.95: {np.mean(cos_dist < 0.95) * 100:.1f}%', ha='center', va='center')
plt.xlabel('cosine similarity')
plt.title('Cosine similarity [max of isoelemental peptides]')
plt.savefig('D:/RawData/CSD/pub/plots/K562/K562_isoelement_cos_distance.svg')

#distances in average charge and cosine for unique AA compositions
z_dist = isoel_peptides.groupby('composition').apply(max_z_distance_g)
z_dist = z_dist[np.isfinite(z_dist)]
cos_dist = isoel_peptides.groupby('composition').apply(max_cos_distance_g)
cos_dist = cos_dist[np.isfinite(cos_dist)]

plt.hist(z_dist, bins=100)
plt.axvline(np.mean(z_dist), ls='--', color='navy')
plt.xlim(left=-.02)
plt.text(np.mean(z_dist) * 1.1 , plt.ylim()[1] * 0.97, f'Mean: {np.mean(z_dist):.4f}', ha='left', va='top')
plt.axvspan(0.1, plt.xlim()[1], color='green', alpha=0.2)
plt.text((plt.xlim()[1] + 0.1) / 2 , plt.ylim()[1] * 0.5, f'Difference > 0.1: {np.mean(z_dist > 0.1) * 100:.1f}%', ha='center', va='center')
plt.xlabel('absoulute average charge difference')
plt.title('Average charge difference [max of strictly isoelemental peptides]')
plt.savefig('D:/RawData/CSD/pub/plots/K562/K562_strict_isoelement_z_distanсe.svg')
plt.close()

plt.hist(cos_dist, bins=100)
plt.axvline(np.mean(cos_dist), ls='--', color='navy')
plt.xlim(-.01, 1.01)
plt.text(np.mean(cos_dist) * 0.9 , plt.ylim()[1] * 0.97, f'Mean: {np.mean(cos_dist):.4f}', ha='right', va='top')
plt.axvspan(plt.xlim()[0], 0.95, color='green', alpha=0.2)
plt.text((plt.xlim()[1] + plt.xlim()[0]) / 2 , plt.ylim()[1] * 0.5, f'Cosine similarity < 0.95: {np.mean(cos_dist < 0.95) * 100:.1f}%', ha='center', va='center')
plt.xlabel('cosine similarity')
plt.title('Cosine similarity [max of strictly isoelemental peptides]')
plt.savefig('D:/RawData/CSD/pub/plots/K562/K562_strict_isoelement_cos_distance.svg')

#overcharging/undercharging
#count basic sites
peptides['n_basic'] = peptides['ProForma'].str.count('[KHR]') - peptides['ProForma'].str.count('\[Acetyl\]-') + 1

#averge charge vs mass
plt.figure(figsize=(7, 7))
for n_basic, pep_group in peptides.groupby('n_basic'):
    if n_basic <= 6:
        plt.subplot(3, 2, n_basic)
        under = pep_group['charge'] - n_basic < -0.05
        over = pep_group['charge'] - n_basic > 0.05
        plt.scatter(pep_group.loc[under, 'M'], pep_group.loc[under, 'charge'], alpha=0.5, s=2)
        plt.scatter(pep_group.loc[over, 'M'], pep_group.loc[over, 'charge'], alpha=0.5, s=2)
        plt.scatter(pep_group.loc[~over & ~under, 'M'], pep_group.loc[~over & ~under, 'charge'], alpha=0.5, s=2)
        plt.axhline(n_basic, ls='--', color='darkgreen', lw=1)
        lowessFit = lowess(pep_group['charge'].values, pep_group['M'].values, is_sorted=False, frac=0.05, it=1)
        plt.plot(lowessFit[:, 0], lowessFit[:, 1], lw=1, color='tab:gray', label='Lowess fit')
        plt.ylim((0.5, 6.5))
        plt.yticks([1, 2, 3, 4, 5, 6], [1, 2, 3, 4, 5, 6])
        plt.xlabel('Peptide mass')
        plt.ylabel('Observed charge')
        plt.title(f'{n_basic} basic sites')

plt.tight_layout()
plt.savefig('D:/RawData/CSD/pub/plots/K562/K562_under_overcharging.png', dpi=150)
```
