## Supplementary material for "Exploiting Charge State Distribution to Probe Intramolecular Interactions in Gas-Phase Phosphopeptides and Enhance Proteomics Analyses": Scripts for data analysis and generation of results and intermediate plots: K562_merging.py

```
# -*- coding: utf-8 -*-
"""
Merging and combining CSDs extracted at the previous steps
Need:
    Raw CSD files

Output:
    Peptde-level CSDs table for K562 dataset
    Plots and graphs

Usage:
    plain script (not to be executed from command line)

@author: Vladimir
"""

import pandas as pd
import numpy as np
import matplotlib.pyplot as plt
from os import path, listdir
from sklearn.metrics.pairwise import cosine_similarity

def correlate(frame):
    data = frame.iloc[:, :6].copy().values
    nonzero = data[:, data.sum(axis=0) > 0]

    if nonzero.shape[0] > 1: # multiple entries
        cos = cosine_similarity(nonzero, nonzero)
        #removing outlier samples (i.e. keep the samples that have cosine similarity above 0.9 with other samples above the 75th percentile)
        correlated = (cos > 0.9).sum(axis=1) >= np.percentile((cos > 0.9).sum(axis=1), 75)
        if correlated.sum() > 1: #more than one well-correlated sample
            result = pd.DataFrame(data[correlated, :].mean(axis=0).reshape(1, -1), columns=frame.columns[:6])
            result['correlation'] = np.median(cosine_similarity(nonzero[correlated], nonzero[correlated])\
                                              [np.triu_indices(correlated.sum(), 1)])
            result['cor_length'] = correlated.sum()
            result['length'] = nonzero.shape[0]
        elif correlated.sum() == 1:#single well-correlated sample
            result = pd.DataFrame(data[correlated, :].mean(axis=0).reshape(1, -1), columns=frame.columns[:6])
            result['correlation'] = 1.0
            result['cor_length'] = 1
            result['length'] = nonzero.shape[0]
        else:#no well-correlated samples
            result = pd.DataFrame(data.mean(axis=0).reshape(1, -1), columns=frame.columns[:6])
            result['correlation'] = 0
            result['cor_length'] = 0
            result['length'] = nonzero.shape[0]

    else: #single entry
        result = pd.DataFrame(data.mean(axis=0).reshape(1, -1), columns=frame.columns[:6])
        result['correlation'] = 1.0
        result['cor_length'] = 1
        result['length'] = 1

    return result


#merging data from individual files
files = []
#LysC
for filename in listdir('D:/RawData/CSD/PXD024364/K562_LysC_HCD'):
    if filename.endswith('csd.csv'):
        label = filename[:-8]
        data = pd.read_csv(path.join('D:/RawData/CSD/PXD024364/K562_LysC_HCD', filename)).fillna(0)
        data = data[~np.all(data.loc[:, '1':'6'] == 0, axis=1)].copy()
        files.append(data)

#trypsin
for filename in listdir('D:/RawData/CSD/PXD024364/K562_Try_HCD'):
    if filename.endswith('csd.csv'):
        label = filename[:-8]
        data = pd.read_csv(path.join('D:/RawData/CSD/PXD024364/K562_Try_HCD', filename)).fillna(0)
        data = data[~np.all(data.loc[:, '1':'6'] == 0, axis=1)].copy()
        files.append(data)

#laoding data
positive = pd.concat(files, axis=0, ignore_index=True)

positive.fillna(0, inplace=True)

#remove empty Proformas
positive.drop(positive.index[positive['ProForma'] == '0'], inplace=True)
#drop empty CSD
positive.drop(positive.index[positive.loc[:, '1':'6'].sum(axis=1) == 0], inplace=True)
#drop axiliary columns
positive.drop(['rt_start', 'rt_end', 'pepid'], axis=1, inplace=True)

#save total intensity and rt separately
positive_abundance = positive.loc[:, '1':'6'].sum(axis=1)
positive_abundance = pd.concat([positive_abundance, positive['rt'], positive['ProForma']], axis=1)
positive_abundance.rename({0: 'abundance'}, axis=1, inplace=True)

#normalize CSD distributions
positive.loc[:, '1':'6'] = positive.loc[:, '1':'6'].values / positive.loc[:, '1':'6'].sum(axis=1).values.reshape(-1, 1)

#remove duplicate rows
unique = ~positive.duplicated()
positive = positive[unique].reset_index(drop=True)
positive_abundance = positive_abundance[unique].reset_index(drop=True)

#calculate average charge
positive['z_av'] = (positive.loc[:, '1':'6'] * np.arange(1, 7).reshape(1, -1)).sum(axis=1)

#spread of average charge
def spread(data):
    if data.shape[0] > 1:
        return data['z_av'].std()
    else:
        return np.nan

z_vect = positive.groupby('ProForma').apply(spread)
z_vect = z_vect[np.isfinite(z_vect)].values

plt.hist(z_vect, bins=np.linspace(0, 0.4, 100))
perc95 = np.percentile(z_vect, 95)
plt.axvline(perc95, ls=':', color='r')
plt.text(perc95 * 1.05, plt.ylim()[1] * 0.95, f'95%-le = {perc95:.2f}', color='r')
under01 = (z_vect < 0.1).mean()
plt.axvline(0.1, ls=':', color='g')
plt.text(0.1 * 1.05, plt.ylim()[1] * 0.5, f'Under 0.1 - {under01 * 100:.2f}%', color='g')
plt.title('Standard deviation of average charge between the same peptides')
plt.savefig('D:/RawData/CSD/pub/plots/K562/K562_z_av_spread.svg')

#group by Proforma and calculate correlation
pos_peptides = positive.groupby('ProForma').apply(correlate)

#drop extra index level
pos_peptides.index = pos_peptides.index.droplevel(1)

#add max observed intensity per peptide
pos_peptides[['abundance', 'rt']] = positive_abundance.groupby('ProForma').agg({'abundance': np.max, 'rt': np.mean})

#correlation histogram
pos_peptides['correlation'].hist(bins=100)
plt.title('Cosine similarity between multiple CSDs of the same peptide sequence')
plt.savefig('D:/RawData/CSD/pub/plots/K562/K562_csd_cosine_correlation.svg')

#correlation histogram closeup
pos_peptides['correlation'].hist(bins=200, figsize=(4, 4))
plt.xlim(0.89, 1.01)
plt.title('Cosine similarity between multiple CSDs of the same peptide sequence (inset)')
plt.savefig('D:/RawData/CSD/pub/plots/K562/K562_csd_cosine_correlation_inset.svg')

#select peptides with decent correlation, i.e. group correlation > 0.9 excluding the cases where only 1 peptide have good correlation out of several
valid = np.logical_and(pos_peptides['correlation'] > 0.9,
                       np.logical_not(np.logical_and(pos_peptides['cor_length'] == 1, pos_peptides['length'] > 1)))

#number of observed charge states
pos_peptides['nr_charges'] = (pos_peptides.loc[:, '1':'6'] > 0).sum(axis=1)

#nr_charges distribution data
with open('D:/RawData/CSD/pub/plots/K562/positive_peptides.stats', 'w') as stats:
    stats.write('{}\n\n{}'.format(
        pos_peptides.loc[valid, :].groupby('nr_charges').count()['1'],
        (pos_peptides.loc[valid, :].groupby('nr_charges').count() / valid.sum())['1']
        )
    )

#recalulate average charge for filtered peptides
pos_peptides['av_charge'] = (pos_peptides.loc[:, '1':'6'].values * np.arange(1,7)).sum(axis=1)

#average charge histogram
pos_peptides.loc[valid, 'av_charge'].hist(bins=100, log=True)
plt.title('Average observed peptide charge')
plt.savefig('D:/RawData/CSD/pub/plots/K562/K562_average_charge_hist.svg')

pos_peptides.loc[valid, :].reset_index().to_csv('D:/RawData/CSD/PXD024364/positive_peptides.csv', index=False)

#log-abundance boxplot
pos_peptides['log_abundance'] = np.log10(pos_peptides['abundance'])
pos_peptides.boxplot(column='log_abundance', by='nr_charges')
plt.ylabel('$log_{10}(Abundance_{max})$')
plt.title('Number of observed charge states vs abundance')
plt.savefig('D:/RawData/CSD/pub/plots/K562/K562_log_abundance_vs_nr_charge.svg')
```
