## Supplementary material for "Exploiting Charge State Distribution to Probe Intramolecular Interactions in Gas-Phase Phosphopeptides and Enhance Proteomics Analyses": Scripts for data analysis and generation of results and intermediate plots: ML_models.py

```
# -*- coding: utf-8 -*-
"""
Comparing different ML predictors for CSD
Need:
    Peptide CSD dataset (regular & phospho) splitted into testing and trainig sets

Output:
    Plots and graphs

Usage:
    plain script (not to be executed from command line)

@author: Vladimir
"""

import numpy as np
import pandas as pd
import matplotlib.pyplot as plt
from scipy.stats import spearmanr
from keras import models
from keras.metrics import cosine_similarity
from seaborn import kdeplot

psi_to_single_ptm = {'[Acetyl]-': 'B',
                     '[Carbamidomethyl]': '',
                     'M[Oxidation]': 'J',
                     'S[Phosphorylation]': 'X',
                     'T[Phosphorylation]': 'O',
                     'Y[Phosphorylation]': 'U'}

def reshapeOneHot(X):
    X = np.dstack(X)
    X = np.swapaxes(X, 1, 2)
    X = np.swapaxes(X, 0, 1)
    return X

def get_single_ptm_code(psi_sequence):
    sequence = psi_sequence
    for ptm in psi_to_single_ptm:
        sequence = sequence.replace(ptm, psi_to_single_ptm[ptm])
    return sequence

def one_hot_encode_peptide(psi_sequence, MAX_LENGTH = 40):
    peptide = get_single_ptm_code(psi_sequence)
    if len(peptide) > MAX_LENGTH:
        print('Peptide length is larger than maximal length of ', str(MAX_LENGTH))
        return ['']
    else:
        AA_vocabulary = 'KRPTNAQVSGILCMJHFYWEDBXOU'#B: acetyl; J: oxidized Met; O: PhosphoT; X: PhosphoS; U: PhosphoY

        one_hot_peptide = np.zeros((len(peptide), len(AA_vocabulary)))

        for j in range(0, len(peptide)):
            try:
                aa = peptide[j]
                one_hot_peptide[j, AA_vocabulary.index(aa)] = 1
            except:
                print(f'WARN: "{aa}" is not in vocabulary; it will be skipped')

        no_front_paddings = int((MAX_LENGTH - len(peptide))/2)
        peptide_front_paddings = np.zeros((no_front_paddings, one_hot_peptide.shape[1]))

        no_back_paddings = MAX_LENGTH - len(peptide) - no_front_paddings
        peptide_back_paddings = np.zeros((no_back_paddings, one_hot_peptide.shape[1]))

        full_one_hot_peptide = np.vstack((peptide_front_paddings, one_hot_peptide, peptide_back_paddings))

        return peptide, full_one_hot_peptide

def sign(x):
    return '+' if x >= 0 else '-'

def plot_charge_correlation(Y, Y_pred):
    for z in range(6):
        plt.subplot(2, 3, z + 1)
        plt.scatter(Y[:, z], Y_pred[:, z], s=2, alpha=0.5)
        fit = np.poly1d(np.polyfit(Y[:, z], Y_pred[:, z], deg=1))
        plt.plot(plt.xlim(), fit(plt.xlim()), 'r--')
        R = spearmanr(Y[:, z], Y_pred[:, z]).statistic
        plt.xlim((0, 1))
        plt.text(0.95, plt.ylim()[1] * 0.25, f'R_s = {R:.3f}', va='center', ha='right')
        plt.text(0.05, plt.ylim()[1] * 0.75,
                 '{:.3f}x {} {:.3f}'.format(fit.coef[0], sign(fit.coef[1]),
                                            fit.coef[1]), va='center', ha='left', color='r')
        plt.xlabel('Real')
        plt.ylabel('Predicted')
        plt.title(f'Z = {z + 1}')

    plt.tight_layout()

#encoding peptides
data = pd.read_csv('D:/RawData/CSD/splits_phospho_cos/peptides+phospho_1.0_train.csv', na_filter=False, low_memory=False)
train_X = reshapeOneHot(data['proforma'].apply(lambda s: one_hot_encode_peptide(s)[1]).values).sum(axis=1) #flattening one-hot-encode
train_Y = data.loc[:, '1':'6'].values
print(f'Train size {data.shape[0]}')

data = pd.read_csv('D:/RawData/CSD/splits_phospho_cos/peptides+phospho_1.0_test.csv', na_filter=False, low_memory=False)
test_X = reshapeOneHot(data['proforma'].apply(lambda s: one_hot_encode_peptide(s)[1]).values).sum(axis=1) #flattening one-hot-encode
test_Y = data.loc[:, '1':'6'].values
print(f'Test size {data.shape[0]}')

with open('D:\RawData\CSD\pub\plots\ML\errors.txt', 'w') as fout:
    #Simple linear regression
    from sklearn.linear_model import LinearRegression

    model = LinearRegression()

    model.fit(train_X, train_Y)

    train_error = np.mean(cosine_similarity(train_Y, model.predict(train_X)))
    test_error = np.mean(cosine_similarity(test_Y, model.predict(test_X)))

    plt.figure('kdeplot')
    kdeplot(cosine_similarity(test_Y, model.predict(test_X)), label='linear regression', color='tab:blue')
    plt.axvline(test_error, ls='--', color='tab:blue')

    print(f'Linear regression\nError on the train set: {train_error:.6f}\nError on the test set: {test_error:.6f}')
    fout.write(f'Linear regression\nError on the train set: {train_error:.6f}\nError on the test set: {test_error:.6f}\n')

    plt.figure()
    plot_charge_correlation(test_Y, model.predict(test_X))
    plt.savefig('D:\RawData\CSD\pub\plots\ML\linear_regression.png', dpi=150)
    plt.close()

    #KNN
    from sklearn.neighbors import KNeighborsRegressor

    model = KNeighborsRegressor(7)

    model.fit(train_X, train_Y)

    train_error = np.mean(cosine_similarity(train_Y, model.predict(train_X)))
    test_error = np.mean(cosine_similarity(test_Y, model.predict(test_X)))

    plt.figure('kdeplot')
    kdeplot(cosine_similarity(test_Y, model.predict(test_X)), label='KNN regression', color='tab:green')
    plt.axvline(test_error, ls='--', color='tab:green')

    print(f'KNN regression\nError on the train set: {train_error:.6f}\nError on the test set: {test_error:.6f}')
    fout.write(f'KNN regression\nError on the train set: {train_error:.6f}\nError on the test set: {test_error:.6f}\n')

    plt.figure()
    plot_charge_correlation(test_Y, model.predict(test_X))
    plt.savefig('D:\RawData\CSD\pub\plots\ML\KNN_regression.png', dpi=150)
    plt.close()

    #Decision tree
    from sklearn.tree import DecisionTreeRegressor

    model = DecisionTreeRegressor()

    model.fit(train_X, train_Y)

    train_error = np.mean(cosine_similarity(train_Y, model.predict(train_X)))
    test_error = np.mean(cosine_similarity(test_Y, model.predict(test_X)))

    plt.figure('kdeplot')
    kdeplot(cosine_similarity(test_Y, model.predict(test_X)), label='decision tree regression', color='tab:orange')
    plt.axvline(test_error, ls='--', color='tab:orange')

    print(f'Decision tree regression\nError on the train set: {train_error:.6f}\nError on the test set: {test_error:.6f}')
    fout.write(f'Decision tree regression\nError on the train set: {train_error:.6f}\nError on the test set: {test_error:.6f}\n')

    plt.figure()
    plot_charge_correlation(test_Y, model.predict(test_X))
    plt.savefig('D:\RawData\CSD\pub\plots\ML\decision_tree_regression.png', dpi=150)
    plt.close()

    #GAM for each charge state
    from pygam import GAM
    from sklearn.multioutput import MultiOutputRegressor

    model = GAM(lam=0.01)
    mulmodel = MultiOutputRegressor(model)

    mulmodel.fit(train_X, train_Y)

    train_error = np.mean(cosine_similarity(train_Y, mulmodel.predict(train_X)))
    test_error = np.mean(cosine_similarity(test_Y, mulmodel.predict(test_X)))

    plt.figure('kdeplot')
    kdeplot(cosine_similarity(test_Y, mulmodel.predict(test_X)), label='MultiGAM', color='tab:red')
    plt.axvline(test_error, ls='--', color='tab:red')

    print(f'GAM regression\nError on the train set: {train_error:.6f}\nError on the test set: {test_error:.6f}')
    fout.write(f'GAM regression\nError on the train set: {train_error:.6f}\nError on the test set: {test_error:.6f}\n')

    plt.figure()
    plot_charge_correlation(test_Y, mulmodel.predict(test_X))
    plt.savefig('D:\RawData\CSD\pub\plots\ML\GAM_multi_regression.png', dpi=150)
    plt.close()

    #Trained LSTM model
    #reload complete data
    data = pd.read_csv('D:/RawData/CSD/splits_phospho_cos/peptides+phospho_1.0_train.csv', na_filter=False, low_memory=False)
    train_X = reshapeOneHot(data['proforma'].apply(lambda s: one_hot_encode_peptide(s)[1]).values)
    train_Y = data.loc[:, '1':'6'].values
    print(f'Train size {data.shape[0]}')

    data = pd.read_csv('D:/RawData/CSD/splits_phospho_cos/peptides+phospho_1.0_test.csv', na_filter=False, low_memory=False)
    test_X = reshapeOneHot(data['proforma'].apply(lambda s: one_hot_encode_peptide(s)[1]).values)
    test_Y = data.loc[:, '1':'6'].values
    print(f'Test size {data.shape[0]}')

    model = models.load_model('D:\RawData\CSD\splits_phospho_cos\CSD_model_LCMSMS_1.0.h5')
    model.compile()

    train_error = np.mean(cosine_similarity(train_Y.astype('float32'), model.predict(train_X, batch_size=2048)))
    test_error = np.mean(cosine_similarity(test_Y.astype('float32'), model.predict(test_X, batch_size=2048)))

    plt.figure('kdeplot')
    kdeplot(cosine_similarity(test_Y.astype('float32'), model.predict(test_X, batch_size=2048)), label='LSTM', color='tab:purple')
    plt.axvline(test_error, ls='--', color='tab:purple')
    plt.legend()
    plt.title('Cosine similarity distribution on the test set')
    plt.savefig('D:\RawData\CSD\pub\plots\ML\models_error_kde.svg')
    plt.close()

    print(f'Deep learning\nError on the train set: {train_error:.6f}\nError on the test set: {test_error:.6f}')
    fout.write(f'Deep learning\nError on the train set: {train_error:.6f}\nError on the test set: {test_error:.6f}\n')

    plt.figure()
    plot_charge_correlation(test_Y, model.predict(test_X, batch_size=2048))
    plt.savefig('D:\RawData\CSD\pub\plots\ML\LSTM_regression.png', dpi=150)
    plt.close()
```
