## Supplementary material for "Exploiting Charge State Distribution to Probe Intramolecular Interactions in Gas-Phase Phosphopeptides and Enhance Proteomics Analyses": Scripts for data analysis and generation of results and intermediate plots: pairwise_correlation.py

```
# -*- coding: utf-8 -*-
"""
Investigating reproducibility of CSD measurement across laboratories and instruments
Regular peptides (nano, micro, tims) and phosphopeptides
Need:
    Extracted CSD tables for all raw files
    Sample description table (as provided in the supplementary files)

Output:
    Correlation heatmaps

Usage:
    plain script (not to be executed from command line)

@author: Vladimir
"""

import pandas as pd
import numpy as np
import seaborn as sb
import matplotlib.pyplot as plt
from sklearn.metrics.pairwise import cosine_similarity
from os import listdir, path

###### pairwise correlations for regular peptides #######
#file info table for regular runs
fileinfo = pd.read_csv('D:/RawData/CSD/selected_filenames.csv')
fileinfo.sort_values(['Gradient length (min)', 'Laboratory', 'Instrument', 'MS1 resolution', 'MS2 resolution', 'MS1 AGC', 'Sample load', 'Replicate'],
                      inplace=True)
fileinfo['MS1 AGC'] = fileinfo['MS1 AGC'].apply(lambda x: '{}e6'.format(int(x/1e6)))

#loading CSD tables
files = {}

for parent in ['D:/RawData/CSD/regular_csd', 'D:/RawData/Micro', 'D:/RawData/CSD/timsTOF/csds']:
    for filename in listdir(parent):
        if filename.endswith('csd.csv'):
            label = filename[:-8]
            data = pd.read_csv(path.join(parent, filename)).fillna(0)
            data = data[~np.all(data.loc[:, '1':'6'] == 0, axis=1)].copy()
            data.loc[:, '1':'6'] = data.loc[:, '1':'6'] / data.loc[:, '1':'6'].sum(axis=1).values.reshape(-1, 1)
            files[label] = data

labels = list(fileinfo['filename'])

#building correlation matrix
results = np.array([np.nan] * len(labels) ** 2).reshape(len(labels), -1)

for i in range(len(labels)):
    for j in range(i + 1, len(labels)):
        csd_two = files[labels[i]].merge(files[labels[j]], on='ProForma',
                                          how='inner', suffixes=('_1', '_2')).drop(['pepid_1', 'pepid_2'], axis=1)

        if csd_two.shape[0] > 0:
            results[j, i] = cosine_similarity(csd_two.loc[:, '1_1':'6_1'].values, csd_two.loc[:, '1_2':'6_2'].values)[np.eye(csd_two.shape[0]).astype(bool)].mean()

#setting labels
labels = fileinfo.loc[:, 'Laboratory':].apply(lambda r: '_'.join(r.astype(str)), axis=1)
results_frame = pd.DataFrame(results, index=labels, columns=labels)

plt.figure(figsize=(20,20))
sb.heatmap(results_frame, mask=np.triu(np.ones_like(results_frame)), square=True, linewidths=0.5, cbar_kws={'shrink': 0.8}, vmax=1)
plt.savefig('D:/RawData/CSD/pub/plots/run_correlation/run_correlation_cos_similarity.svg')

#building representative examples diferent cosine correlations
#chosing two random files and selecting common peptides
label1, label2 = np.random.choice(labels, 2)
csd_cross = files[label1].merge(files[label2], on='ProForma',
                                how='inner', suffixes=('_1', '_2')).drop(['pepid_1', 'pepid_2'], axis=1)
csd_cross['cos'] = cosine_similarity(csd_cross.loc[:, '1_1':'6_1'].values, csd_cross.loc[:, '1_2':'6_2'].values)[np.eye(csd_cross.shape[0]).astype(bool)]
csd_cross['ProForma'] = csd_cross['ProForma'].str.replace('[Carbamidomethyl]', '')

#plotting distribution around several key regions
for n, (target_cos, target_tol) in enumerate(zip([0.999, 0.97, 0.95, 0.90, 0.80, 0.50],
                                                 [0.0009, 0.002, 0.002, 0.002, 0.05, 0.1])):
    plt.subplot(2, 3, n + 1)
    selected = np.random.choice(csd_cross.index[np.abs(csd_cross['cos'] - target_cos) < target_tol], 1)[0]
    plt.bar(np.arange(1, 7) - 0.05, csd_cross.loc[selected, '1_1':'6_1'].values, width=0.1, align='center')
    plt.bar(np.arange(1, 7) + 0.05, csd_cross.loc[selected, '1_2':'6_2'].values, width=0.1, align='center')
    plt.text(6, plt.ylim()[1] * 0.9, 'Cosine: {:.3f}'.format(csd_cross.at[selected, 'cos']),
             va='center', ha='right')
    plt.xticks(np.arange(1, 7), [f'{x}+' for x in np.arange(1, 7)])
    plt.ylabel('Rel. abundance')
    plt.title(csd_cross.at[selected, 'ProForma'])

plt.tight_layout()
plt.savefig('D:/RawData/CSD/pub/plots/run_correlation/cos_similarity_examples.svg')

#adding comparison with microflow experiments
microfileinfo = pd.read_csv('D:/RawData/Micro/selected_filenames.csv')
microfileinfo.sort_values(['Gradient length (min)', 'Laboratory', 'Instrument', 'MS1 resolution', 'MS2 resolution', 'MS1 AGC', 'Sample load'],
                      inplace=True)
microfileinfo['MS1 AGC'] = microfileinfo['MS1 AGC'].apply(lambda x: '{}e6'.format(int(x/1e6)))

x_labels = list(fileinfo['filename'])
y_labels = list(microfileinfo['filename'])

#building correlation matrix
results = np.array([np.nan] * (len(x_labels) + len(y_labels)) * len(y_labels)).reshape((len(x_labels) + len(y_labels)),
                                                                                      len(y_labels))

#micro vs nanoflow
for i in range(len(x_labels)):
    for j in range(len(y_labels)):
        csd_two = files[x_labels[i]].merge(files[y_labels[j]], on='ProForma',
                                          how='inner', suffixes=('_1', '_2')).drop(['pepid_1', 'pepid_2'], axis=1)

        if csd_two.shape[0] > 0:
            results[i, j] = cosine_similarity(csd_two.loc[:, '1_1':'6_1'].values, csd_two.loc[:, '1_2':'6_2'].values)[np.eye(csd_two.shape[0]).astype(bool)].mean()

#micro vs micro
for i in range(len(y_labels)):
    for j in range(i + 1, len(y_labels)):
        csd_two = files[y_labels[i]].merge(files[y_labels[j]], on='ProForma',
                                          how='inner', suffixes=('_1', '_2')).drop(['pepid_1', 'pepid_2'], axis=1)

        if csd_two.shape[0] > 0:
            results[i + len(x_labels), j] = cosine_similarity(csd_two.loc[:, '1_1':'6_1'].values, csd_two.loc[:, '1_2':'6_2'].values)[np.eye(csd_two.shape[0]).astype(bool)].mean()

#setting labels
x_labels = fileinfo.loc[:, 'Laboratory':].apply(lambda r: '_'.join(r.astype(str)), axis=1)
y_labels = microfileinfo.loc[:, 'Laboratory':].apply(lambda r: '_'.join(r.astype(str)), axis=1)
results_frame = pd.DataFrame(results, index=np.concatenate([x_labels, y_labels]), columns=y_labels)

plt.figure(figsize=(20,6))
sb.heatmap(results_frame.T, square=True, linewidths=0.5, cbar_kws={'shrink': 0.8}, vmax=1)
plt.savefig('D:/RawData/CSD/pub/plots/run_correlation/run_correlation_micro_cos_similarity.svg')

#comparison with timsTOF data
timsfileinfo = pd.read_csv('D:/RawData/CSD/timsTOF/csds/selected_filenames.csv', dtype=object)
timsfileinfo.sort_values(['Sample'], inplace=True)

#converting CSD tables to common format
for filename in timsfileinfo['filename']:
    csd_table = files[filename]
    csd_table.rename({'Unnamed: 0': 'ProForma'}, axis=1, inplace=True)
    csd_table['ProForma'] = csd_table['ProForma'].str.replace('camC', 'C[Carbamidomethyl]', regex=False)
    csd_table['ProForma'] = csd_table['ProForma'].str.replace('oxM', 'M[Oxidation]', regex=False)
    csd_table['ProForma'] = csd_table['ProForma'].str.replace('ac-', '[Acetyl]-', regex=False)

x_labels = list(fileinfo['filename'])
y_labels = list(timsfileinfo['filename'])

results = np.array([np.nan] * (len(x_labels) + len(y_labels)) * len(y_labels)).reshape((len(x_labels) + len(y_labels)), len(y_labels))

#timsTOF vs Orbitrap
for i in range(len(x_labels)):
    for j in range(len(y_labels)):
        csd_two = files[x_labels[i]].merge(files[y_labels[j]], on='ProForma',
                                          how='inner', suffixes=('_1', '_2')).drop(['pepid'], axis=1)

        if csd_two.shape[0] > 0:
            results[i, j] = cosine_similarity(csd_two.loc[:, '1_1':'6_1'].values, csd_two.loc[:, '1_2':'6_2'].values)[np.eye(csd_two.shape[0]).astype(bool)].mean()

#timsTOF vs timsTOF
for i in range(len(y_labels)):
    for j in range(i + 1, len(y_labels)):
        csd_two = files[y_labels[i]].merge(files[y_labels[j]], on='ProForma',
                                          how='inner', suffixes=('_1', '_2'))

        if csd_two.shape[0] > 0:
            results[i + len(x_labels), j] = cosine_similarity(csd_two.loc[:, '1_1':'6_1'].values, csd_two.loc[:, '1_2':'6_2'].values)[np.eye(csd_two.shape[0]).astype(bool)].mean()


#setting labels
x_labels = fileinfo.loc[:, 'Laboratory':].apply(lambda r: '_'.join(r.astype(str)), axis=1)
y_labels = timsfileinfo.loc[:, 'Laboratory':].apply(lambda r: '_'.join(r.astype(str)), axis=1)
results_frame = pd.DataFrame(results, index=np.concatenate([x_labels, y_labels]), columns=y_labels)

plt.figure(figsize=(20,6))
sb.heatmap(results_frame.T, square=True, linewidths=0.5, cbar_kws={'shrink': 0.8}, vmax=1)
plt.savefig('D:/RawData/CSD/pub/plots/run_correlation/run_correlation_tims_cos_similarity.svg')


###### pairwise correlations for phospho peptides #######
#file info table for phospho runs
fileinfo = pd.read_csv('D:/RawData/PhosphoCSD/selected_filenames.csv')
fileinfo.sort_values(['Gradient length (min)', 'Laboratory', 'Instrument', 'MS1 resolution',  'Replicate'],
                      inplace=True)

#loading CSD tables
files = {}

parent = 'D:/RawData/PhosphoCSD'
for filename in listdir(parent):
    if filename.endswith('csd.csv'):
        label = filename[:-8]
        data = pd.read_csv(path.join(parent, filename)).fillna(0)
        data = data[~np.all(data.loc[:, '1':'6'] == 0, axis=1)].copy()
        data.loc[:, '1':'6'] = data.loc[:, '1':'6'] / data.loc[:, '1':'6'].sum(axis=1).values.reshape(-1, 1)
        files[label] = data

labels = list(fileinfo['filename'])

#building correlation matrix
results = np.array([np.nan] * len(labels) ** 2).reshape(len(labels), -1)

for i in range(len(labels)):
    for j in range(i + 1, len(labels)):
        csd_two = files[labels[i]].merge(files[labels[j]], on='proforma',
                                          how='inner', suffixes=('_1', '_2')).drop(['pepid_1', 'pepid_2'], axis=1)

        if csd_two.shape[0] > 0:
            results[j, i] = cosine_similarity(csd_two.loc[:, '1_1':'6_1'].values, csd_two.loc[:, '1_2':'6_2'].values)[np.eye(csd_two.shape[0]).astype(bool)].mean()

#setting labels
labels = fileinfo.loc[:, 'Laboratory':].apply(lambda r: '_'.join(r.astype(str)), axis=1)
results_frame = pd.DataFrame(results, index=labels, columns=labels)

plt.figure(figsize=(20,20))
sb.heatmap(results_frame, mask=np.triu(np.ones_like(results_frame)), square=True, linewidths=0.5, cbar_kws={'shrink': 0.8}, vmax=1)
plt.savefig('D:/RawData/CSD/pub/plots/run_correlation/run_correlation_cos_similarity.svg')
```
