## Supplementary material for "Exploiting Charge State Distribution to Probe Intramolecular Interactions in Gas-Phase Phosphopeptides and Enhance Proteomics Analyses": Scripts for data analysis and generation of results and intermediate plots: PercolatorCSD.py

```
# -*- coding: utf-8 -*-
"""
Adding CSD dimension to Percolator input
Need:
    Regular Percolator .pin file
    corresponding mzIdentML file for every raw file
    corresponding biosaur2 features file for every raw file
    FASTA file (needed for Percolator to perform protein inference)
    ThermoRawFileParser (has to be in PATH and as `thermoparser`) and corresponding raw files
    Trained CSD model

    All files has to share the base name (before the file extension) and location

Output:
    Updates .pin file with CSD-features and runs Percolator on it
    new .pin file will be saved as {base name}_csd.pin

Usage:
    <script_name> <path to pin file> <path to FASTA file>

@author: Vladimir
"""
import logging
import json
import pandas as pd
import numpy as np
from pyteomics.mzid import MzIdentML
from pyteomics.proforma import ProForma
from lxml import etree
from collections import Counter
from subprocess import check_output, CalledProcessError
from keras import models

logging.basicConfig(format='%(asctime)s %(message)s', datefmt='%H:%M:%S', level=logging.INFO)

mzTol = 7e-6
PROTON = 1.007276
index_max = 2

psi_to_single_ptm = {'[Acetyl]-': 'B',
                     '[Carbamidomethyl]': '',
                     'M[Oxidation]': 'J',
                     'S[Phospho]': 'X',
                     'T[Phospho]': 'O',
                     'Y[Phospho]': 'U'}

def read_pin(filepath):
    with open(filepath, 'r') as input_file:
        headers = input_file.readline().strip().split('\t')
        num_headers = len(headers)
        data = []
        for line in input_file.readlines():
            fields = line.strip().split('\t', num_headers - 1)
            data.append({headers[n]: value for (n, value) in enumerate(fields)})

    return pd.DataFrame(data)

def write_pin(dataframe, subheader, filepath):
    with open(filepath, 'w') as output_file:
        output_file.write('\t'.join(dataframe.columns) + '\n')
        output_file.write(subheader + '\n')
        for row in dataframe.values:
            output_file.write('\t'.join([str(field) for field in row]) + '\n')

def mapModifications(psm):
    peptide = [''] + list(psm['PeptideSequence']) + ['']
    mods = psm.get('Modification', [])

    if len(mods) > 0:
        for mod in mods:
            loc = int(mod['location'])
            modtype = mod.get('name', mod['monoisotopicMassDelta'])
            peptide[loc] += f'[{modtype}]'

        if peptide[0] != '':
            peptide[0] = peptide[0] + '-'
        if peptide[-1] != '':
            peptide[-1] = '-' + peptide[-1]

    return ''.join(peptide)

def parse_psm(psm):
    proteins = [ev['accession'] for ev in psm['PeptideEvidenceRef']]
    proforma = mapModifications(psm)
    isDecoy = np.all([protein.startswith('DECOY_') for protein in proteins])
    charge = psm['chargeState']
    num_matched = psm['NumMatchedMainIons']
    return {'proforma': proforma, 'decoy': isDecoy, 'charge': charge, 'num_matched': num_matched}

def matchPSM(psm, features, faims_mode=False):
    if faims_mode:
        match_index = (features['rtStart'] < psm['rt']).astype(int) +\
                    (features['rtEnd'] > psm['rt']).astype(int) +\
                    (features['charge'] == psm['charge']).astype(int) +\
                    (np.abs((features['massCalib'] - psm['mass'])/psm['mass']) < mzTol).astype(int) +\
                    (features['FAIMS'] == psm['CV']).astype(int) == 5
    else:
        match_index = (features['rtStart'] < psm['rt']).astype(int) +\
                    (features['rtEnd'] > psm['rt']).astype(int) +\
                    (features['charge'] == psm['charge']).astype(int) +\
                    (np.abs((features['massCalib'] - psm['mass'])/psm['mass']) < mzTol).astype(int) == 4

    if sum(match_index) > 1:
        logging.warning(f'Warning: multiple PSM match\n{features.loc[match_index]}')
        features.loc[np.logical_and(match_index, features['match'] == ''), 'match'] += f';{psm["id"]}'
        logging.warning(f'Final snip\n{features.loc[match_index]}')
        return True

    elif sum(match_index) == 1:
        features.loc[match_index, 'match'] += f';{psm["id"]}'
        return True

    return False

def groupPFMs(features, faims_mode=False):
    features['pepid'] = -1
    pepid = 1

    for index in features.index[features['match_count'] > 0]:
        if features.at[index, 'pepid'] == -1: #not assigned so far
                mass = features.at[index, 'massCalib']
                start = features.at[index, 'rtStart']
                end = features.at[index, 'rtEnd']
                cv = features.at[index, 'FAIMS']

                #select features based on similar mass and overlapping elution
                if faims_mode:
                    join_candidates = features[(np.abs(features['massCalib'] - mass) / mass < mzTol).astype(int) +\
                                            (features['rtStart'] <= end).astype(int) +\
                                            (features['rtEnd'] >= start).astype(int) +\
                                            (features['FAIMS'] == cv).astype(int) == 4]
                else:
                    join_candidates = features[(np.abs(features['massCalib'] - mass) / mass < mzTol).astype(int) +\
                                            (features['rtStart'] <= end).astype(int) +\
                                            (features['rtEnd'] >= start).astype(int) == 3]

                #require apex indices to be not more then index_max away
                group_ind = join_candidates.index[np.abs(join_candidates['scanApex'] - join_candidates.at[index, 'scanApex']) <= index_max]

                occupied = features.loc[group_ind, 'pepid'] != -1
                if np.any(occupied):
                    occupied_pepid = np.unique(features.loc[group_ind[occupied], "pepid"])
                    if len(occupied_pepid) > 1:
                        logging.warning(f'WARNING Mixed peptide group observed at {index}; candidate IDS: {group_ind}; PEPIDs: {occupied_pepid}')
                        logging.info('Finding group with the smallest mass difference:')
                        best_delta = mass * mzTol
                        best_pepid = -1
                        for i in occupied_pepid:
                            delta = np.abs(mass - np.mean(features.loc[features['pepid'] == i, 'massCalib']))
                            if delta < best_delta:
                                best_pepid = i
                                best_delta = delta

                        logging.info(f'BestID: {best_pepid}; BestDelta: {best_delta}')
                        features.loc[group_ind[~occupied], 'pepid'] = best_pepid

                    else:
                        features.loc[group_ind, 'pepid'] = occupied_pepid[0]

                else:
                    features.loc[group_ind, 'pepid'] = pepid
                    pepid += 1

    features.drop(features.index[features['pepid'] == -1], axis=0, inplace=True)

    logging.info('Solving multiple assignements')
    #try solving multiple assignment cases (in quite a shitty way)
    #also geting some stats
    counts = {'Cases': 0,
              'Joinable': 0,
              'Disjoint': 0,
              'Multicase': 0}

    for gid, group in features.groupby(['pepid', 'charge']):
        if group.shape[0] > 1:
            counts['Cases'] += 1
            z = gid[1]
            candidates = features[features['pepid'] == gid[0]].copy()
            #more than one double assignment per peptide
            if not np.all(candidates[candidates['charge'] != z].groupby('charge').count()['massCalib'] == 1):
                counts['Multicase'] += 1
                logging.warning(f'WARNING multiple cases per peptide:\n{candidates}')

            scans = group['mono_hills_scan_lists'].str.slice(1, -1).apply(np.fromstring, sep=',')
            sorted_index = (scans.apply(np.min).sort_values()).index
            #feature can be merged - all scan numbers are consequtive with max 1 gap
            if np.all(scans[sorted_index[1:]].apply(np.min).values - scans[sorted_index[0:-1]].apply(np.max).values <= 2):
                counts['Joinable'] += 1
                remove = group.index.drop(group.index[np.argmax(group['intensityApex'])])
                #mark as useless
                features.loc[remove, 'charge'] = 0
            else:
                counts['Disjoint'] += 1
                logging.warning(f'WARNING Disjoint features:\n{candidates}')
                #mark all as suspicious
                features.loc[group.index, 'charge'] = 0

    #drop useless features
    features.drop(features.index[features['charge'] == 0], inplace=True)
    logging.info(f'Solving done:\n{counts}')

def extract_feature_properties(group, faims_mode=False):
    group['sequence'] = group['match'].str.replace(r'\/\d+?', '', regex=True)
    sequence_vect = group.loc[group['match'].astype(str) != '', 'sequence'].str.split(';', expand=True).values.flatten()
    sequence_vect = np.unique(sequence_vect[sequence_vect != None])
    if len(sequence_vect) > 0:
        if faims_mode:
            feat_vect = [group.agg({'massCalib': np.mean, 'rtApex': np.mean, 'rtStart': np.min, 'rtEnd': np.max, 'FAIMS': np.mean}).values.reshape(1, -1) for _ in sequence_vect]
            frame = pd.DataFrame(np.concatenate(feat_vect), columns=['mass', 'rtApex', 'rtStart', 'rtEnd', 'FAIMS'], index=sequence_vect)
        else:
            feat_vect = [group.agg({'massCalib': np.mean, 'rtApex': np.mean, 'rtStart': np.min, 'rtEnd': np.max}).values.reshape(1, -1) for _ in sequence_vect]
            frame = pd.DataFrame(np.concatenate(feat_vect), columns=['mass', 'rtApex', 'rtStart', 'rtEnd'], index=sequence_vect)
        return frame

def AUC(x, y):
    return (0.5 * (y[1:] + y[:-1]) * (x[1:] - x[:-1])).sum()

def aggregate_csd(g):
    result = {k: v for k,v in zip(g['z'], g['area'])}
    result['id'] = g['id'].iloc[0]
    result['rt_start'] = g['rt_start'].iloc[0]
    result['rt_end'] = g['rt_end'].iloc[0]
    result['rt'] = g['rt'].iloc[0]
    result['pepid'] = g['pepid'].iloc[0]
    return result

def expand_charges(row, min_charge, max_charge, mode):
    charge = np.arange(min_charge, max_charge + 1).reshape(-1, 1)
    mz = row['mass']/charge + PROTON * mode
    index = [f'{row["pepid"]}_{z[0]}' for z in charge]
    result = pd.DataFrame(mz, columns=['mz'], index=index)
    result['z'] = charge
    result['rt_start'] = row['rtStart']
    result['rt_end'] = row['rtEnd']
    result['rt'] = row['rtApex']
    result['id'] = row['id']
    if 'FAIMS' in row.keys():
        result['scan_filter'] = f'ms cv={row["FAIMS"]}'

    return result

def reshapeOneHot(X):
    X = np.dstack(X)
    X = np.swapaxes(X, 1, 2)
    X = np.swapaxes(X, 0, 1)
    return X

def get_single_ptm_code(psi_sequence):
    sequence = psi_sequence
    for ptm in psi_to_single_ptm:
        sequence = sequence.replace(ptm, psi_to_single_ptm[ptm])
    return sequence

def one_hot_encode_peptide(psi_sequence, MAX_LENGTH = 40):
    peptide = get_single_ptm_code(psi_sequence)
    if len(peptide) > MAX_LENGTH:
        print('Peptide length is larger than maximal length of ', str(MAX_LENGTH))
        return ['', None]
    else:
        AA_vocabulary = 'KRPTNAQVSGILCMJHFYWEDBXOU'#B: acetyl; O: Carbamyl; J: oxidized Met; X:pyro_glu

        one_hot_peptide = np.zeros((len(peptide), len(AA_vocabulary)))

        for j in range(0, len(peptide)):
            try:
                aa = peptide[j]
                one_hot_peptide[j, AA_vocabulary.index(aa)] = 1
            except:
                print(f'WARN: "{aa}" is not in vocabulary; it will be skipped')

        no_front_paddings = int((MAX_LENGTH - len(peptide))/2)
        peptide_front_paddings = np.zeros((no_front_paddings, one_hot_peptide.shape[1]))

        no_back_paddings = MAX_LENGTH - len(peptide) - no_front_paddings
        peptide_back_paddings = np.zeros((no_back_paddings, one_hot_peptide.shape[1]))

        full_one_hot_peptide = np.vstack((peptide_front_paddings, one_hot_peptide, peptide_back_paddings))

        return peptide, full_one_hot_peptide


def process(filepath, fastapath):
    logging.info(f'Using FASTA: {fastapath}')
    #reading Percolator PIN
    logging.info(f'Reading .pin file from {filepath}')
    #reading Percolator PIN
    pin_data = read_pin(filepath)

    logging.info(f'{pin_data.shape[0]} entries read')

    subheader = '\t'.join(pin_data.loc[0, :].astype(str)[:-2])

    pin_data.drop(0, inplace=True)

    pin_data['id'] = pin_data['SpecId'].str.extract('(SII_\d+?_\d+?)_')

    logging.info('Building SII to SIR cache')
    #making mapping of SII id to SIR id
    idFile = etree.parse(filepath.replace('.pin', '.mzid'))

    sii2sir = {sii.get('id'): sii.getparent().get('id')
               for sii in idFile.xpath("//a:DataCollection//a:SpectrumIdentificationItem",
                                       namespaces = {"a": idFile.getroot().nsmap[None]})}

    logging.info(f'Reading PSMs from {filepath.replace(".pin", ".mzid")}')
    #reading PIN an mapping data
    psms = MzIdentML(filepath.replace('.pin', '.mzid'))

    psm_table = pd.DataFrame(pin_data['id'].apply(lambda p: parse_psm(psms.get_by_id(p))).tolist(), index=pin_data['id'])
    psm_table['rt'] = [psms.get_by_id(sii2sir[i])['scan start time'].real for i in psm_table.index]
    psm_table['mass'] = psm_table['proforma'].apply(lambda s: ProForma.parse(s).mass)
    psm_table['id'] = psm_table.index

    #reading features
    logging.info('Reading features')
    features = pd.read_csv(filepath.replace('.pin', '.features.tsv'), sep='\t')
    logging.info(f'Read {features.shape[0]} features')

    logging.info(f'Mapping {psm_table.shape[0]} PSMs..')
    features['match'] = ''
    matched_psms = psm_table.apply(matchPSM, axis=1, features=features)
    logging.info(f'Matched PSMs: {matched_psms.sum()} ({100 * matched_psms.mean():.1f}%)')
    features['match'] = features['match'].apply(lambda s: ';'.join(set(s.strip(';').split(';'))))
    features['match_count'] = features['match'].str.count(';') + 1
    features.loc[features['match'] == '', 'match_count'] = 0
    logging.info(f'Found PFMs: {Counter(features["match_count"])}')

    logging.info('Grouping PFMs')
    groupPFMs(features)

    if features.shape[0] == 0:
        logging.warning('no features left after grouping, the rest will be skipped')
        logging.info('Processing completed\n')
        return

    #merging PFM groups
    peptides = features.groupby('pepid').apply(extract_feature_properties)
    peptides.index = peptides.index.droplevel(0)
    peptides.index.names = ['id']
    peptides.reset_index(inplace=True)
    peptides['pepid'] = np.arange(peptides.shape[0])
    logging.info(f'Total peptides: {peptides.shape[0]}')

    #making mz-table
    logging.info('Building XIC table')
    mz_table = pd.concat(peptides.apply(expand_charges, axis=1, args=(1, 6, 1)).tolist())
    mzmin = np.floor(features['mz'].min())
    mzmax = np.ceil(features['mz'].max())
    mz_table = mz_table.query('mz >= @mzmin & mz <= @mzmax').copy()

    #writing JSON input for TRFP
    mz_table['tolerance'] = 5
    mz_table['tolerance_unit'] = 'ppm'
    mz_table['comment'] = mz_table.index
    with open(filepath.replace('.pin', '_perc_mz.json'), 'w') as out:
        json.dump(mz_table.drop(['z', 'id', 'rt'], axis=1).apply(lambda row: {k:v for k,v in zip(row.index, row)}, axis=1).tolist(), out)

    trfp_params = ['thermoparser', 'xic', '-i', f'{filepath.replace(".pin", ".raw")}', '-j', f'{filepath.replace(".pin", "_perc_mz.json")}']

    try:
        logging.info(f'Using TRFP to extract {mz_table.shape[0]} XICs')
        check_output(trfp_params, shell=True)
    except CalledProcessError as ex:
        logging.critical(f'TRFP execution failed\nOutput:\n{ex.output}')
        raise Exception('TRFP XIC extraction failed')

    #reading TRFP output
    with open(filepath.replace('.pin', '.json'), 'r') as xic_input:
        chromatograms = json.load(xic_input)

    #integrity check
    if not np.all(mz_table['comment'] == [cr['Meta']['Comment'] for cr in chromatograms['Content']]):
        logging.critical('Misalignment between XICs and mz_table!')
        raise Exception('Misalignment; processing failed')

    #calculating areas
    mz_table['area'] = [AUC(np.array(cr['RetentionTimes']), np.array(cr['Intensities'])) for cr in chromatograms['Content']]
    mz_table['pepid'] = mz_table['comment'].str.split('_', expand=True).iloc[:, 0]

    #calculating CSD table
    csd_table = pd.DataFrame(mz_table.groupby('pepid').apply(aggregate_csd).tolist())
    csd_table.rename(columns={k:v for k,v in zip(csd_table.columns, csd_table.columns.astype(str))}, inplace=True)
    csd_table = csd_table.reindex(columns=sorted(csd_table.columns))

    #summing duplicated CSDs
    csd_table = csd_table.groupby('id').sum().loc[:, '1':'6'].copy()

    #adding sequences
    csd_table['proforma'] = psm_table.reindex(csd_table.index)['proforma']

    #discard too long peptides
    csd_table = csd_table[csd_table['proforma'].apply(lambda s: one_hot_encode_peptide(s)[0] != '')].copy()

    #saving empty spots
    not_possible = np.isnan(csd_table.loc[:, '1':'6'].values)

    csd_table.fillna(0, inplace=True)
    csd_table.loc[:, '1':'6'] /= csd_table.loc[:, '1':'6'].sum(axis=1).values.reshape(-1, 1)
    csd_table.insert(6, 'z', (csd_table.loc[:, '1':'6'] * np.arange(1, 7).reshape(1, 6)).sum(axis=1))

    CSD_LCMSMS = models.load_model('D:/RawData/CSD/splits_phospho_cos/CSD_model_LCMSMS_1.0.h5', compile=False)
    CSD_LCMSMS.compile()

    csd_table = csd_table[csd_table['proforma'].apply(lambda s: one_hot_encode_peptide(s)[0] != '')].copy()
    X = reshapeOneHot(csd_table['proforma'].apply(lambda s: one_hot_encode_peptide(s)[1]).values)
    Y = CSD_LCMSMS.predict(X, batch_size=2048)
    Y[not_possible] = 0
    Y /= Y.sum(axis=1).reshape(-1, 1)
    z_pred = (Y * np.arange(1, 7).reshape(1, 6)).sum(axis=1)

    #error in CSD as z-score
    csd_err = (csd_table.loc[:, '1':'6'] - Y)
    csd_err = (csd_err - np.nanmean(csd_err, axis=0)) / np.nanstd(csd_err, axis=0)

    #error in Z as z-score
    z_err = (csd_table.loc[:, 'z'] - z_pred)
    z_err = (z_err - np.nanmean(z_err, axis=0)) / np.nanstd(z_err, axis=0)

    #merging and upscaling to psm_table and applying defaults
    csd_err['z'] = z_err
    complete_csd_err = csd_err.reindex(psm_table['id']).copy()

    #default z-scores for target and decoy
    complete_csd_err = pd.concat([complete_csd_err.loc[psm_table['decoy'], :].fillna(complete_csd_err.loc[psm_table['decoy'], :].mean(axis=0)),
                                  complete_csd_err.loc[~psm_table['decoy'], :].fillna(complete_csd_err.loc[~psm_table['decoy'], :].mean(axis=0))])

    pin_data[['1', '2', '3', '4', '5', '6', 'z']] = complete_csd_err.reindex(psm_table['id']).loc[:, '1':'z'].values
    pin_data['nMatch'] = psm_table.reindex(pin_data['id'])['num_matched'].values
    pin_data['cover'] = pin_data['nMatch'] / pin_data['PepLen'].astype(float)

    #reordering columns
    cols = list(pin_data.columns)
    cols.remove('Peptide')
    cols.remove('Proteins')
    cols.remove('id')
    cols.extend(['Peptide', 'Proteins'])

    #writing result
    write_pin(pin_data[cols], subheader, filepath.replace('.pin', '_csd.pin'))

    #running percolator
    perc_params = ['C:/Program Files/percolator-v3-05/bin/percolator.exe', '-Y', '-P', 'DECOY_', '-c', '-g',
                   '-f', fastapath, '-z', 'trypsin',
                   '-X', filepath.replace('.pin', '_csd.pout.xml'), '-w', filepath.replace('.pin', '_csd.weights'),
                   filepath.replace('.pin', '_csd.pin')]

    try:
        logging.info('Running Percolator...')
        check_output(perc_params, shell=True)
    except CalledProcessError as ex:
        logging.critical(f'Percolator execution failed\nOutput:\n{ex.output}')
        raise Exception('Percolator failed')

    logging.info(f'Processing of {filepath} completed')

if __name__ == '__main__':
    import sys
    process(sys.argv[1], sys.argv[2])
```
