## Supplementary material for "Exploiting Charge State Distribution to Probe Intramolecular Interactions in Gas-Phase Phosphopeptides and Enhance Proteomics Analyses": Scripts for data analysis and generation of results and intermediate plots: phospho_exploration.py

```
# -*- coding: utf-8 -*-
"""
Investigating CSDs properties of phosphopeptides and their counterparts from K562 dataset
Need:
    peptide-level CSD table for K562 and phospho set

Output:
    Plots and graphs

Usage:
    plain script (not to be executed from command line)

@author: Vladimir
"""
import re
import pandas as pd
import numpy as np
import matplotlib.pyplot as plt
from pyteomics.proforma import ProForma
from pyteomics.mass import Composition
from pyteomics.electrochem import gravy
from functools import reduce
from itertools import combinations
from collections import Counter
from sklearn.metrics.pairwise import cosine_similarity
from sklearn.cluster import AgglomerativeClustering
from statsmodels.nonparametric.smoothers_lowess import lowess
from seaborn import kdeplot

def proforma_to_composition(proforma_string):
    p = ProForma.parse(proforma_string)
    mods = []
    sequence = []

    for aa, mod in p.sequence:
        sequence.append(aa)

        if not mod is None:
            for m in mod:
                mods.append(m.composition)

    if not p.n_term is None:
        for m in p.n_term:
            mods.append(m.composition)

    if not p.c_term is None:
        for m in p.c_term:
            mods.append(m.composition)

    composition = Composition(sequence=''.join(sequence)) + reduce(lambda a,b: a + b, mods, Composition())
    return ''.join([f'{k}{composition[k]}' for k in sorted(composition.keys())])

def cosine(data):
    if data.shape[0] > 1:
        return cosine_similarity(data.loc[:, 'z=1':'z=6'].values)[np.triu_indices(data.shape[0], 1)].mean()
    else:
        return np.nan

def cluster_size(data, metric='cosine', linkage='average', threshold=0.05):
    if data.shape[0] > 1:
        AC = AgglomerativeClustering(n_clusters=None, affinity=metric, linkage=linkage, distance_threshold=threshold)
        AC.fit(data.loc[:, 'z=1':'z=6'])
        return Counter(Counter(AC.labels_).values())
    else:
        return {1: 1}

def count_letters(sequence, relative=False):
    counts = Counter(sequence)
    if relative:
        length = len(sequence)
        return {f'rel_{k}': counts[k]/length for k in sorted(counts.keys())}
    else:
        return ''.join([f'{k}{counts[k]}' for k in sorted(counts.keys())])

#largest difference in charge
def max_z_distance_i(frame):
    return np.max(frame.loc[:, 'charge'].values.reshape(-1, 1) - frame.loc[:, 'charge'].values.reshape(1, -1))

#smallet cosine similarity
def max_cos_distance_i(frame):
    return cosine_similarity(frame.loc[:, '1':'6']).min()

#extract sequence span upstream and downstream from phosphosite; pad with "Z" - non existing AA
def extract_padded_sequence(proforma, span):
    results = []
    seq = re.sub(r'\[.+?\]|\-', '', proforma)
    pep = ProForma.parse(proforma)
    for n, aa in enumerate(pep):
        if not aa[1] is None and np.any([mod.name == 'Phospho' for mod in aa[1]]):
            start = n
            end = n + 1
            results.append('Z' * (span - start) + seq[max(0, start - span):start] +\
                           seq[start] + seq[end:min(len(seq), end + span)] +\
                           'Z' * (span - len(seq) + end))

    return results

#charge shift by proforma
def get_z_shifts(frame):
    return pd.DataFrame({'proforma': frame.loc[frame['Phospho_count'] > 0, 'proforma'],
                         'shift': (frame.loc[frame['Phospho_count'] > 0, 'charge'].values - frame.loc[frame['Phospho_count'] == 0, 'charge'].values),
                         'charge': frame.loc[frame['Phospho_count'] == 0, 'charge'].values[0]})

#cosine similarity by proforma
def get_cos_sim(frame):
    return pd.DataFrame({'proforma': frame.loc[frame['Phospho_count'] > 0, 'proforma'],
                         'cos': cosine_similarity(frame.loc[frame['Phospho_count'] > 0, '1':'6'].values, frame.loc[frame['Phospho_count'] == 0, '1':'6'].values).flatten()})

#max charge difference between the groups of peptides with the same number of phosphosites
def max_z_sites(frame):
    groups = {n: g.loc[:, 'charge'].values for n, g in frame.groupby('Phospho_count')}
    return {'{}-{}'.format(*pair): np.abs(groups[pair[0]].reshape(-1,1) - groups[pair[1]].reshape(1, -1)).max() for pair in combinations(groups.keys(), 2)}

#min cosine similarity between the groups of peptides with the same number of phosphosites
def min_cos_sites(frame):
    groups = {n: g.loc[:, '1':'6'].values for n, g in frame.groupby('Phospho_count')}
    return {'{}-{}'.format(*pair): cosine_similarity(groups[pair[0]], groups[pair[1]]).min() for pair in combinations(groups.keys(), 2)}

#detect positions of phosphorylations
def get_phospho_pos(proforma_string):
    positions = []
    for n, (aa, mods) in enumerate(ProForma.parse(proforma_string).sequence):
        if not mods is None:
            for mod in mods:
                if mod.name == 'Phospho':
                    positions.append(n)

    return positions

#weighted standard deviation along axis
def weighted_std(x, w, a):
    average = np.average(x, weights=w, axis=a).reshape(-1, 1)
    return np.sqrt(np.average((x - average) ** 2, weights=w, axis=a))

#load phosphopeptide data
peptidesP = pd.read_csv('D:/RawData/PhosphoCSD/peptides.csv')
#load K562 data
peptidesK = pd.read_csv('D:/RawData/CSD/PXD024364/positive_peptides.csv').rename(columns={'ProForma': 'proforma'})
#merging into single dataset
peptides = pd.concat([peptidesK, peptidesP])

#adding compositional information
peptides.fillna(0, inplace=True)
peptides['sequence'] = [re.sub(r'\[.+?\]|\-', '', seq) for seq in peptides['proforma']]
peptides['charge'] = (peptides.loc[:, '1':'6'] * np.arange(1, 7).reshape(1, -1)).sum(axis=1)

peptides['AA_composition'] = peptides['sequence'].apply(count_letters)
peptides['M'] = peptides['proforma'].apply(lambda s: ProForma.parse(s).mass)
peptides['composition'] = peptides['proforma'].apply(proforma_to_composition)
peptides['Phospho_count'] = peptides['proforma'].str.count('\[Phosphorylation\]')

#saving for later
peptides.to_csv('D:/RawData/CSD/peptides+phospho_more.csv', index=False)

#dephosphorylated sequence
peptides['dephosphorylated'] = peptides['proforma'].str.replace('[Phosphorylation]', '', regex=False)

#find groups of peptides with the same dephosphorylated sequence and at least one plain and one phosphorylated peptide
phospho_paired = peptides.groupby('dephosphorylated').filter(lambda f: np.logical_and(np.any(f['Phospho_count'] > 0),
                                                                                      np.any(f['Phospho_count'] == 0)))

#group sizes
group_size = Counter(phospho_paired.groupby('dephosphorylated').size())
plt.bar(group_size.keys(), group_size.values(), align='center')
plt.xticks(sorted(group_size.keys()), sorted(group_size.keys()))
plt.text(plt.xlim()[1] * 0.95, plt.ylim()[1] * 0.95,
         f'Total groups: {phospho_paired.groupby("dephosphorylated").size().shape[0]}',
         ha='right', va='top')
plt.xlabel('Group size')
plt.ylabel('Nr cases')
plt.title('Size of paired phospho/nonphospho groups')
plt.savefig('D:/RawData/CSD/pub/plots/phospho/phospho_paired_group_size.svg')

#changes in average observed charge and cosine similarity
z_shifts = phospho_paired.groupby('dephosphorylated').apply(get_z_shifts)
z_shifts['Phospho_count'] = z_shifts['proforma'].str.count('\[Phosphorylation\]')
cos_sims = phospho_paired.groupby('dephosphorylated').apply(get_cos_sim)
cos_sims['Phospho_count'] = cos_sims['proforma'].str.count('\[Phosphorylation\]')

z_dist = z_shifts['shift'].values
bin_edges = np.linspace(z_dist.min(), z_dist.max(), 101)
bottoms = np.zeros(100)
for n_phos, shifts in z_shifts.groupby('Phospho_count'):
    bins, _ = np.histogram(shifts['shift'], bins=bin_edges)
    plt.bar((bin_edges[:-1] + bin_edges[1:]) / 2, bins, bin_edges[1] - bin_edges[0], bottoms, label=f'{n_phos}-Phospho')
    bottoms += bins
plt.xlim(left=z_dist.min() - 0.02, right=z_dist.max() + 0.02)
plt.axvline(np.mean(z_dist), ls='--', color='navy')
plt.text(np.mean(z_dist) * 1.1 , plt.ylim()[1] * 0.97, f'Mean: {np.mean(z_dist):.4f}', ha='right', va='top')
plt.axvspan(plt.xlim()[0], -0.1, color='green', alpha=0.2)
plt.axvspan(0.1, plt.xlim()[1], color='green', alpha=0.2)
plt.text((plt.xlim()[0] - 0.1) / 2 , plt.ylim()[1] * 0.5, f'Difference < -0.1: {np.mean(z_dist < -0.1) * 100:.1f}%', ha='center', va='center')
plt.text((plt.xlim()[1] + 0.1) / 2 , plt.ylim()[1] * 0.5, f'Difference > 0.1: {np.mean(z_dist > 0.1) * 100:.1f}%', ha='center', va='center')
plt.xlabel('average charge difference')
plt.title('Average charge difference [phospho - regular]')
plt.legend()
plt.savefig('D:/RawData/CSD/pub/plots/phospho/phospho_nonphospho_z_distance.svg')
plt.close()

bin_edges = np.linspace(0.0, 1.0, 101)
bottoms = np.zeros(100)
for n_phos, sims in cos_sims.groupby('Phospho_count'):
    bins, _ = np.histogram(sims['cos'], bins=bin_edges)
    plt.bar((bin_edges[:-1] + bin_edges[1:]) / 2, bins, bin_edges[1] - bin_edges[0], bottoms, label=f'{n_phos}-Phospho')
    bottoms += bins
cos_dist = cos_sims['cos'].values
plt.hist(cos_dist, bins=100)
plt.axvline(np.mean(cos_dist), ls='--', color='navy')
plt.xlim(left=-0.02, right=1.02)
plt.text(np.mean(cos_dist) * 0.9 , plt.ylim()[1] * 0.97, f'Mean: {np.mean(cos_dist):.4f}', ha='right', va='top')
plt.axvspan(plt.xlim()[0], 0.95, color='green', alpha=0.2)
plt.text((plt.xlim()[1] + plt.xlim()[0]) / 2 , plt.ylim()[1] * 0.5, f'Cosine similarity < 0.95: {np.mean(cos_dist < 0.95) * 100:.1f}%', ha='center', va='center')
plt.xlabel('cosine similarity')
plt.title('Cosine similarity [phospho - regular]')
plt.legend()
plt.savefig('D:/RawData/CSD/pub/plots/phospho/phospho_nonphospho_cos_similarity.svg')

phospho_size = phospho_paired.groupby('Phospho_count').size()
phospho_size /= phospho_size[1:].sum()
plt.bar(np.arange(1, phospho_size.shape[0]), phospho_size[1:])
plt.title('Number of phosphosites / peptide')
plt.savefig('D:/RawData/CSD/pub/plots/phospho/phospho_paired_number_of_phospho.svg')

###Properties of shifted peptides
z_shifts['n_basic'] = z_shifts['proforma'].str.count('[KHR]') - z_shifts['proforma'].str.count('\[Acetyl\]-') + 1
z_shifts['M'] = z_shifts['proforma'].apply(lambda s: ProForma.parse(s).mass)
z_shifts['sequence'] = z_shifts['proforma'].str.replace('\[.+?\]\-?', '', regex=True)
z_shifts['GRAVY'] = z_shifts['sequence'].apply(gravy)

#plotting different properties
pos_index = z_shifts['shift'] > 0.1
neg_index = z_shifts['shift'] < -0.1
no_index = np.abs(z_shifts['shift']) < 0.1

plt.figure(figsize=(8.5, 8))

#Number of basic sites
plt.subplot(3, 2, 1)
sites = Counter(z_shifts.loc[neg_index, 'n_basic'])
plt.bar(np.array(list(sites.keys())) + 0.3, np.array(list(sites.values())) / sum(sites.values()),
        width=0.3, label='decreased')
sites = Counter(z_shifts.loc[pos_index, 'n_basic'])
plt.bar(np.array(list(sites.keys())) - 0.3, np.array(list(sites.values())) / sum(sites.values()),
        width=0.3, label='increased')
sites = Counter(z_shifts.loc[no_index, 'n_basic'])
plt.bar(np.array(list(sites.keys())) - 0.0, np.array(list(sites.values())) / sum(sites.values()),
        width=0.3, label='unchanged')
plt.xlim(1.5, 7.5)
plt.legend()
plt.xlabel('Number of basic sites')
plt.ylabel('Relative abundance')

#GRAVY
plt.subplot(3, 2, 2)
kdeplot(z_shifts.loc[neg_index, 'GRAVY'], label='decreased')
kdeplot(z_shifts.loc[pos_index, 'GRAVY'], label='increased')
kdeplot(z_shifts.loc[no_index, 'GRAVY'], label='unchanged')
plt.legend()
plt.xlabel('GRAVY')

#charge
plt.subplot(3, 2, 3)
kdeplot(z_shifts.loc[neg_index, 'charge'], label='decreased')
kdeplot(z_shifts.loc[pos_index, 'charge'], label='increased')
kdeplot(z_shifts.loc[no_index, 'charge'], label='unchanged')
plt.legend()
plt.xlabel('Charge of dephosphorylated peptide')

#overcharge
plt.subplot(3, 2, 4)
kdeplot(z_shifts.loc[neg_index, 'charge'] - z_shifts.loc[neg_index, 'n_basic'], label='decreased')
kdeplot(z_shifts.loc[pos_index, 'charge'] - z_shifts.loc[pos_index, 'n_basic'], label='increased')
kdeplot(z_shifts.loc[no_index, 'charge'] - z_shifts.loc[no_index, 'n_basic'], label='unchanged')
plt.legend()
plt.xlabel('Overcharging')

#basic sites per 1000 Da
plt.subplot(3, 2, 5)
kdeplot(1000 * z_shifts.loc[neg_index, 'n_basic'] / z_shifts.loc[neg_index, 'M'], label='decreased')
kdeplot(1000 * z_shifts.loc[pos_index, 'n_basic'] / z_shifts.loc[pos_index, 'M'], label='increased')
kdeplot(1000 * z_shifts.loc[no_index, 'n_basic'] / z_shifts.loc[no_index, 'M'], label='unchanged')
plt.legend()
plt.xlabel('Basic sites per 1000 Da')

plt.subplot(3, 2, 6)
kdeplot(z_shifts.loc[neg_index, 'M'], label='decreased')
kdeplot(z_shifts.loc[pos_index, 'M'], label='increased')
kdeplot(z_shifts.loc[no_index, 'M'], label='unchanged')
plt.legend()
plt.xlabel('Mass (Da)')

plt.tight_layout()
plt.savefig('D:/RawData/CSD/pub/plots/phospho/phospho_pair_properties.svg')

#CSD spread
#adding CSDs to phosphorylated peptides
z_shifts_csd = z_shifts.merge(peptides.loc[:, :'6'], on='proforma', how='inner')
z_shifts_csd = z_shifts_csd[(z_shifts_csd.loc[:, '1':'6'] > 0).sum(axis=1) > 1].copy() #exclude single charge peptides

#adding CSD to dephosphorylated peptides
z_shifts_csd_dp = z_shifts.drop('proforma', axis=1).merge(peptides.loc[:, :'6'],
                                                          left_on='dephosphorylated', right_on='proforma', how='inner')
z_shifts_csd_dp = z_shifts_csd_dp[(z_shifts_csd_dp.loc[:, '1':'6'] > 0).sum(axis=1) > 1].copy() #exclude single charge peptides

#applying class labels
def class_label(x):
    if x < -0.1:
        return 'negative'
    elif x > 0.1:
        return 'positive'
    else:
        return 'neutral'

z_shifts_csd['class'] = z_shifts_csd['shift'].apply(class_label)
z_shifts_csd_dp['class'] = z_shifts_csd_dp['shift'].apply(class_label)

#since dephosphorylated sequences can be duplicated (i.e. coming from phosphorylation isoforms),
#we need to remove these, but keeping the class
z_shifts_csd_dp.drop_duplicates(['class', 'proforma'], inplace=True)

#cumulative CSD plot
for (cl, col) in zip(['negative', 'positive'], ['tab:blue', 'tab:orange']):
    plt.hist(z_shifts_csd.loc[z_shifts_csd['class'] == cl, '1':'6'].max(axis=1), bins=np.linspace(0.5, 1.0, 21),
             cumulative=True, histtype='step', density=True, color=col, label=f'{cl} (phosphorylated)')
    plt.hist(z_shifts_csd_dp.loc[z_shifts_csd_dp['class'] == cl, '1':'6'].max(axis=1), bins=np.linspace(0.5, 1.0, 21),
             cumulative=True, histtype='step', density=True, color=col, ls='--', label=f'{cl} (dephosphorylated)')

plt.xlabel('Relative abundance of the major charge state')
plt.ylabel('Proportion of peptides')
plt.xlim((0.5, 1))
plt.legend(loc='upper left')
plt.savefig('D:/RawData/CSD/pub/plots/phospho/phospho_major_charge.svg')

#weighted STD
for (cl, col) in zip(['negative', 'positive'], ['tab:blue', 'tab:orange']):
    kdeplot(weighted_std(np.broadcast_to(np.arange(1,7), z_shifts_csd.loc[z_shifts_csd['class'] == cl, '1':'6'].shape),
                         z_shifts_csd.loc[z_shifts_csd['class'] == cl, '1':'6'], 1), color=col, label=f'{cl} (phosphorylated)')
    kdeplot(weighted_std(np.broadcast_to(np.arange(1,7), z_shifts_csd_dp.loc[z_shifts_csd_dp['class'] == cl, '1':'6'].shape),
                         z_shifts_csd_dp.loc[z_shifts_csd_dp['class'] == cl, '1':'6'], 1), color=col, ls='--',
            label=f'{cl} (dephosphorylated)')

plt.xlabel('Weighted standard deviation of CSD')
plt.legend()
plt.xlim(-0.1, 1,1)
plt.savefig('D:/RawData/CSD/pub/plots/phospho/phospho_weighted_std.svg')

#find phosphopeptides that share dephosphorylated sequences, but have different position or number of phosphorylations
phospho_isomers = peptides.groupby('dephosphorylated').filter(lambda f: np.logical_and(np.all(f['Phospho_count'] > 0),
                                                                                       f.shape[0] > 1))

group_size = Counter(phospho_isomers.groupby('dephosphorylated').size())
plt.bar(group_size.keys(), group_size.values(), align='center')
plt.xticks(sorted(group_size.keys()), sorted(group_size.keys()))
plt.text(plt.xlim()[1] * 0.95, plt.ylim()[1] * 0.95,
         f'Total groups: {phospho_isomers.groupby("dephosphorylated").size().shape[0]}',
         ha='right', va='top')
plt.xlabel('Group size')
plt.ylabel('Nr cases')
plt.title('Size of isomer groups')
plt.savefig('D:/RawData/CSD/pub/plots/phospho/phospho_isomers_group_size.svg')

#largest (in magnitude) changes for phosphopeptide positional isomers (same total number of phosphosites)
z_dist = phospho_isomers.groupby(['dephosphorylated', 'Phospho_count']).filter(lambda g: g.shape[0] > 1).groupby(['dephosphorylated', 'Phospho_count']).apply(max_z_distance_i)
cos_dist = phospho_isomers.groupby(['dephosphorylated', 'Phospho_count']).filter(lambda g: g.shape[0] > 1).groupby(['dephosphorylated', 'Phospho_count']).apply(max_cos_distance_i)

bin_edges = np.linspace(z_dist.min(), z_dist.max(), 101)
bottoms = np.zeros(100)
for n in z_dist.index.levels[1]:
    bins, _ = np.histogram(z_dist[:, n], bins=bin_edges)
    plt.bar((bin_edges[:-1] + bin_edges[1:]) / 2, bins, bin_edges[1] - bin_edges[0], bottoms, label=f'{n}-Phospho')
    bottoms += bins
plt.axvline(np.mean(z_dist), ls='--', color='navy')
plt.xlim(left=z_dist.min() - 0.02, right=z_dist.max() + 0.02)
plt.text(np.mean(z_dist) * 1.1 , plt.ylim()[1] * 0.97, f'Mean: {np.mean(z_dist):.4f}', ha='left', va='top')
plt.axvspan(0.1, plt.xlim()[1], color='green', alpha=0.2)
plt.text((plt.xlim()[1] + 0.1) / 2 , plt.ylim()[1] * 0.5, f'Difference > 0.1: {np.mean(z_dist > 0.1) * 100:.1f}%', ha='center', va='center')
plt.xlabel('absolute average charge difference')
plt.title('Average charge difference [max of isoforms]')
plt.legend()
plt.savefig('D:/RawData/CSD/pub/plots/phospho/phospho_isomers_z_distance.svg')
plt.close()

bin_edges = np.linspace(0.0, 1.0, 101)
bottoms = np.zeros(100)
for n in cos_dist.index.levels[1]:
    bins, _ = np.histogram(cos_dist[:, n], bins=bin_edges)
    plt.bar((bin_edges[:-1] + bin_edges[1:]) / 2, bins, bin_edges[1] - bin_edges[0], bottoms, label=f'{n}-Phospho')
    bottoms += bins
plt.axvline(np.mean(cos_dist), ls='--', color='navy')
plt.xlim(left=-0.02, right=1.02)
plt.text(np.mean(cos_dist) * 0.9 , plt.ylim()[1] * 0.97, f'Mean: {np.mean(cos_dist):.4f}', ha='right', va='top')
plt.axvspan(plt.xlim()[0], 0.95, color='green', alpha=0.2)
plt.text((plt.xlim()[1] + plt.xlim()[0]) / 2 , plt.ylim()[1] * 0.5, f'Cosine similarity < 0.95: {np.mean(cos_dist < 0.95) * 100:.1f}%', ha='center', va='center')
plt.xlabel('cosine similarity')
plt.title('Cosine similarity [max of isoforms]')
plt.legend()
plt.savefig('D:/RawData/CSD/pub/plots/phospho/phospho_isomers_cos_similarity.svg')

#phosphopeptides with different number of phosphosites
site_isomers_z = pd.DataFrame(phospho_isomers.groupby('dephosphorylated').apply(max_z_sites).tolist())
site_isomers_cos = pd.DataFrame(phospho_isomers.groupby('dephosphorylated').apply(min_cos_sites).tolist())

bin_edges = np.linspace(np.nanmin(site_isomers_z.values), np.nanmax(site_isomers_z.values), 101)
for column in site_isomers_z.columns:
    zs = site_isomers_z.loc[np.isfinite(site_isomers_z[column]), column]
    plt.hist(zs, bins=bin_edges)
    plt.axvline(np.mean(zs), ls='--', color='navy')
    plt.xlim(left=np.nanmin(site_isomers_z.values) - 0.02, right= np.nanmax(site_isomers_z.values) + 0.02)
    plt.text(np.mean(zs) * 1.1 , plt.ylim()[1] * 0.97, f'Mean: {np.mean(zs):.4f}', ha='left', va='top')
    plt.axvspan(0.1, plt.xlim()[1], color='green', alpha=0.2)
    plt.text((plt.xlim()[1] + 0.1) / 2 , plt.ylim()[1] * 0.5, f'Difference > 0.1: {np.mean(zs > 0.1) * 100:.1f}%', ha='center', va='center')
    plt.xlabel('absolute average charge difference')
    plt.title(f'Average charge difference [max of {column}]')
    plt.savefig(f'D:/RawData/CSD/pub/plots/phospho/phospho_isomers_z_distance_{column}.svg')
    plt.close()


bin_edges = np.linspace(0.0, 1.0, 101)
for column in site_isomers_cos.columns:
    coses = site_isomers_cos.loc[np.isfinite(site_isomers_cos[column]), column]
    plt.hist(coses, bins=bin_edges)
    plt.axvline(np.mean(coses), ls='--', color='navy')
    plt.xlim(left=-0.02, right=1.02)
    plt.text(np.mean(coses) * 0.9 , plt.ylim()[1] * 0.97, f'Mean: {np.mean(coses):.4f}', ha='right', va='top')
    plt.axvspan(plt.xlim()[0], 0.95, color='green', alpha=0.2)
    plt.text((plt.xlim()[1] + plt.xlim()[0]) / 2 , plt.ylim()[1] * 0.5, f'Cosine similarity < 0.95: {np.mean(coses < 0.95) * 100:.1f}%', ha='center', va='center')
    plt.xlabel('cosine similarity')
    plt.title(f'Cosine similarity [max of {column}]')
    plt.savefig(f'D:/RawData/CSD/pub/plots/phospho/phospho_isomers_cos_similarity_{column}.svg')
    plt.close()

###Charge shift peptides
#filter to singly-phosphorylated peptides
z_shifts = z_shifts.loc[z_shifts['Phospho_count'] == 1, :].copy()

#positive shift
phospho_sequences = z_shifts.loc[z_shifts['shift'] > 0.1, 'proforma'].values
pos_motifs = reduce(lambda a,b: a + b, [extract_padded_sequence(seq, 7) for seq in phospho_sequences], [])
with open('D:/RawData/CSD/pub/plots/phospho/motif_pos_span7.txt', 'w') as out:
    out.write('\n'.join(pos_motifs))

#non-positive shift
phospho_sequences = z_shifts.loc[z_shifts['shift'] <= 0.1, 'proforma'].values
nonpos_motifs = reduce(lambda a,b: a + b, [extract_padded_sequence(seq, 7) for seq in phospho_sequences], [])
with open('D:/RawData/CSD/pub/plots/phospho/motif_nonpos_span7.txt', 'w') as out:
    out.write('\n'.join(nonpos_motifs))

#negative shift
phospho_sequences = z_shifts.loc[z_shifts['shift'] < -0.1, 'proforma'].values
neg_motifs = reduce(lambda a,b: a + b, [extract_padded_sequence(seq, 7) for seq in phospho_sequences], [])
with open('D:/RawData/CSD/pub/plots/phospho/motif_neg_span7.txt', 'w') as out:
    out.write('\n'.join(neg_motifs))

#non-negative shift
phospho_sequences = z_shifts.loc[z_shifts['shift'] >= -0.1, 'proforma'].values
nonneg_motifs = reduce(lambda a,b: a + b, [extract_padded_sequence(seq, 7) for seq in phospho_sequences], [])
with open('D:/RawData/CSD/pub/plots/phospho/motif_nonneg_span7.txt', 'w') as out:
    out.write('\n'.join(nonneg_motifs))

#overcharging/undercharging
#only phospho peptides
peptides = peptides[peptides['Phospho_count'] > 0].copy()
#count basic sites
peptides['n_basic'] = peptides['proforma'].str.count('[KHR]') - peptides['proforma'].str.count('\[Acetyl\]-') + 1

#averge charge vs mass
plt.figure(figsize=(7, 7))
for n_basic, pep_group in peptides.groupby('n_basic'):
    if n_basic <= 6:
        plt.subplot(3, 2, n_basic)
        under = pep_group['av_charge'] - n_basic < -0.05
        over = pep_group['av_charge'] - n_basic > 0.05
        plt.scatter(pep_group.loc[under, 'M'], pep_group.loc[under, 'av_charge'], alpha=0.5, s=2)
        plt.scatter(pep_group.loc[over, 'M'], pep_group.loc[over, 'av_charge'], alpha=0.5, s=2)
        plt.scatter(pep_group.loc[~over & ~under, 'M'], pep_group.loc[~over & ~under, 'av_charge'], alpha=0.5, s=2)
        plt.axhline(n_basic, ls='--', color='darkgreen', lw=1)
        lowessFit = lowess(pep_group['av_charge'].values, pep_group['M'].values, is_sorted=False, frac=0.05, it=1)
        plt.plot(lowessFit[:, 0], lowessFit[:, 1], lw=1, color='tab:gray', label='Lowess fit')
        plt.ylim((0.5, 6.5))
        plt.yticks([1, 2, 3, 4, 5, 6], [1, 2, 3, 4, 5, 6])
        plt.xlabel('Peptide mass')
        plt.ylabel('Observed charge')
        plt.title(f'{n_basic} basic sites')

plt.tight_layout()
plt.savefig('D:/RawData/CSD/pub/plots/phospho/phospho_under_overcharging.png', dpi=150)
```
