## Supplementary material for "Exploiting Charge State Distribution to Probe Intramolecular Interactions in Gas-Phase Phosphopeptides and Enhance Proteomics Analyses": Scripts for data analysis and generation of results and intermediate plots: shotgun.py

```
# -*- coding: utf-8 -*-
"""
Using CSD in shotgun context
Need:
    Percolator .pin files with CSD features included
    corresponding biosaur2 features file for every raw file
    ThermoRawFileParser (has to be in PATH and as `thermoparser`) and corresponding raw files
    Trained CSD model

    All files has to share the base name (before the file extension) and location

Output:
    Creates .pin file with feature subsets and runs Percoltor on each of them
    Creates .pin file with shuffled features and runs Percolator
    Produces plots and graphs

Usage:
    plain script (not to be executed from command line)

@author: Vladimir
"""

import pandas as pd
import numpy as np
import matplotlib.pyplot as plt
from os import mkdir, path, listdir
from subprocess import check_output, CalledProcessError
from lxml import etree

def read_pin(filepath):
    with open(filepath, 'r') as input_file:
        headers = input_file.readline().strip().split('\t')
        num_headers = len(headers)
        data = []
        for line in input_file.readlines():
            fields = line.strip().split('\t', num_headers - 1)
            data.append({headers[n]: value for (n, value) in enumerate(fields)})

    return pd.DataFrame(data)

def write_pin(dataframe, subheader, filepath):
    with open(filepath, 'w') as output_file:
        output_file.write('\t'.join(dataframe.columns) + '\n')
        output_file.write(subheader + '\n')
        for row in dataframe.values:
            output_file.write('\t'.join([str(field) for field in row]) + '\n')


### DirectMS
#load file description
description = pd.read_csv('D:/RawData/CSD/DirectMS/DirectMS_files.tsv', sep='\t')
description['label'] = description.apply(lambda row: f'{row["Instrument"]}_{row["FAIMS"]}_{row["Sample load"]}_{row["Replicate"]}', axis=1)
description.set_index('filename', inplace=True)

processing_mapping = {'csd_cos': 'Complete CSD',
                      'nocsd_mark': 'Reduced CSD',
                      'nocsd': 'Regular'}

#loading DirectMS1 results
data = []
for processing in ['csd_cos', 'nocsd', 'nocsd_mark']:
    for filename in listdir(path.join('D:/RawData/CSD/DirectMS/deeplc/minlh1', processing)):
        if filename.endswith('_proteins.tsv'):
            proteins = pd.read_csv(path.join('D:/RawData/CSD/DirectMS/deeplc/minlh1', processing, filename), sep='\t')
            proteins = proteins.loc[~proteins['decoy'], :].copy()
            data.append({'processing': processing,
                         'filename': filename.replace('.features_proteins.tsv', ''),
                         'n_proteins': proteins.shape[0],
                         'proteins': proteins})

result = pd.DataFrame(data)

#renaming of columns
result['label'] = result['filename'].map(description['label'].to_dict())
result['Processing'] = result['processing'].map(processing_mapping)

#plotting protein counts
pd.pivot_table(result, values='n_proteins',
               index=['label'], columns=['Processing']).plot.bar(rot=90, figsize=(12, 7))
plt.axvspan(-0.5, 2.5, color='tab:olive', alpha=0.2)
plt.ylabel('Proteins')
plt.xlabel(None)
plt.tight_layout()

plt.savefig('D:/RawData/CSD/pub/plots/shotgun/DirectMS.svg')


### MSMS + Percolator
## feature exclusion
charge_features = ['1', '2', '3', '4', '5', '6', 'z']
fragment_features = ['MeanErrorTop7', 'sqMeanErrorTop7', 'StdevErrorTop7', 'lnExplainedIonCurrentRatio',
                     'lnNTermIonCurrentRatio', 'lnCTermIonCurrentRatio', 'lnMS2IonCurrent']
score = ['RawScore', 'DeNovoScore', 'ScoreRatio', 'Energy', 'lnEValue']
default1 = ['SpecId', 'Label', 'ScanNr', 'ExpMass', 'CalcMass', 'nMatch', 'cover']
default2 = ['Peptide', 'Proteins']

cond = {'charge': default1 + charge_features,
        'score+charge': default1 + score + charge_features,
        'fragment+charge': default1 + fragment_features + charge_features,
        'fragment+score+charge': default1 + fragment_features + score + charge_features,
        'score': default1 + score,
        'fragment': default1 + fragment_features,
        'fragment+score': default1 + fragment_features + score}

#preparing percolator input with limited features and running percolator
parent = 'D:/RawData/Eclipse/csd_6'
for filename in listdir(parent):
    if filename.endswith('_csd.pin'):
        print(f'Working on {filename}')
        filepath = path.join(parent, filename)

        pin_data = read_pin(filepath)
        subheader = pin_data.loc[0, :].astype(str)[:-2]
        pin_data.drop(0, inplace=True)

        subheader['nMatch'] = '1'
        subheader['cover'] = '1'
        subheader = subheader.str.replace('nan', '0')

        for subpath, preserve in cond.items():
            print(f'Preserve {subpath}')
            if not path.exists(path.join(parent, subpath)):
                mkdir(path.join(parent, subpath))

            subheader_string = '\t'.join(subheader[preserve].copy())
            new_filepath = path.join(parent, subpath, filename)

            write_pin(pin_data[preserve + default2].copy(), subheader_string, new_filepath)

            #running percolator
            perc_params = ['C:/Program Files/percolator-v3-05/bin/percolator.exe', '-Y', '-P', 'DECOY_', '-c', '-g',
                            '-f', 'D:/Fasta/uniprot-human_decoy.fasta', '-z', 'trypsin',
                            '-X', new_filepath.replace('_csd.pin', '.pout.xml'), '-w',
                            new_filepath.replace('_csd.pin', '.weights'),
                            new_filepath]

            try:
                print('Running Percolator...')
                check_output(perc_params, shell=True)
            except CalledProcessError as ex:
                print(f'Percolator execution failed\nOutput:\n{ex.output}')

            weights = pd.read_csv(new_filepath.replace('_csd.pin', '.weights'), sep='\t')
            weights = weights.iloc[np.array([1, 0, 0, 1, 0, 0, 1, 0]).astype(bool), :].astype(float).copy()

            #plotting weights
            plt.bar(np.arange(weights.shape[1]), weights.mean(axis=0))
            plt.xticks(np.arange(weights.shape[1]), weights.columns, rotation=90)
            plt.tight_layout()
            plt.savefig(new_filepath.replace('_csd.pin', '_weights.png'))
            plt.close()

#usual feature set
subpath = 'usual'
for filename in listdir(parent):
    if filename.endswith('_csd.pin'):
        print(f'Working on {filename}')
        filepath = path.join(parent, filename)

        pin_data = read_pin(filepath)
        subheader = pin_data.loc[0, :].astype(str)[:-2]
        pin_data.drop(0, inplace=True)

        if not path.exists(path.join(parent, subpath)):
            mkdir(path.join(parent, subpath))

        subheader_string = '\t'.join(subheader.drop(['nMatch', 'cover'] + charge_features))
        new_filepath = path.join(parent, subpath, filename)

        write_pin(pin_data.drop(['nMatch', 'cover'] + charge_features, axis=1), subheader_string, new_filepath)

        #running percolator
        perc_params = ['C:/Program Files/percolator-v3-05/bin/percolator.exe', '-Y', '-P', 'DECOY_', '-c', '-g',
                        '-f', 'D:/Fasta/uniprot-human_decoy.fasta', '-z', 'trypsin',
                        '-X', new_filepath.replace('_csd.pin', '.pout.xml'), '-w',
                        new_filepath.replace('_csd.pin', '.weights'),
                        new_filepath]

        try:
            print('Running Percolator...')
            check_output(perc_params, shell=True)
        except CalledProcessError as ex:
            print(f'Percolator execution failed\nOutput:\n{ex.output}')

        weights = pd.read_csv(new_filepath.replace('_csd.pin', '.weights'), sep='\t')
        weights = weights.iloc[np.array([1, 0, 0, 1, 0, 0, 1, 0]).astype(bool), :].astype(float).copy()

        #plotting weights
        plt.bar(np.arange(weights.shape[1]), weights.mean(axis=0))
        plt.xticks(np.arange(weights.shape[1]), weights.columns, rotation=90)
        plt.tight_layout()
        plt.savefig(new_filepath.replace('_csd.pin', '_weights.png'))
        plt.close()

#parsing identification results
#file info
fileinfo = pd.read_csv('D:/RawData/CSD/selected_filenames.csv')
fileinfo.sort_values(['Gradient length (min)', 'Laboratory', 'Instrument', 'MS1 resolution', 'MS2 resolution', 'MS1 AGC', 'Sample load', 'Replicate'],
                      inplace=True)
fileinfo['MS1 AGC'] = fileinfo['MS1 AGC'].apply(lambda x: '{}e6'.format(int(x/1e6)))

fileinfo['label'] = fileinfo.loc[:, 'Laboratory':].apply(lambda r: '_'.join(r.astype(str)), axis=1)
fileinfo.set_index('filename', inplace=True)

parent = 'D:/RawData/Eclipse/csd_6'
id_data = []
for subpath in list(cond.keys()) + ['usual']:
    for filename in listdir(path.join(parent, subpath)):
        if filename.endswith('.pout.xml'):
            poutFile = etree.parse(path.join(parent, subpath, filename))
            ns = {"a": poutFile.getroot().nsmap[None]}

            id_data.append({'Name': filename,
                            'Experiment': subpath,
                            'PSMs': int(poutFile.xpath('//a:psms_qlevel', namespaces=ns)[0].text),
                            'Peptides': int(poutFile.xpath('//a:peptides_qlevel', namespaces=ns)[0].text),
                            'Proteins': int(poutFile.xpath('//a:proteins_qlevel', namespaces=ns)[0].text)})

id_data = pd.DataFrame(id_data).sort_values('Experiment')
id_data['Label'] = id_data['Name'].str.slice(0, -9).map(fileinfo['label'].to_dict())

plt.figure(figsize=(16, 8))
items = ['PSMs', 'Peptides', 'Proteins']
for i, t in enumerate(items):
    ax = plt.subplot(1, 3, i + 1)
    pd.pivot(id_data, index='Label', columns='Experiment')[t].plot(kind='bar', ax=ax)
    plt.title(t)
    plt.xlabel(None)

plt.tight_layout()
plt.savefig('D:/RawData/CSD/pub/plots/shotgun/MSMS_Percolator_feature.svg')

#phosphodata
fileinfo = pd.read_csv('D:/RawData/PhosphoCSD/shotgun/selected_filenames.csv')
fileinfo.sort_values(['Gradient length (min)', 'Laboratory', 'Instrument', 'MS1 resolution', 'MS2 resolution', 'Replicate'],
                      inplace=True)

fileinfo['label'] = fileinfo.loc[:, 'Laboratory':].apply(lambda r: '_'.join(r.astype(str)), axis=1)
fileinfo.set_index('filename', inplace=True)

parent = 'D:/RawData/PhosphoCSD/shotgun'
id_data = []
for filename in listdir(parent):
    if filename.endswith('.pout.xml'):
        experiment = 'CSD' if filename.find('_csd.') != -1 else 'Regular'
        poutFile = etree.parse(path.join(parent, filename))
        ns = {"a": poutFile.getroot().nsmap[None]}

        id_data.append({'Name': filename[:-9].replace('_csd', ''),
                        'Experiment': experiment,
                        'PSMs': int(poutFile.xpath('//a:psms_qlevel', namespaces=ns)[0].text),
                        'Peptides': int(poutFile.xpath('//a:peptides_qlevel', namespaces=ns)[0].text),
                        'Proteins': int(poutFile.xpath('//a:proteins_qlevel', namespaces=ns)[0].text)})

        if experiment == 'CSD':
            weights = pd.read_csv(path.join(parent, filename.replace('_csd.pout.xml', '_csd.weights')), sep='\t')
            weights = weights.iloc[np.array([1, 0, 0, 1, 0, 0, 1, 0]).astype(bool), :].astype(float).copy()

            #plotting weights
            plt.bar(np.arange(weights.shape[1]), weights.mean(axis=0))
            plt.xticks(np.arange(weights.shape[1]), weights.columns, rotation=90)
            plt.tight_layout()
            plt.savefig(path.join(parent, filename.replace('_csd.pout.xml', '_weights.png')))
            plt.close()

id_data = pd.DataFrame(id_data).sort_values('Experiment')
id_data['Label'] = id_data['Name'].map(fileinfo['label'].to_dict())

plt.figure(figsize=(16, 8))
items = ['PSMs', 'Peptides', 'Proteins']
for i, t in enumerate(items):
    ax = plt.subplot(1, 3, i + 1)
    pd.pivot(id_data, index='Label', columns='Experiment')[t].plot(kind='bar', ax=ax)
    plt.title(t)
    plt.xlabel(None)

plt.tight_layout()
plt.savefig('D:/RawData/CSD/pub/plots/shotgun/MSMS_phospho_Percolator.svg')


#shuffling scores
ratio = 0.05
for filename in listdir(parent):
    if filename.endswith('_csd.pin'):
        print(f'Working on {filename}')
        filepath = path.join(parent, filename)

        pin_data = read_pin(filepath)
        subheader = pin_data.loc[0, :].astype(str)[:-2]
        pin_data.drop(0, inplace=True)

        subheader['nMatch'] = '1'
        subheader['cover'] = '1'
        subheader = subheader.str.replace('nan', '0')

        #shuffling ratio of data
        subset = np.random.choice(pin_data.index, int(pin_data.shape[0] * ratio), replace=False)
        subset_shuffled = subset.copy()
        np.random.shuffle(subset_shuffled)

        shuffle = {k: v for (k,v) in zip(subset, subset_shuffled)}
        new_index = pin_data.rename(index=shuffle).index

        subpath = 'shuffle'
        preserve = cond['score']
        print('Preserve score')
        if not path.exists(path.join(parent, subpath)):
            mkdir(path.join(parent, subpath))

        subheader_string = '\t'.join(subheader[preserve].copy())
        new_filepath = path.join(parent, subpath, filename)

        limited_data = pin_data[preserve + default2].copy()

        limited_data[['nMatch', 'cover'] + score] = limited_data.reindex(new_index)[['nMatch', 'cover'] + score].values

        write_pin(limited_data, subheader_string, new_filepath)

        #running percolator
        perc_params = ['C:/Program Files/percolator-v3-05/bin/percolator.exe', '-Y', '-P', 'DECOY_', '-c', '-g',
                        '-f', 'D:/Fasta/uniprot-human_decoy.fasta', '-z', 'trypsin',
                        '-X', new_filepath.replace('_csd.pin', '.pout.xml'), '-w',
                        new_filepath.replace('_csd.pin', '.weights'),
                        new_filepath]

        try:
            print('Running Percolator...')
            check_output(perc_params, shell=True)
        except CalledProcessError as ex:
            print(f'Percolator execution failed\nOutput:\n{ex.output}')

        weights = pd.read_csv(new_filepath.replace('_csd.pin', '.weights'), sep='\t')
        weights = weights.iloc[np.array([1, 0, 0, 1, 0, 0, 1, 0]).astype(bool), :].astype(float).copy()

        plt.bar(np.arange(weights.shape[1]), weights.mean(axis=0))
        plt.xticks(np.arange(weights.shape[1]), weights.columns, rotation=90)
        plt.tight_layout()
        plt.savefig(new_filepath.replace('_csd.pin', '_weights.png'))
        plt.close()

        subpath = 'shuffle_charge'
        preserve = cond['score+charge']
        print('Preserve score+charge')
        if not path.exists(path.join(parent, subpath)):
            mkdir(path.join(parent, subpath))

        subheader_string = '\t'.join(subheader[preserve].copy())
        new_filepath = path.join(parent, subpath, filename)

        limited_data = pin_data[preserve + default2].copy()

        limited_data[['nMatch', 'cover'] + score] = limited_data.reindex(new_index)[['nMatch', 'cover'] + score].values

        write_pin(limited_data, subheader_string, new_filepath)

        #running percolator
        perc_params = ['C:/Program Files/percolator-v3-05/bin/percolator.exe', '-Y', '-P', 'DECOY_', '-c', '-g',
                        '-f', 'D:/Fasta/uniprot-human_decoy.fasta', '-z', 'trypsin',
                        '-X', new_filepath.replace('_csd.pin', '.pout.xml'), '-w',
                        new_filepath.replace('_csd.pin', '.weights'), new_filepath,
                        '2>', new_filepath.replace('_csd.pin', '.stderr')]

        try:
            print('Running Percolator...')
            check_output(perc_params, shell=True)
        except CalledProcessError as ex:
            print(f'Percolator execution failed\nOutput:\n{ex.output}')

        weights = pd.read_csv(new_filepath.replace('_csd.pin', '.weights'), sep='\t')
        weights = weights.iloc[np.array([1, 0, 0, 1, 0, 0, 1, 0]).astype(bool), :].astype(float).copy()

        plt.bar(np.arange(weights.shape[1]), weights.mean(axis=0))
        plt.xticks(np.arange(weights.shape[1]), weights.columns, rotation=90)
        plt.tight_layout()
        plt.savefig(new_filepath.replace('_csd.pin', '_weights.png'))
        plt.close()

id_data = []
for subpath in ['shuffle', 'shuffle_charge']:
    for filename in listdir(path.join(parent, subpath)):
        if filename.endswith('.pout.xml'):
            poutFile = etree.parse(path.join(parent, subpath, filename))
            ns = {"a": poutFile.getroot().nsmap[None]}

            id_data.append({'Name': filename,
                            'Experiment': subpath,
                            'PSMs': int(poutFile.xpath('//a:psms_qlevel', namespaces=ns)[0].text),
                            'Peptides': int(poutFile.xpath('//a:peptides_qlevel', namespaces=ns)[0].text),
                            'Proteins': int(poutFile.xpath('//a:proteins_qlevel', namespaces=ns)[0].text)})

id_data = pd.DataFrame(id_data).sort_values('Experiment')
id_data['Label'] = id_data['Name'].str.slice(0, -9).map(fileinfo['label'].to_dict())
id_data['Experiment'] = id_data['Experiment'].map({'shuffle': '5% shuffled', 'shuffle_charge': '5% shuffled + CSD'})

plt.figure(figsize=(16, 8))
items = ['PSMs', 'Peptides', 'Proteins']
for i, t in enumerate(items):
    ax = plt.subplot(1, 3, i + 1)
    pd.pivot(id_data, index='Label', columns='Experiment')[t].plot(kind='bar', ax=ax)
    plt.title(t)
    plt.xlabel(None)

plt.tight_layout()
plt.savefig('D:/RawData/CSD/pub/plots/shotgun/MSMS_Percolator_shuffle.svg')
```
