## Extended experimental details and additional results for "Exploiting Charge State Distribution to Probe Intramolecular Interactions in Gas-Phase Phosphopeptides and Enhance Proteomics Analyses"

### Extended Methods

#### Software tools

Unless stated otherwise, data manipulation and plotting were performed in Python (3.10.2) with the following modules installed: ***numpy*** (1.23.5)[1], ***scipy*** (1.10.0)[2], ***matplotlib*** (3.5.1)[3], ***pandas*** (1.4.0)[4], ***scikit-learn*** (1.0.2)[5], ***seaborn*** (0.11.2)[6], ***pyteomics*** (4.6)[7], ***keras*** (2.10.0)[8], ***statsmodels*** (0.13.1)[9], ***pygam*** (0.8.0)[10]. Relevant scripts are provided as supplementary material.

#### Database search (non-phosphorylated)

Raw files were searched using MSGF+ (2022.01.07)[11] against the UniProtKB human protein database supplemented with reversed decoy sequences. The following parameters were used: trypsin as a digestion enzyme; parent ion mass accuracy – 10 ppm, isotopic error of 0 and 1, precursor charge from 2 to 6 (both included), theoretical peptide length from 6 to 40 (both included), carbamidomethylation of cysteine as the fixed modification, oxidation of methionine and acetylation of protein N-terminus as variable ones. Instrument protocol and enzyme were set according to the LC-MS experiment. Search results were validated using Percolator (3.05)[12], q-values and PEPs calculated by Percolator were added to the resulting mzIdentML file, and PSMs were filtered to q-value < 0.01 for any further processing.

#### Database search (phosphorylated)

Deposited MaxQuant results were used for phosphoproteomics datasets when available. Raw files belonging to the PXD000138 dataset (synthetic phosphopeptides) were reanalyzed using MaxQuant (2.0.3.0)[13]. The search was performed against the deposited FASTA file. Trypsin has been selected as an enzyme and up to two missed cleavages were allowed. Carbamidomethylation of cysteine was used as fixed, and oxidation of methionine and phosphorylation of serine, threonine, and tyrosine were used as variable modifications. The minimal peptide length was 7, and the maximal peptide mass was 4600. Results were filtered to 0.01 FDR for proteins, peptides, and sites. All other parameters were set to their default values.

#### Phosphoproteomics datasets analysis

MaxQuant results (*msms.txt* file) were read using ***psm-utils*** (0.2.3)[14]. PSMs with all phosphorylated sites having a localization score above 0.95 and Andromeda score above 50 were considered for further processing. For the synthetic phosphopeptide datasets, peptides matching the synthesis goal were considered valid and accepted for further processing. Since *msms.txt* files contain only calibrated mass values, corresponding *evidence.txt* files were used to calculate the observed (non-calibrated) mass values matching the features detected in the raw files.

#### Charge-state distribution calculation

Raw files were converted to mzML format using ***ThermoRawFileParser*** (1.4.2)[15] with default parameters and further processed by ***biosaur2*** (0.2.11)[16] to detect LC-MS features. The minimal length of a hill was set to 2, and the mass accuracy for hills and isotopes was set to 8. Peptide-spectrum matches (PSMs) were mapped to detected LC-MS features, by the following criteria: PSM retention time has to be between the start and the end of LC-MS feature elution, the charge of the PSM has to match the charge of the LC-MS feature, and the absolute difference between peptide mass and the feature mass must be below 5 ppm. Unmatched features coeluting with the matched ones (overlapping retention time) and having similar mass (absolute difference below 5 ppm) were grouped. The averaged feature mass, the average elution apex, the smallest start, and the largest end time were extracted for every feature group and peptide sequence. If more than one peptide sequence was matched to a feature group, each of the peptide sequences had the same result associated with it. The masses were used to calculate theoretical monoisotopic m/z values for all charge states from 1 to 6 (including both ends), m/z values falling outside the m/z range of the LC-MS file were discarded. ThermoRawFileParser was used to produce extracted ion chromatograms (XICs) for every m/z value, using a 5-ppm mass window, and retention time borders determined earlier for every peptide. The area under the curve (AUC) was calculated for every XIC and, finally, XICs belonging to the same peptide were assembled into a vector representing abundances of the corresponding charge states (raw CSDs). Raw CSDs were calculated for every LC-MS run in the dataset and the results merged for the complete dataset. Next, all raw CSDs were normalized (divided by the sum of all XIC areas), and complete duplicates (the same peptide sequence, and normalized charge state abundancies) were discarded. CSDs were grouped by peptide sequence and the consensus CSD was calculated: cosine similarity was calculated for all members of the group and only the largest subgroup having cosine similarity above 0.9 for all its members was used for the consensus CSD calculation by averaging all individual CSDs; groups with no correlated samples were discarded; groups with single member were used without any processing. Consensus CSDs for every unique peptide sequence were stored for every dataset.

CSDs for the timsTOF data were calculated by the *extract_csd* script published in https://github.com/regev-lab/extract-csd (accessed 18.11.2023)[17] and used without any further processing.

#### CSD prediction model

Peptide sequences in a dataset were converted into single-letter codes, with special codes used for N-terminal acetylation, oxidated methionine, and phosphorylated serine, threonine, and tyrosine (25 in total). Cysteine was always considered carbamidomethylated and, thus, was encoded as cysteine. Peptides longer than 40 elements were discarded. For the deep learning model, peptide sequences were one-hot encoded, i.e. the input matrix for every peptide had 40 rows and 25 columns, one element in each row was set to one indicating the corresponding amino acid in the sequence, and all others set to zero. For all other models, the input was a row vector with 25 elements representing the number of amino acids of the corresponding type. The complete dataset was split into training (90%) and test (10%) parts (*train_test_split* from ***scikit-learn***). The following model classes were used for the prediction: *LinearRegression*, *KNeighborsRegressor* (with K = 7), *DecisionTreeRegressor*, from ***scikit-learn***, *GAM* (with lambda = 0.01) from ***pygam***, in the latter case 6 regressors were joined using *MultiOutputRegressor* from ***scikit-learn***. All other parameters were set to their default values. Deep learning model architecture was described earlier[18], briefly, the network consisted of a sequentially connected masking layer, two bidirectional long short-term memory (LSTM) layers, and two dense layers, the latest one had an output dimension of 6. The model was compiled with an Adam optimizer and negative cosine similarity loss function. The training was performed on a computer equipped with NVIDIA A100 GPU running ***tensorflow***/***keras*** 2.12.0 under Python 3.8.10. The training was performed with a batch size of 512, for a maximum of 100 epochs and early stopping monitoring loss on a validation set (validation split 0.2) and patience of 5. Three models using 10%, 30%, and 100% of the data selected at random were trained. Since a monotonous decrease in the prediction error (calculated on the holdout set) has been observed, the final model (trained on the complete dataset) was used.

#### DirectMS1 analysis

Raw files were converted to mzML using ***ThermoRawFileParser*** (1.4.2) and, next, ***biosaur2*** (0.2.11) has been used for feature detection. The minimal length of a hill was set to 1, and the mass accuracy for hills and isotopes was set to 8. The search was performed by ***ms1searchpy***[19] against the SwissProt human protein database, concatenated with decoy sequences generated by pseudo-shuffling[20]. The following parameters were used: minimum number of scans was 1, minimum number of isotopes was 2, no missed cleavages, minimum precursor charge was 1, initial precursor tolerance was 8 ppm, protein FDR was 1%, two staged RT training, employing additive model first and DeepLC model next, minimal candidate peptide length was 6. CSD-specific features were included in the feature table used in the machine-learning-based peptide-feature match (PFM) classification. For the reduced CSD experiment, the following features were included: the number of times a peptide sequence is observed in the PFM list; the number of unique charge states a peptide sequence is observed in; and the number of unique ion mobility values a peptide sequence is observed. For the complete CSD experiment, a CSD prediction step has been implemented in ***ms1searchpy*** (both reduced and complete options are included in version 2.6.3). To this end all PFMs were collected and m/z values corresponding to charge states from 1 to 4 (both included) were calculated; m/z values outside the m/z range of the instrument were set to zero; **ThermoRawFileParser** (1.4.2) was used to produce XICs using the calculated m/z values, 5 ppm tolerance window, and retention time borders of each LC-MS feature; AUC has been calculated for each XIC and these values were assembled to experimental CSD vectors; the model trained earlier was employed to predict CSDs for every unique peptide sequence; charge states corresponding to m/z values outside instrument m/z range were set to zero; both experimental and predicted CSD vectors were normalized; average observed charge state was calculated for both experimental and predicted CSDs as discussed earlier. The following properties were calculated for every PFM and added to the table for the machine-learning-assisted classification: spectral angle between the experimental and predicted CSD, the difference between experiment and prediction of each charge state, and average observed charge all expressed as z-score, i.e. number of standard deviations from the mean error of each property prediction. Note, that in the current implementation, the LC-MS file in vendor RAW format and **ThermoRawFileParser** (version 1.4.2 or later) are necessary to perform the XIC extraction.

#### MS/MS analysis with CSD

Raw files were searched using MSGF+ and Percolator and LC-MS features were calculated as described earlier. The Percolator input file was modified to include CSD-related features for each entry in it. Each entry in a Percolator input represents PSM (both target and decoy and without any quality filtering) and, thus, experimental CSD calculation followed the same basic steps as described earlier. Briefly, PSMs were mapped to LC-MS features, matched features were grouped by co-elution and mass, m/z values for charges from 1 to 6 were calculated, and, lastly, corresponding XICs were extracted using **ThermoRawFileParser** and AUCs assembled to experimental CSDs. The deep learning model was employed to predict CSDs for each unique peptide sequence, and later entries belonging to m/z values outside the instrument m/z range were zeroed and CSDs normalized. Errors between the experimental and predicted values of each charge state relative abundance and observed average charge represented as z-score were used as features for Percolator (feature names: *1*, *2*, *3*, *4*, *5*, *6*, *z*). For PSMs lacking experimental CSD, an average value of the corresponding features was used (for target and decoy populations separately). Finally, the simple matching score was calculated as two extra features: *nMatch* – the number of theoretical fragments matched in the fragmentation spectrum, and *cover* – *nMatch* divided by peptide length. A separate input file was generated for every combination of employed features and submitted to Percolator. Another set of input files was created by shuffling randomly score-related features with and without charge features.

### Supplementary Text 1

#### Building CSD prediction model

The K562 dataset (non-phosphorylated peptides) and phosphopeptides dataset were combined and used to train a deep-learning prediction model. The architecture of the model has been published earlier[18], but the output dimension layer has been changed to six to accommodate the size of the calculated CSD vectors. A holdout test set consisting of 27740 peptide sequences has been randomly selected from the dataset, the remaining part (249655 peptides) has been used during training and validation. The prediction accuracy (cosine similarity) calculated for the holdout set was 0.9889 and 0.9850 for all peptides and only phosphopeptides, respectively. Altogether, the model demonstrated a level of accuracy in line with the reproducibility of the previously observed CSD data. A slightly worse error for phosphopeptides could be due to the worse quality of the CSD data itself (such as erroneous phosphosite localization, see the discussion in the *Charge-state distribution of phosphorylated peptides* section) or just since phosphopeptides were a minor peptide class, representing about 15% of peptides in the dataset. Since we have observed that the CSD of sequence isomers were generally well correlated our initial predictor used simpler and more explainable models, using peptide amino acid composition as the input. Simpler models, however, demonstrated considerably lower prediction accuracy (summarized in Table A and Figure A). Thus, we can conclude that amino-acid interactions (that are not accounted for in the composition-based input) play an important role in the observed CSD. For our further investigation, we have used the deep learning model.

Table A. Prediction accuracy (mean cosine similarity) for different machine learning models

| Model | Training set error | Test set error |
|---|---|---|
| Linear regression | 0.8798 | 0.8790 |
| K-nearest neighbors (K=7) | 0.9126 | 0.8813 |
| Decision tree | 0.9997 | 0.9321 |
| Generalized additive model (per charge state) | 0.8933 | 0.8935 |
| Deep learning (LSTM) | 0.9902 | 0.9889 |

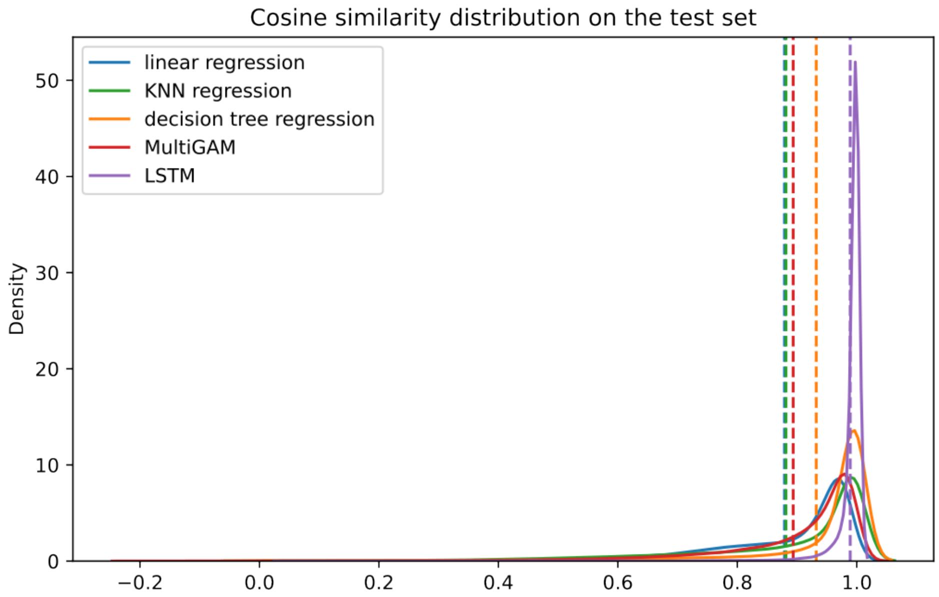


Figure A: Distribution of cosine similarity between predicted and experimental CSDs on the holdout set for different predictors. The dashed line corresponds to the mean value.

#### Investigation of human proteome digest with the trained model

The established CSD prediction model was used for a more systematic investigation of the discrimination performance of the CSD features. To that end, we performed *in silico* digestion of the complete human proteome (6 – 35 amino acids, up to 2 missed cleavages, no PTMs) resulting in 2243657 unique peptide sequences. The *in-silico* digest dataset is an order of magnitude larger, than the experimental (K562) one and, although a significant part of the peptides is unlikely to be observed in the LC-MS experiment (for example, due to low proteotypicity), it allows us to test the discriminatory performance in a much more complex scenario. We employed our trained model to predict the relative intensities for the first six charge states and DeepLC (1.1.2) to predict the relative retention time of each peptide, masses were calculated as uncharged monoisotopic masses. Figure B below shows the distribution of the cosine similarity between sequence and constitutional (including sequence) isomers, for comparison, we also show the distribution of the cosine similarity for 25000 randomly chosen peptide pairs. The overall performance appears to be quite similar to the one observed for the K562 dataset, i.e. discrimination between the sequence isomers is only slightly better than the reproducibility of CSD measurements, while constitutional isomers, in general, are distinguishable.

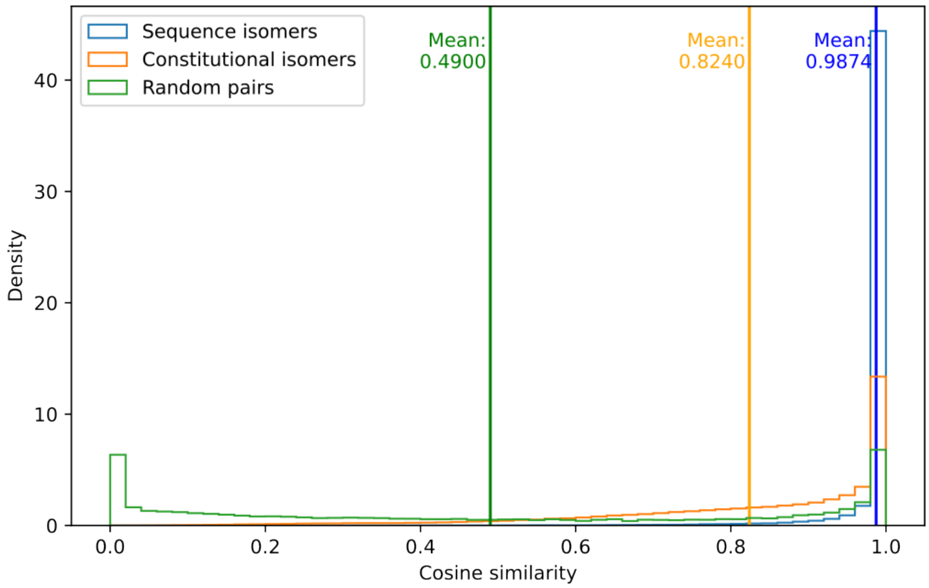


Figure B. Distribution of cosine similarity between peptide classes.

Next, we assessed the discrimination power by using the binning approach similar to the K562 dataset. Since DeepLC generates peptide retention times in relative units (from 0 to 10), we have used peak capacity (the number of fully resolved peaks over a complete elution window) instead of retention time for bin width calculation. Bin size was determined as 10 divided by peak capacity, for comparison, the binning used for the K562 dataset, corresponded to a peak capacity of 126 (63 minutes / 0.5 minutes). Results showing the influence of peak capacity are presented in Figure C. Despite visibly lower discriminatory power, for example, almost no peptides could be distinguished by mass alone (using 5 ppm bins), the changes in discrimination power are very similar to the one observed in the experimental (K562) dataset, confirming the validity of our earlier findings. The inclusion of CSDs allows a significant (approximately 20%) increase in the number of bins with the single peptide, despite an even larger increase observed when using retention time (especially with high peak capacity), applying CSD in combination with retention time proves to be even more beneficial. Cosine distance shows weaker discrimination compared to the average charge state. All other plots are provided as supplementary material.

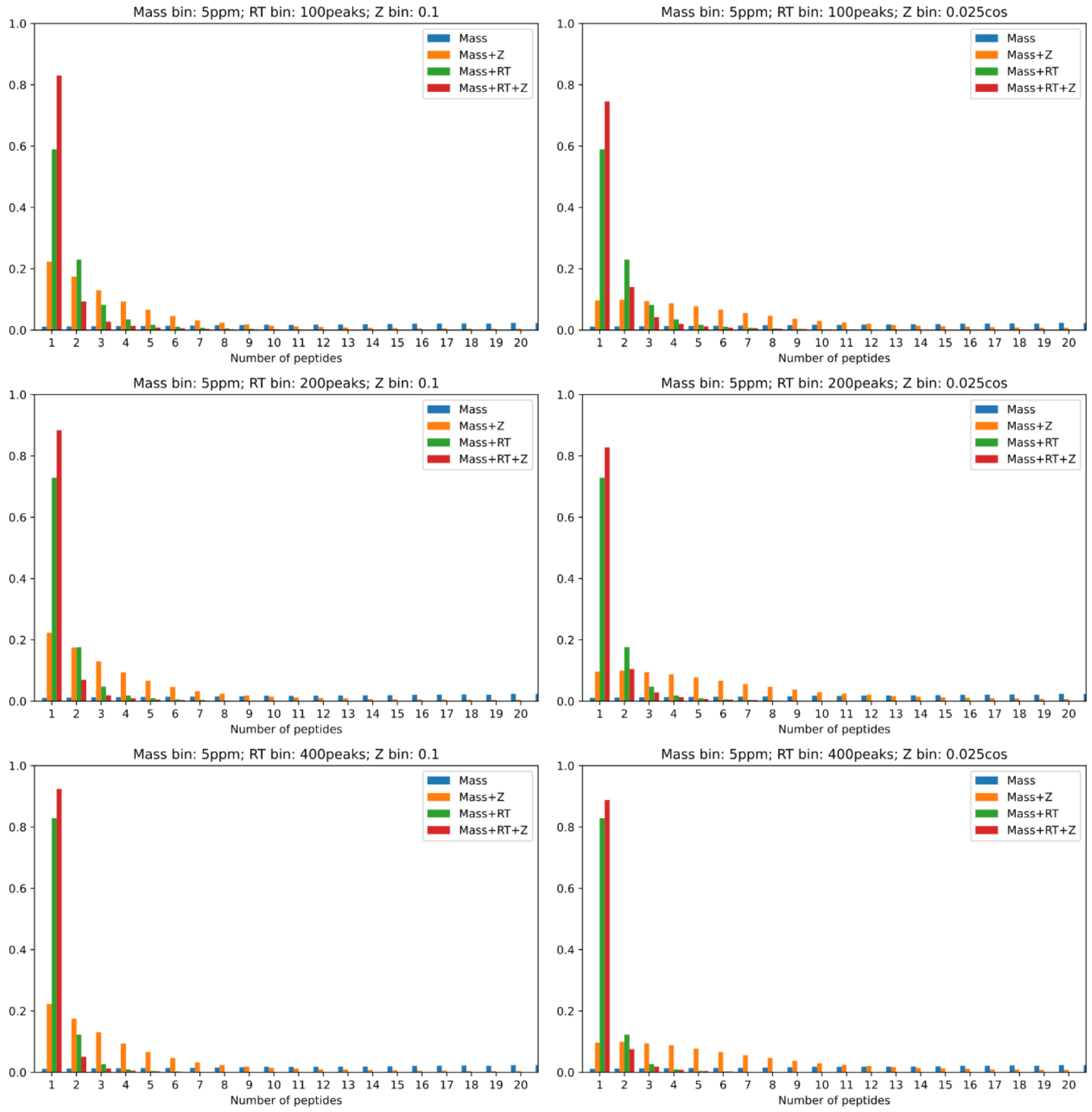


Figure C. Influence of peak capacity on the discrimination power of CSD features. Mass bin: 5 ppm; RT bin: top – 100 peaks; middle – 200 peaks; bottom – 400 peaks; Z bin: 0.1 (left column) or cosine distance of 0.025 (right column). Binning by mass (blue), mass and charge-state (orange), mass and retention time (green), and mass, retention time and charge-state (red).

(2) Virtanen, P.; Gommers, R.; Oliphant, T. E.; Haberland, M.; Reddy, T.; Cournapeau, D.; Burovski, E.; Peterson, P.; Weckesser, W.; Bright, J.; Van Der Walt, S. J.; Brett, M.; Wilson, J.; Millman, K. J.; Mayorov, N.; Nelson, A. R. J.; Jones, E.; Kern, R.; Larson, E.; Carey, C. J.; Polat, İ.; Feng, Y.; Moore, E. W.; VanderPlas, J.; Laxalde, D.; Perktold, J.; Cimrman, R.; Henriksen, I.; Quintero, E. A.; Harris, C. R.; Archibald, A. M.; Ribeiro, A. H.; Pedregosa, F.; Van Mulbregt, P.; SciPy 1.0 Contributors; Vijaykumar, A.; Bardelli, A. P.; Rothberg, A.; Hilboll, A.; Kloeckner, A.; Scopatz, A.; Lee, A.; Rokem, A.; Woods, C. N.; Fulton, C.; Masson, C.; Häggström, C.; Fitzgerald, C.; Nicholson, D. A.; Hagen, D. R.; Pasechnik, D. V.; Olivetti, E.; Martin, E.; Wieser, E.; Silva, F.; Lenders, F.; Wilhelm, F.; Young, G.; Price, G. A.; Ingold, G.-L.; Allen, G. E.; Lee, G. R.; Audren, H.; Probst, I.; Dietrich, J. P.; Silterra, J.; Webber, J. T.; Slavič, J.; Nothman, J.; Buchner, J.; Kulick, J.; Schönberger, J. L.; De Miranda Cardoso, J. V.; Reimer, J.; Harrington, J.; Rodríguez, J. L. C.; Nunez-Iglesias, J.; Kuczynski, J.; Tritz, K.; Thoma, M.; Newville, M.; Kümmerer, M.; Bolingbroke, M.; Tartre, M.; Pak, M.; Smith, N. J.; Nowaczyk, N.; Shebanov, N.; Pavlyk, O.; Brodtkorb, P. A.; Lee, P.; McGibbon, R. T.; Feldbauer, R.; Lewis, S.; Tygier, S.; Sievert, S.; Vigna, S.; Peterson, S.; More, S.; Pudlik, T.; Oshima, T.; Pingel, T. J.; Robitaille, T. P.; Spura, T.; Jones, T. R.; Cera, T.; Leslie, T.; Zito, T.; Krauss, T.; Upadhyay, U.; Halchenko, Y. O.; Vázquez-Baeza, Y. SciPy 1.0: Fundamental Algorithms for Scientific Computing in Python. *Nat. Methods* **2020**, *17* (3), 261–272. https://doi.org/10.1038/s41592-019-0686-2.

(9) Seabold, S.; Perktold, J. Statsmodels: Econometric and Statistical Modeling with Python. In *9th Python in Science Conference*; 2010.
